# Machine learning cross-platform proteomic imputation enables protein quality scoring and replication of epidemiological associations

**DOI:** 10.64898/2026.05.05.723059

**Authors:** Linke Li, Ahmed Alaa, Youxin Tan, Ilker Demirel, Samuel Friedman, Qiayi Zha, Michael Honigberg, Russell Tracy, Kent D. Taylor, Bing Yu, Christie M. Ballantyne, Rajat Deo, Ruth Dubin, Michael Y. Tsai, Gina M. Peloso, Jennifer Brody, Tom Austin, Bruce M. Psaty, Jayna Nicholas, Laura M. Raffield, Usman Tahir, Josef Coresh, Whitney Hornsby, Andrew Chan, Stephen S. Rich, Jerome I. Rotter, Peter Ganz, Robert Gerszten, Anthony Philippakis, Pradeep Natarajan, Zhi Yu

**Affiliations:** Clinical and Translational Epidemiology Unit, Massachusetts General Hospital, Boston, MA, USA; Program in Medical and Population Genetics and the Cardiovascular Disease Initiative, Broad Institute of Harvard and MIT, Cambridge, MA, USA; Department of Electrical Engineering and Computer Science, University of California, Berkeley, Berkeley, CA, USA; University of California, San Francisco, San Francisco, CA, USA; Department of Biostatistics, Harvard T.H. Chan School of Public Healt, USA h, Boston, MA, USA; Massachusetts Institute of Technology, Cambridge, MA, USA; ML4H, Broad Institute of Harvard and MIT, Cambridge, MA, USA; Heart and Vascular Institute, Mass General Brigham, Boston, MA, USA; Department of Pathology and Laboratory Medicine, Larner College of Medicine, University of Vermont, Burlington, VT, USA; The Institute for Translational Genomics and Population Sciences, Department of Pediatrics, The Lundquist Institute for Biomedical Innovation at Harbor-UCLA Medical Center, Torrance, CA, USA; Department of Epidemiology, Human Genetics and Environmental Sciences, School of Public Health, University of Texas Health Science Center at Houston, Houston, TX, USA; Department of Medicine, Baylor College of Medicine, Houston, TX, USA; Division of Cardiovascular Medicine, Perelman School of Medicine at the University of Pennsylvania, Philadelphia, PA, USA; Department of Medicine, University of Texas Southwestern Medical Center, Dallas, TX, USA; Department of Laboratory Medicine & Pathology, University of Minnesota Medical School, Minneapolis, MN, USA; Department of Biostatistics, Boston University School of Public Health, Boston, MA, USA; Cardiovascular Health Research Unit, Department of Medicine & Deparment of Epidemiology, University of Washington, Seattle, WA, USA; Department of Genetics, University of North Carolina at Chapel Hill, Chapel Hill, NC, USA; Division of Cardiovascular Medicine, Beth Israel Deaconess Medical Center, Boston, MA, USA; Department of Medicine, NYU Grossman School of Medicine, New York, NY, USA; Department of Medicine, Harvard Medical School, Boston, MA, USA; Department of Genome Sciences, University of Virginia, Charlottesville, VA, USA; Division of Cardiology, Zuckerberg San Francisco General Hospital and Department of Medicine, University of California San Francisco, San Francisco, CA, USA; Google Ventures, Cambridge, MA, USA

## Abstract

High-throughput affinity-based proteomics has advanced biomedical research, yet fundamental, persistent discordance between mainstream platforms (SomaScan and Olink) routinely undermines the replication of findings. This platform-driven non-replication complicates downstream biological validation and biomarker prioritization. Here, we develop a machine learning-based framework for cross-platform protein value imputation to navigate this translational bottleneck. Using paired proteomic data measured by both SomaScan and Olink from 5,325 participants of the Multi-Ethnic Study of Atherosclerosis, we developed models to impute cross-platform measurements and applied them to two independent and demographically distinct cohorts (Cardiovascular Health Study [N=3,171] and UK Biobank [UKB; N=41,405]) to evaluate downstream replication. Our bidirectional model 1) established an imputation performance-based protein fidelity index, validated against gold-standard / orthogonal measurements (ELISA and Simoa/Quanterix in Atherosclerosis Risk in Communities study [ARIC, N=101] and Nurses’ Health Study [NHS, N=54], and published mass spectrometry data), 2) enabled imputation of platform-exclusive protein measurements, and 3) facilitated calibration of overlapping proteins. We demonstrate the utility of this framework through three applications: 1) fidelity-informed analyses enhanced the replication of biomarker discovery, 2) recovery of SomaScan signals that were previously inaccessible in UKB’s original Olink measurements, and 3) improved replication performance for overlapping proteins. Our study offers a translational roadmap that helps researchers identify more confident cross-platform measurements to support replication efforts, identify potential assays warranting future optimization, and distinguish proteins where biological signal likely dominates platform-specific measurement.

## INTRODUCTION

The identification and quantification of circulating plasma proteins have become central to advancing biomarker discovery, particularly in large-scale epidemiological studies^1–3^. High-throughput affinity-based proteomics platforms, which measure thousands of circulating proteins, have greatly accelerated biomedical research in the identification of biomarkers and potential drug targets^3,4^. Two widely adopted high-throughput measuring services are SomaScan (SomaLogic, Inc) and Olink (Thermo Fisher Scientific, Inc). SomaScan utilizes a DNA-based approach of slow off-rate modified aptamers (SOMAmers)^5^, while Olink platform adopts an antibody-based approach known as proximity extension assay (PEA)^6^.

This rapid expansion in proteome measurements has advanced population and medical research at an unprecedented scale, but it has also introduced significant challenges. While affinity-based platforms have excelled beyond traditional methods like enzyme-linked immunosorbent assay (ELISA) in scale and efficiency, this approach comes with an inherent trade-off. Their reliance on secondary probe-binding signals, rather than direct molecular detection of peptide mass, makes them highly susceptible to off-target interactions driven by protein-altering variants, platform-specific artifacts, and complex structural conformations^7,8^. Consequently, correlations between measurements are frequently modest to low. For example, median or mean cross-platform correlations of 0.29 (China Kadoorie) to 0.46 have been reported across large cohort comparisons, with only 15–19% of proteins reaching strong concordance (r ≥0.75–0.8): not only between SomaScan and Olink, but also when compared against gold-standard measures^4,9,10^. This leads to the non-replication of associations in studies using different platforms^4,9–11^. Another contributor to this lack of replication is the limited number of shared proteins between platforms and panels^12^. For instance, between SomaScan v4.0 (∼5000 proteins) and Olink Explore 3072 (∼3000 proteins), only ∼1,800 appear on both panels. Naturally, the thousands of proteins unique to each platform are inaccessible for orthogonal validation. As a result, researchers are often left unable to distinguish whether the non-replication arose from false discovery or simply discrepancies between the measuring platforms.

This necessitates a systematic and translational framework to evaluate protein measurement reliability and bridge the gap in coverage between technologies. To address these challenges, we leveraged machine learning approaches and paired proteomic measurements of the same samples from the Multi-ethnic Study of Atherosclerosis (MESA)^13^. By training on these concurrent proteomics data, our models captured the relationships between the two technologies in a data-driven fashion. This framework allowed us to 1) conduct cross-platform imputation and 2) establish a protein tiered reliability system based on cross-platform fidelity that assists in identifying measurements with higher confidence for reproducible translational research. To validate the translational utility of our framework, we improved replication of published epidemiological associations across independent cohorts with disjoint proteomic and demographic profiles: UK Biobank (UKB)^3,14,15^ (Olink-only) and Cardiovascular Health Study (CHS)^16,17^ (SomaScan-only), demonstrating that fidelity-informed prioritization can partially mitigate the reproducibility bottleneck induced by platform-driven discordance (**Figure 1**).

**Figure 1.**
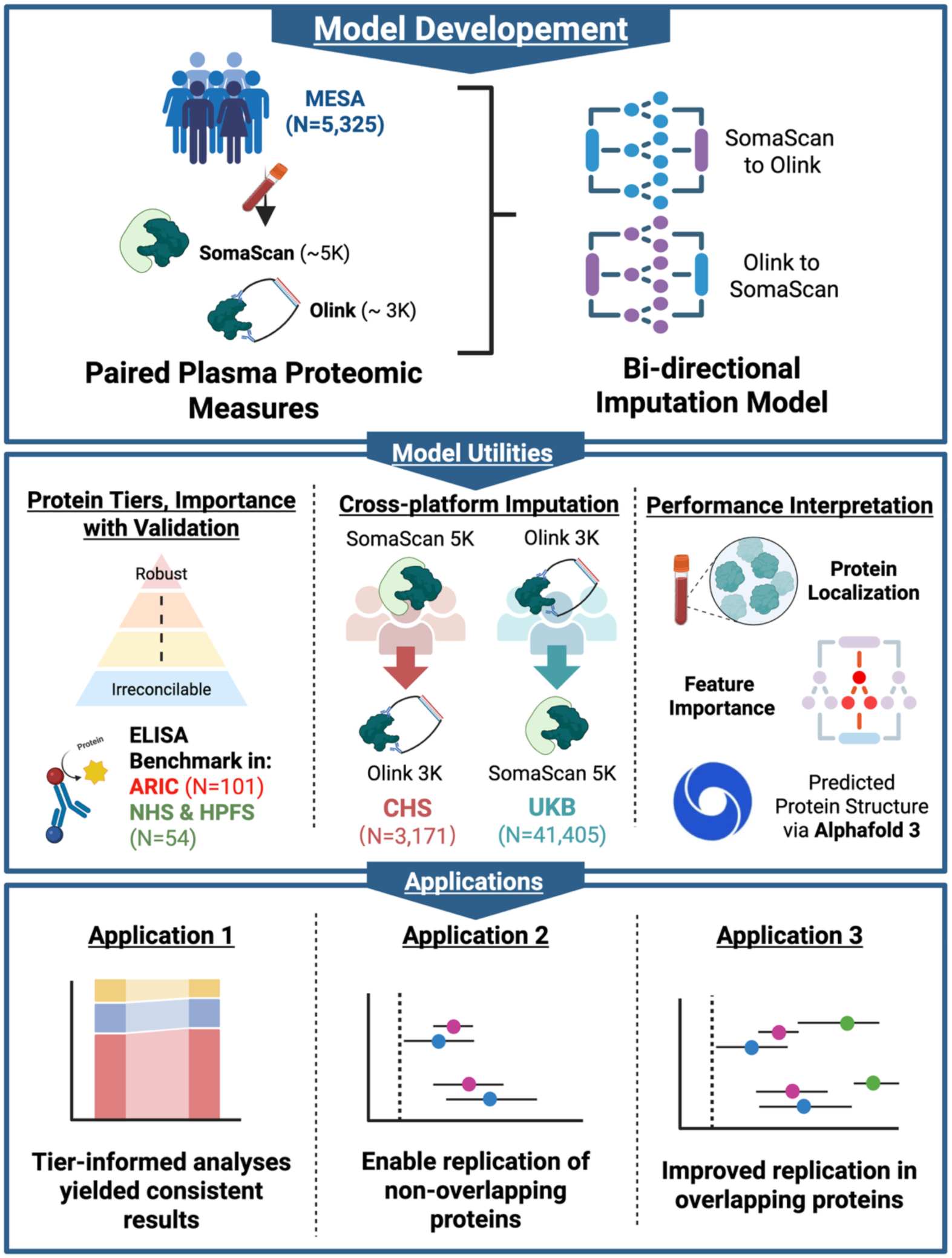
Scheme of Study Design. We developed bidirectional imputation models using paired SomaScan and Olink measurements from <u>MESA</u>. The models would impute the measurements from another platform in single-platform cohorts using measured values (<u>CHS</u>, <u>UKB</u>). We also proposed a tier system for protein measurement qualities and model recoverability and supported the various protein performances using empirical validations such as protein class (secreted / plasma, membrane, intracellular), ELISA, feature importance, and predicted protein structure by Alphafold 3. The validity of the imputed values and the tier system was further tested in 3 real-world applications: 1) tier-informed analyses yielded highly replicable and consistent results, 2) imputation enables validation of non-overlapping proteins in single-platform cohorts, and 3) imputed values improved replication of signals compared to using the naive measures. MESA: Multi-Ethnic Study of Atherosclerosis; ELISA: enzyme-linked immunosorbent assay; ARIC: Atherosclerosis Risk in Communities Study; NHS & HPFS: Nurses’ Health Study and Health Professionals Follow-Up Study; CHS: Cardiovascular Health Study; UKB: UK Biobank. This figure was created with BioRender.com

## RESULTS

### Study Population and Protein Processing

To establish a robust framework for cross-platform proteomic imputation and quality scoring, we curated and harmonized data from three large-scale cohorts: the Multi-Ethnic Study of Atherosclerosis (MESA), the UK Biobank (UKB), and the Cardiovascular Health Study (CHS) (**Supplementary Table 1**,**Supplementary Figure 1**). The MESA cohort is our dataset for model development, providing a resource of 5,325 participants with concurrent proteomic measurements from both SomaScan (v4.1, ∼7000 proteins) and Olink (Explore 3072, ∼3000 proteins) platforms. We randomly divided MESA into a training set (N=3,993) for model development and a held-out internal testing set (N=1,332). After quality control, UKB (N=41,405; Olink Explore 3072, ∼ 3000 proteins) and CHS (N=3,171; SomaScan v4.0, ∼5000 proteins) were used as external application cohorts (**Supplementary Figure 1**).

**Supplementary Table 1** displays the characteristics of the three demographically diverse cohorts. The three study populations span different life stages, with mean ages of 57.3 years (standard deviation [SD]=8.19) in UKB and 62.2 years (SD: 10.3) in MESA, compared to a significantly older CHS cohort (mean age: 74.4 [4.9]). The majority of UKB and CHS participants (UKB: 93.1%, CHS: 83.5%) are white, and MESA participants have higher diversity in self-reported race and ethnicity, including representation from Black (25.3%) and Hispanic/Latino (23.0%) participants. No significant difference was observed for health status in the MESA train and test sets (**Supplementary Table 2**).

We standardized protein measurements across the three cohorts to define the platform-specific and shared feature spaces (**Supplementary Figure 1**). SomaScan aptamers were harmonized to the v4.0 panel using SomaDataIO (R package, version 6.3.0)^18^ lift-over to ensure consistency between MESA and CHS, resulting in a final set of 4,972 SomaScan proteins. For Olink, 2,871 proteins remained after filtering for missingness. The intersection of these two platforms yielded 1,737 overlapping proteins (35% of SomaScan proteins and 60% of Olink proteins).

### Cross-platform Correlation, Predictive Modeling, and Performance by Protein Localization

Consistent with previous reports^4,9,10,12,19–21^, baseline concordance between raw SomaScan and Olink measurements was limited. Among the 1,737 overlapping proteins in MESA, the mean Pearson correlation (r) was 0.36, with the majority of proteins (N=988, 57%) exhibiting below moderate correlation (r<0.4) and only 334 (19%) proteins showing strong agreement (r>0.7). This baseline discordance varied across the proteome, ranging from null (r=0.0001 for CRISP3 [cysteine-rich secretory protein 3]) to highly correlated (r=0.95 for SIGLEC5 [sialic acid-binding Ig-like lectin 5]). We also report Spearman’s correlation (rho) as a complementary, rank-based metric alongside Pearson’s r (**Supplementary Table 3**).

To bridge this gap, we used histogram-based gradient boosting (HGB) regressors trained on paired MESA data to develop a bidirectional imputation framework. This approach enabled the bidirectional imputation of measurements of Olink using SomaScan (SO) and of SomaScan using Olink (OS) in single-platform cohorts such as UKB and CHS, thereby avoiding the forced integration of potential mismeasured values.

Our imputation yielded systematic improvement in concordance for both prediction directions, reflected by a widespread shift of proteins above the identity line relative to baseline (**Figure 2**, **Panel A & B; Supplementary Figure 2A**). While our models impute the entire proteome, comparison between post-imputation and baseline correlation was restricted to the overlapping proteins. We define post-imputation correlation as Pearson correlation between machine-learning imputed platform measurement and their measured counterparts (e.g., between Olink-imputed SomaScan and measured Olink), and gains as the improvement in correlation after imputation. The mean cross-platform concordance between imputed values and their measured counterparts increased from 0.36 to 0.48 for both directions. The number of well-correlated proteins (r>0.7) doubled from ∼19% to ∼36–38% (N_SO_=656, N_OS_=633), and the proportion of poorly correlated proteins (r<0.4) was reduced from ∼57% to 46% (N_SO_ = 801, N_OS_=790) (**Figure 2**, **Panel A & B**).

**Figure 2.**
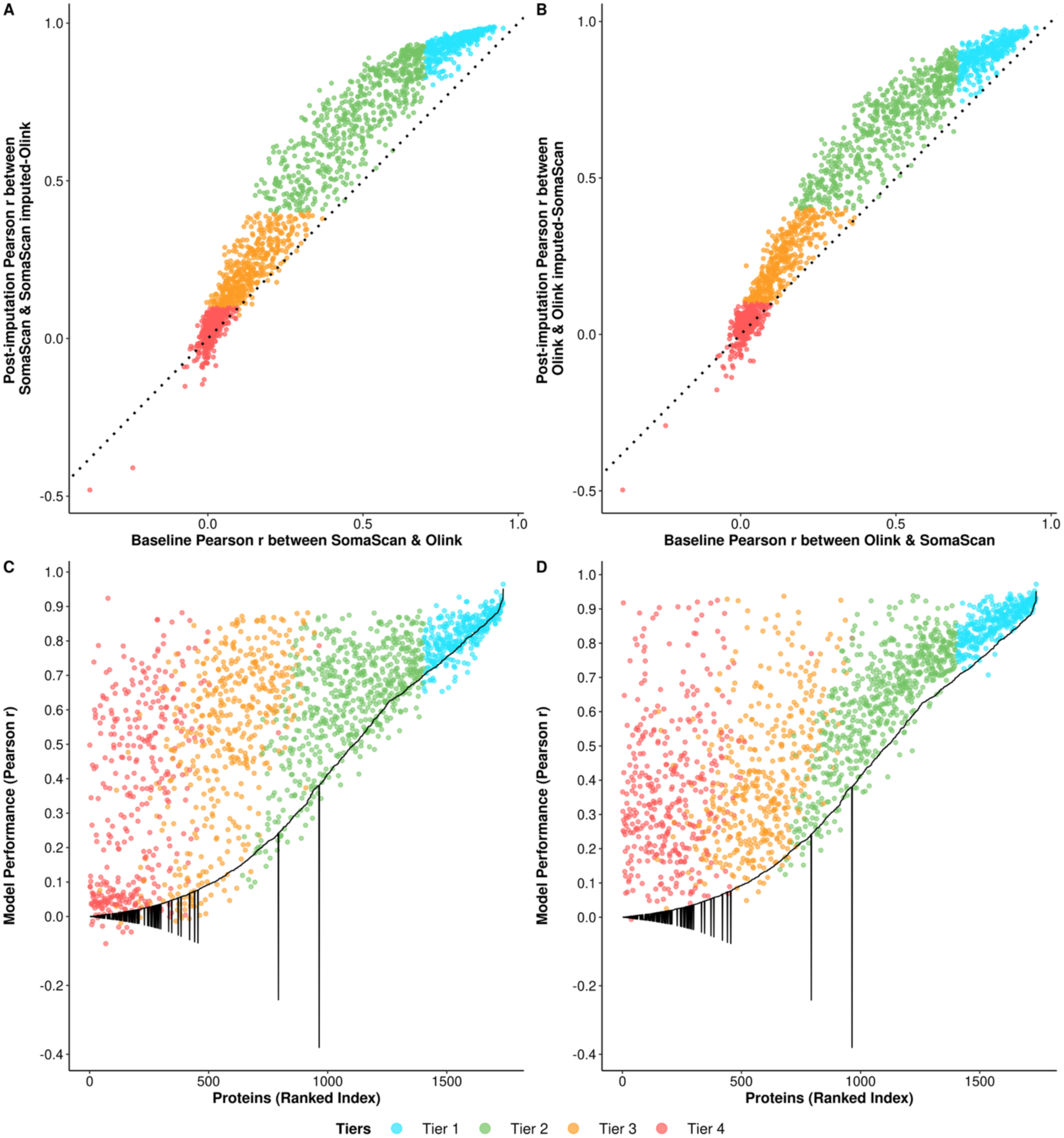
Comparison of baseline and post-imputation protein correlations in the MESA cohort of 1,737 overlapping proteins. The MESA cohorts were separated into 75% training (N = 3,993) and 25% hold-out internal validation (N=1,332) sets. **Scatter plots A. SomaScan to Olink direction, B. Olink to SomaScan direction -** Scatter plots comparing baseline correlation (Pearson correlation of paired measured SomaScan and measured Olink values) and post-imputation correlation (Pearson correlation between measured SomaScan and SomaScan imputed-Olink in SomaScan to Olink direction, and between measured Olink and Olink-imputed SomaScan in Olink to SomaScan direction). Each point represents an individual protein, colored by its tier membership. The black dotted identity line (x=y) indicates no change in correlation; points above this line represent proteins showing improved correlation post imputation, and vice versa. **Scatter plot C. SomaScan to Olink direction, D. Olink to SomaScan direction -** Model predictive performance in relation to baseline correlation. The predictive power of the model is defined as the Pearson correlation between model-imputed value and measured value in the MESA held-out test set. Proteins are ranked by their baseline Pearson correlation squared along the x-axis to illustrate the magnitude of baseline correlation, including negative correlations. The black line represents baseline correlation. Each point, colored by proteins’ respective tier, denotes the model performance of that protein in relation to its baseline correlation

For most proteins, imputation success (i.e., gain in correlation) relates to baseline cross-platform correlation. Proteins that were initially well-correlated between SomaScan and Olink were more amenable to accurate imputation. However, there were a number of exceptions where our models recovered signal despite poor baseline agreement. For instance, IVD (isovaleryl-CoA dehydrogenase, SO direction), which exhibited a low baseline correlation (r=0.21) between platforms, achieved a substantially higher correlation of r=0.61 after imputation (gain=0.4). We also observed significant increases in correlation in receptor proteins in the OS direction (e.g., LEPR [leptin receptor], 0.40→0.75). Cross-platform post-imputation correlation remains highly similar for the same protein when comparing two directions (**Supplementary Figure 2C**).

We further evaluated model performance using the MESA held-out test set for all proteins by comparing imputed values against actual measurements (e.g., Olink-imputed SomaScan vs. actual measured SomaScan) (**Figure 2**, **Panel C & D**). The overall predictive performance was broadly effective for both directions (mean r=0.57), with 684 (39%) of 1,737 proteins exhibiting high performance, and there were 384 (22%) proteins with moderate to low performance (r<0.4, **Supplementary Figure 2B**); in general, overlapping proteins have higher predictive performances than non-overlapping proteins.

In addition, we examined whether protein localization influenced cross-platform concordance both at baseline and post-imputation. Proteins secreted into the blood or plasma exhibited higher baseline correlations compared to membrane-bound or intracellular proteins (**Figure 3**). While all protein classes benefited from our framework, secreted and plasma proteins showed the most substantial improvements, frequently achieving moderate-to-strong imputed correlations. This suggests that the extracellular proteome may be more consistently captured across affinity-based technologies. Among the few proteins with a gain <-0.05 (N_SO_=20, N_OS_=18), membrane proteins were significantly enriched in the SO direction (**Supplementary Figure 3**).

**Figure 3.**
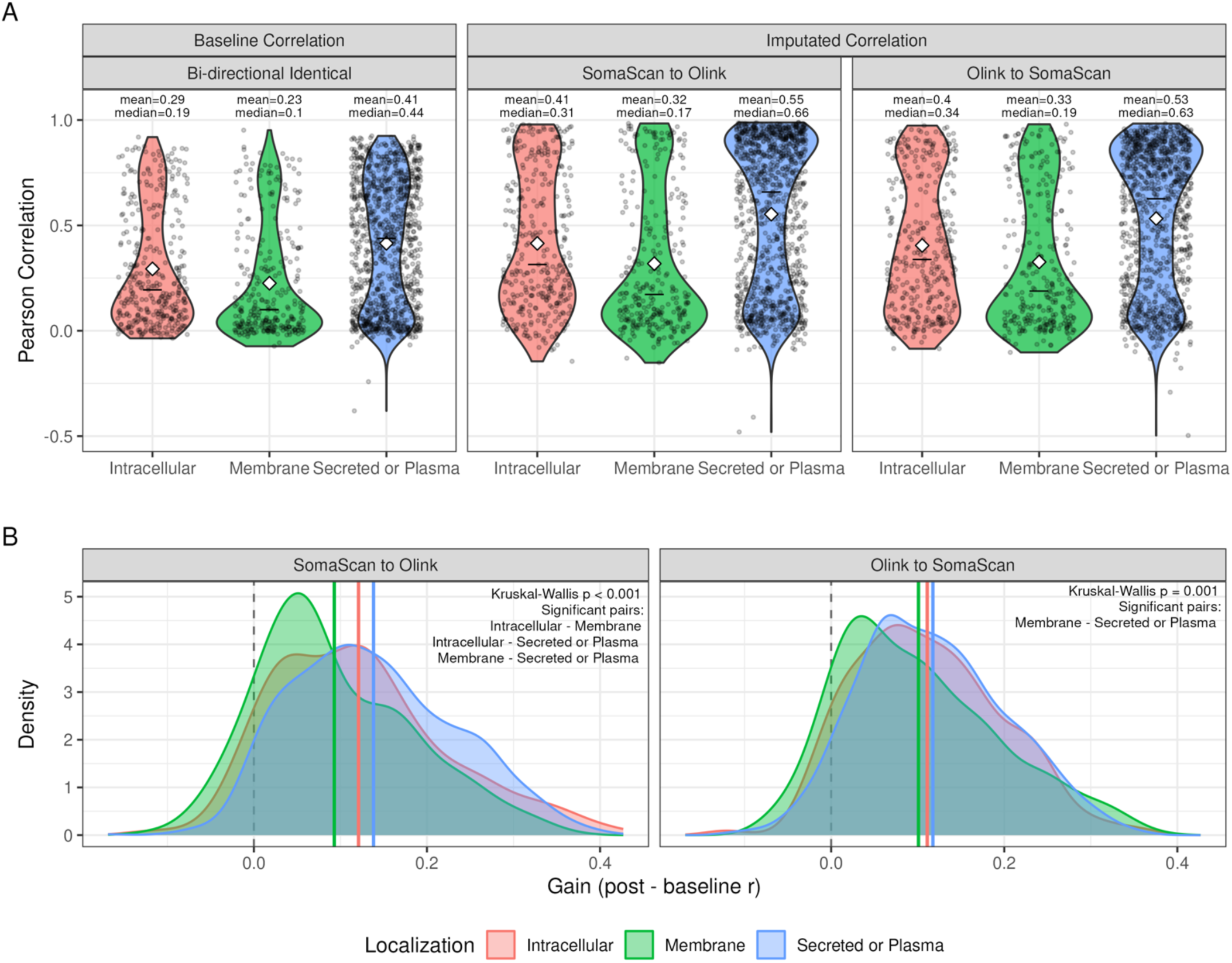
Correlation between two measuring platforms by protein localization. **Panel A. Distribution of the bidirectional baseline and post-imputation correlations stratified by protein localization.** Proteins are classified as intracellular, membrane, secreted or plasma, as this study utilized plasma proteomic datasets based on Human Protein Atlas’s protein class information. The Uniprot subcellular location information was utilized for the 5 proteins (LILRA3, MENT, LEG1, DDX58, KIR2DL2) missing location information in HPA. Violin plots display the distribution of correlations between the two platforms. Mean and median of the cross-platform correlation is annotated. Baseline correlation is identical across both directions. Imputed correlation was calculated using SomaScan and SomaScan-imputed Olink, or Olink and Olink-imputed SomaScan. Plasma or proteins secreted into the plasma benefited the most from imputation as the model reconstructed their measurement value with information from their biological ‘neighbors”. **Panel B. Density plots of the cross-platform correlations**. Within each imputation direction, differences in gain (post-imputation corr - baseline corr) across protein localization classes were assessed by Kruskal-Wallis test, with pairwise comparisons via Dunn’s test. The red, green and blue vertical lines represent class means.

### Model Calibration, Variance Preservation, and Uncertainty

Beyond Pearson correlation, we assessed calibration slope and intercept, root-mean-square error (RMSE), and variance preservation (the ratio of standard-deviation of imputed and measured values) across tiers (**Supplementary Table 4**). Calibration slopes and SD ratios declined monotonically from tier 1 to tier 4 in both directions, while RMSE increased, indicating that higher-tier proteins are not only more correlated but also better calibrated. We additionally quantified prediction uncertainty using split conformal prediction^22^, reporting mean, median, minimum, and maximum interval half-widths by tier (**Supplementary Table 5**); uncertainty intervals widened consistently from tier 1 to tier 4, reinforcing the ordinal validity of the tiering system.

### Benchmarking Against Alternative Modeling Approaches

To evaluate whether our histogram-based gradient boosting (HGB) approach is empirically justified relative to simpler, more commonly used epidemiological modeling strategies, we benchmarked HGB against elastic net (a standard regularized linear approach commonly used in epidemiological studies) and LightGBM (LGB, an independently implemented boosting algorithm). We quantified (1) pairwise tier-membership concordance (Cohen’s κ), (2) each method’s tier assignment’s association with an independent mass-spectrometry validation dataset (η^2^; Sissala et al., 2025^8^), and (3) Steiger’s Z tests comparing each method’s correlation with this ground truth. HGB and LightGBM showed near-identical tier concordance (κ=0.96-0.98) and comparable performance against the mass-spectrometry benchmark (η^2^∼0.145 for SO direction, ∼0.175 for OS direction), while elastic net trailed modestly (κ=0.83-0.88; η²=0.122 for SO & 0.172 for OS). In the SomaScan-to-Olink direction, HGB and LightGBM significantly outperformed elastic net (Z=7.10 & 6.99, respectively; P <10⁻¹¹ for both). These results supported using HGB as the main method (**Supplementary Table 6**).

### Tiered Protein Reliability System

Based on the different imputation performances, we categorized the 1,737 overlapping proteins into a tiered-reliability system (**Supplementary Table 7**) for each training direction independently, based on conventional thresholds to interpret correlation coefficients (e.g., >0.7 is strong, <0.1 is weak or negligible)^23^. We define tier 1 as the most ‘robust’ group, with both baseline correlation and imputed concordance being r≥0.7. This high-trust subset should exhibit exceptional cross-platform portability and measurement quality, allowing researchers to replicate primary protein-outcome findings in independent cohorts using proteomics measured by either platform reliably, regardless of the use of our predictive models. In contrast, tier 4 is the ‘irreconcilable’ group, defined by having both baseline and imputed concordance r<0.1. These proteins exhibit persistent discordance and cannot be recovered by our framework, which could be caused by platforms’ binding affinities (e.g., PILRA)^1,21^ or potential off-target binding.

After isolating tier 1 (robust) and tier 4 (irreconcilable) proteins, the remaining proteins were further categorized into two groups (tier 2: imputed r≥ 0.4, tier 3: imputed r<0.4). Tier 2 is the ‘model recoverable’ group, representing proteins with modest to low baseline correlation that gained and achieved acceptable (r>0.4) correlation post-imputation. This group expanded the ‘usable’ portion of the proteome, allowing its use in cross-platform validation. The protein IVD mentioned previously is a tier 2 protein. Lastly, the remaining proteins were categorized as tier 3, the ‘ambivalent’ group: while our imputed value offered a slight increase in concordance, the concordance remained relatively low. It is also advised that associations identified with tier 3 proteins be assessed carefully and skeptically, as their cross-platform concordance is insufficient for reproducibility.

In each direction, there are 19% tier 1, 35% tier 2, 25% tier 3, and 20% tier 4 proteins, respectively. Bidirectionally, all of tier 1 (N=334), and the vast majority (>90%) of tier 2 and tier 4 proteins are identical, reflecting the robustness of the membership of the tiers: tier 1 and 2 proteins are consistently reliable, and tier 4 proteins are consistently discordant. The most variability of protein tier memberships was among tier 3 proteins. Proteins that shifted their tier assignment were more likely to be non-secreted, non-plasma proteins, relative to their background frequency among the overlapping proteins (**Supplementary Figure 4**). At the same time, no significant difference in post-imputation correlation between demographic subgroups was observed (**Supplementary Figure 5**). We further tested the sensitivity of tier assignment to the correlation thresholds used to define tiers, applying alternatives other than conventional cutoffs (0.1, 0.4, 0.7) (**Figure 4, Panel A**) and the uncertainty of tier assignment with bootstrap (1,000 resamples; **Figure 4, Panel B & C**) in the MESA test set.

**Figure 4.**
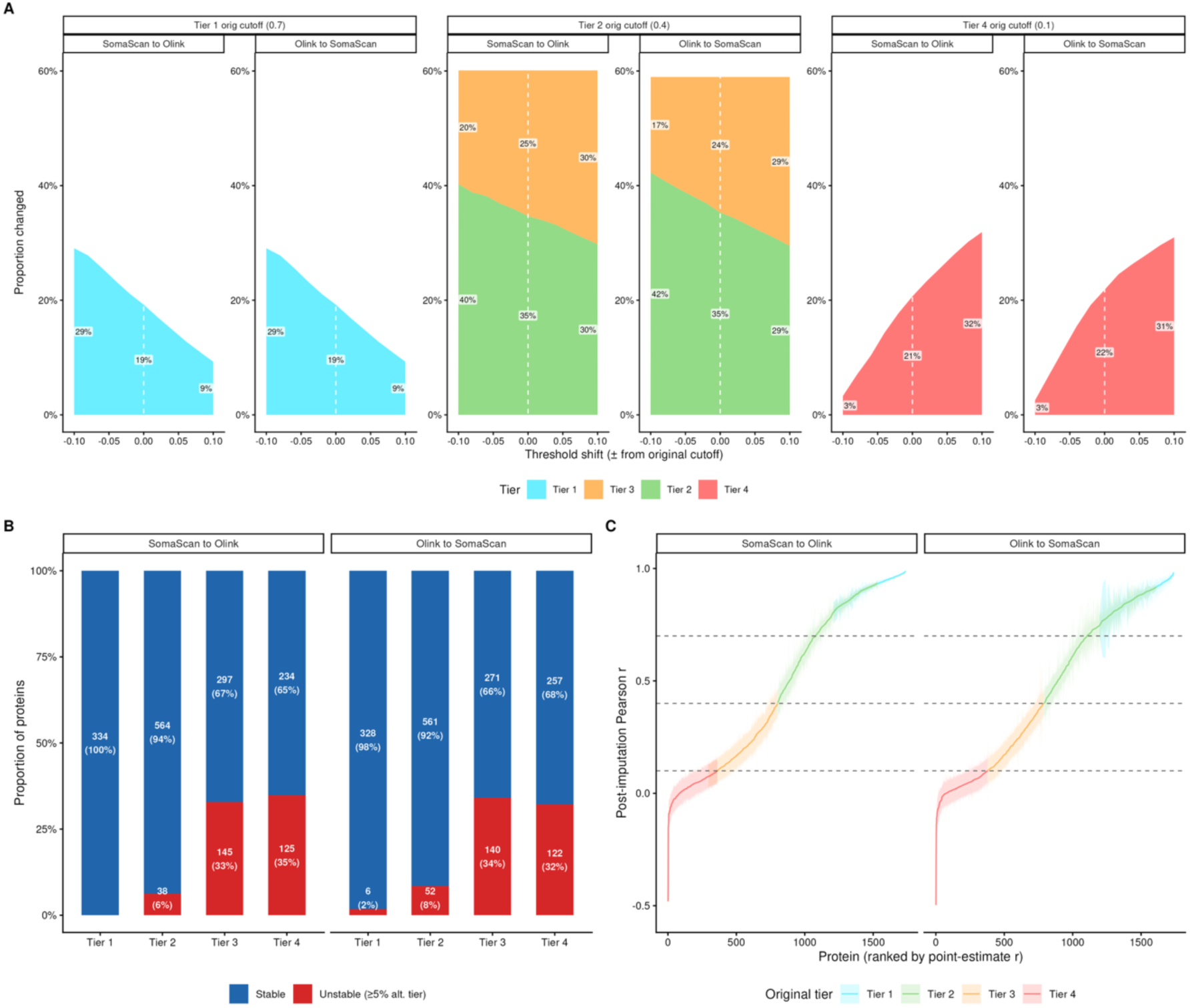
Tier assignment sensitivity by shifting threshold & bootstrapping and protein correlation bootstrapping 95% confidence intervals. **Panel A. Change in proportion of proteins within each tier when threshold changes from the conventional correlation strength definition (e.g., 0.7 =strong; 0.4=moderate; 0.1=weak)**. The threshold changes with a -0.1 to 0.1 threshold with an interval of 0.02. At each alternative threshold, proportion of proteins within each tier is re-calculated. **Panel B. proportion of proteins that has either stable or unstable tier assignments in 1,000 bootstrap resamples**. For each bootstrap, the post-imputation was recalculated in each sample and thus a bootstrap tier is re-assigned. Among the 1,000 resamples/bootstraps, if a protein was assigned another tier larger than or equal to 5% (alpha=0.05) of the resamples, it would be categorized as an ‘unstable’ protein. **Panel C. 95% confidence interval of post-imputation correlation using bootstrap**. 95% confidence intervals were constructed from the 2.5th and 97.5th percentiles of the resulting bootstrap distribution.

We observed clear increases in correlation across proteins in all tiers except for tier 4. Tier 1 proteins exhibited a ceiling effect, approaching a correlation of 0.93 for SO and 0.90 for OS. The most substantial gains were observed in tier 2 and some tier 3 proteins; tier 2 proteins, on average, gained 0.21 in Pearson correlation in SO direction and 0.19 in OS direction, while tier 3 proteins gained 0.08 in SO direction and 0.10 in OS direction (**Figure 2**, **Panel A & B**).

### Orthogonal Validation of the Tiered Reliability System

To further evaluate the robustness of our tiered reliability system, we performed three complementary analyses using established or independent measured gold standards (enzyme-linked immunosorbent assay [ELISA] and Quanterix, mass spectrometry [MS]) as benchmarks.

In line with our definition, all three tier 1 proteins measured in Nurses’ Health Study (NHS) and Health Professionals Follow-up study (HPFS) (N=54, **Supplementary Figure 6**)^24^ showed moderate-to-strong correlation with ELISA measurements (CHI3L1: 0.67; GDF15: 0.85; SELE: 0.83), and the single tier 2 protein available for validation, IL1RN, showed a comparable correlation (Spearman rho=0.69). On the other hand, GFAP (glial fibrillary acidic protein) and NfL (neurofilament light chain) measured by SomaScan were validated to have low correlation with Alamar (NULISA) or Simoa/Quanterix (Ibanez et al., r∼0.2, N=43^25^; and in Atherosclerosis Risk in Communities [ARIC], N=101, **Supplementary Figure 7**, **Supplementary Table 8**), while both SomaScan and Olink measures of MAPT (microtubule-associated protein tau) are not correlated with Simoa. These validated concordance and discrepancies support the quality of tier 1 proteins and also imply that epidemiological or clinical associations identified with tier 4 proteins could be spurious and warrant caution in findings based on proteins in this group.

We further compared our tier assignment to another independently collected mass spectrometry dataset (Sissala et al. 2025^8^) as an external orthogonal benchmark (N=821 overlapping proteins), finding that higher-tiered proteins showed higher correlation with these experimental measurements (**Supplementary Figure 8**).

### Protein Importance

Beyond assessing cross-platform reliability itself, we wanted to examine whether a protein’s reliability tier relates to how heavily it is already used or trusted in biomedical research, since proteins with high scientific or clinical attention are often assumed, implicitly, to be well-measured by different services. Therefore, we annotated each protein with its ‘importance’ using two main criteria: 1) clinical utility as assessed using the Human Protein Atlas (HPA)^26,27^, classifying proteins as “FDA-approved” or “potential” drug targets, and 2) scientific attention as assessed by the MalaCards and DISEASE^28–30^ databases to aggregate the total number of publications associated with each gene symbol.

**Supplementary Figure 9** visualizes protein importance stratified by our tiered reliability system. Gain in correlation was not restricted to a subset of well-studied markers, i.e. proteins with high publication counts, shown as darker bins in **panel A**, were distributed across all four tiers rather than concentrated in tier 1 only. In contrast, other frequently studied proteins remained poorly measured despite their prominence. TNF (tumor necrosis factor, **panel B**), an FDA-approved drug target with substantial publication attention, falls in tier 4 (OS direction) or tier 3 (SO direction), indicating persistent cross-platform discordance despite its central role in epidemiological and clinical research.

This pattern of high-attention proteins appearing across the full range of reliability tiers rather than clustering in tier 1 or 2 suggested that scientific or clinical prominence does not guarantee cross-platform measurement reliability, and raises a cautionary flag for interpreting study results involving these proteins. As there is no single, universally accepted metric for a protein’s scientific or clinical importance, this analysis is intended as a motivating illustration.

#### Application 1 - Tier 1 Protein for Robust and Replicable Associations

To validate our tiered reliability system, application 1 demonstrates how prioritizing tier 1 would enhance the replicability of findings. We utilized elastic net regression to model body mass index (BMI) against the 1,737 overlapping or 334 tier 1 proteins across the UKB (Olink only) and CHS (SomaScan only) cohorts, adjusting for age, sex, and race as unpenalized covariates. After removing participants with missing BMI, the final analytic dataset of UKB and CHS contained 41,185 and 3,153 participants, respectively. The mean BMI of UKB participants is 27.4 (4.8), and 26.7 (4.5) in CHS. The CHS cohort represents a geriatric US population, distinct from the middle-aged UK Biobank population (**Supplementary Table 9**). Because the full-panel and tier1-only models differ substantially in non-zero predictor counts, we additionally report adjusted R^2^ which accounts for the number of predictors alongside raw R^2^.

A naive cross-platform analysis using the full 1,737-protein overlapping panel revealed inconsistencies in biomarker prioritization (**Supplementary Table 10**). Specifically, the rank-ordering of proteins with the highest predictive power for BMI showed disagreement between the UKB and CHS cohorts, with only LEP exhibiting consistent importance across both platforms (**Figure 5**). In addition, CHS analysis was heavily penalized (raw R^2^: 0.723, adjusted R^2^: 0.381) due to the high number of predictors and smaller sample size compared to UKB. In contrast, prioritizing the 334 tier 1 (robust) proteins resolved the discrepancy. This subset maintained comparable model performance across both cohorts (raw R^2^ & adjusted R^2^), despite using far fewer predictors (334 vs. 1,737, **Figure 5**). Moreover, this tier-informed analysis selected more consistent proteins across platforms and cohorts, with or without our imputation models (**Figure 5, right panel**). Among the top five proteins, LEP and FABP4 (fatty acid binding protein 4) ranked in the top five in all four tier-1 analyses, and NCAN (Neurocan) and RNASE1 (ribonuclease A family member 1, pancreatic) in three of four, demonstrating highly similar relative importance (**Figure 5**, **Supplementary Table 10**). These results emphasized that selecting or prioritizing tier 1 proteins significantly enhances the replicability of cross-platform epidemiological studies.

**Figure 5.**
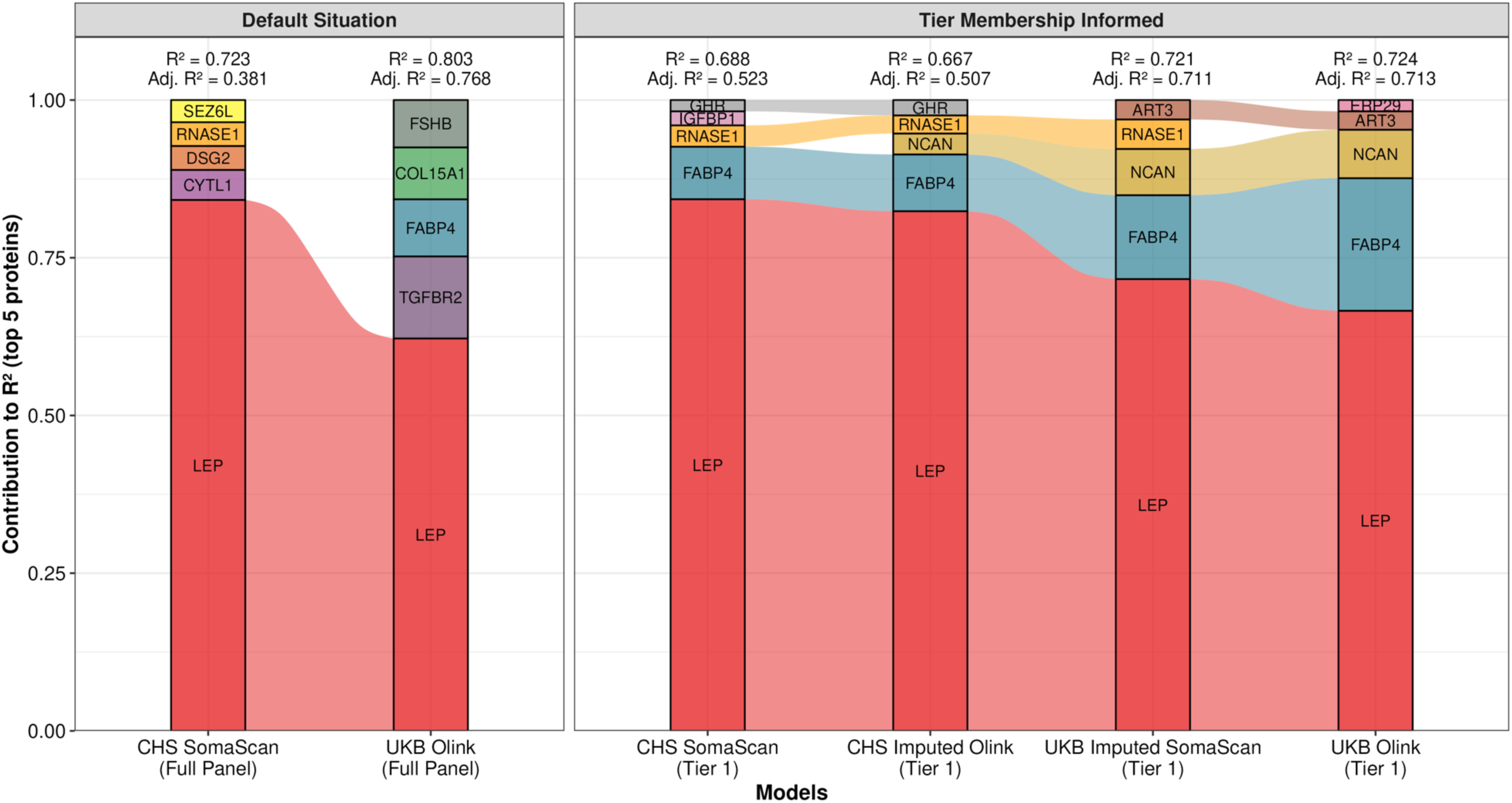
Application 1 - Feature importance of most important proteins for BMI prediction in UKB and CHS. The size of colored areas is permutation feature importance of each protein, and it is calculated using the Permutation-based approach. For each model, the raw R^2^ (variance in BMI explained by the model) and adjusted R^2^ (adjusted for the number of non-zero predictors in the test set) is reported as measures of model performance. For each protein, we randomly permuted its values in the test data, generated new predictions using the refitted model, and calculated the resulting model predictive performance (R^2^) change as permutation based feature importance. A larger change indicates more influential proteins. The ‘default situation’ refers to situations using all 1,737 overlapping proteins, while analyses in ‘tier membership informed’ situations were restricted to using only 334 tier 1 (most robust) proteins.

#### Application 2 - Enabling Replication of Non-overlapping Proteins (Walker et al., *Nature Aging* 2021)

To evaluate the validity of imputed values, we used imputed ∼5000 UKB “SomaScan” proteins based on its original Olink values to conduct epidemiological replication of previously reported protein-dementia associations by Walker et al.^31^. We selected this study because the top associated proteins were SomaScan exclusive (e.g., SVEP1 [sushi, von willebrand factor type A, epidermal growth factor, and pentraxin domain containing 1]). In brief, proteome-wide association scan was conducted using Cox proportional hazard model adjusting for covariates (**Methods**) identical to Walker et al.^31^. The analytical cohort for this use case was also derived from the original UKB population (**Supplementary Table 1**, N=41,405), with specific inclusion criteria of unrelated participants with available whole exome sequencing for apolipoprotein-E epsilon 4 (APOE-ε4) allele calling (**Supplementary Table 11**, N=36,854). The incidence rate of dementia per 100,000 person-years in UKB was 76.8, with a median follow-up time of 13.5 years (interquartile range [IQR]: 12.8-14.3 years).

SVEP1 is the leading discovery in the study, the only protein being highlighted in the abstract of the paper. It was strongly associated with incident dementia in discovery using ARIC latelife cohort, validated internally in ARIC midlife cohort, and externally in AGES-Reykjavik cohort^31^. In the ARIC midlife analysis, which has the most similar mean age compared to UKB, Walker et al. demonstrated that one SD increase in the two baseline SVEP1 measurements from different aptamers on the log2 scale was associated with a 1.42-fold (95% confidence interval [CI]: 1.20-1.67) and 1.49-fold (95% CI: 1.28-1.74) increase in hazard of incident dementia. In UKB, the hazard ratio of one SD increase in baseline imputed SVEP1 value on the log2 scale and incident dementia is 1.47 (95% CI: 1.17-1.83, P=7.30 × 10^-4^) and 1.33 (95% CI: 1.07-1.66, P= 0.01), demonstrating nearly identical associations (**Figure 6A, Supplementary Table 12**).

**Figure 6.**
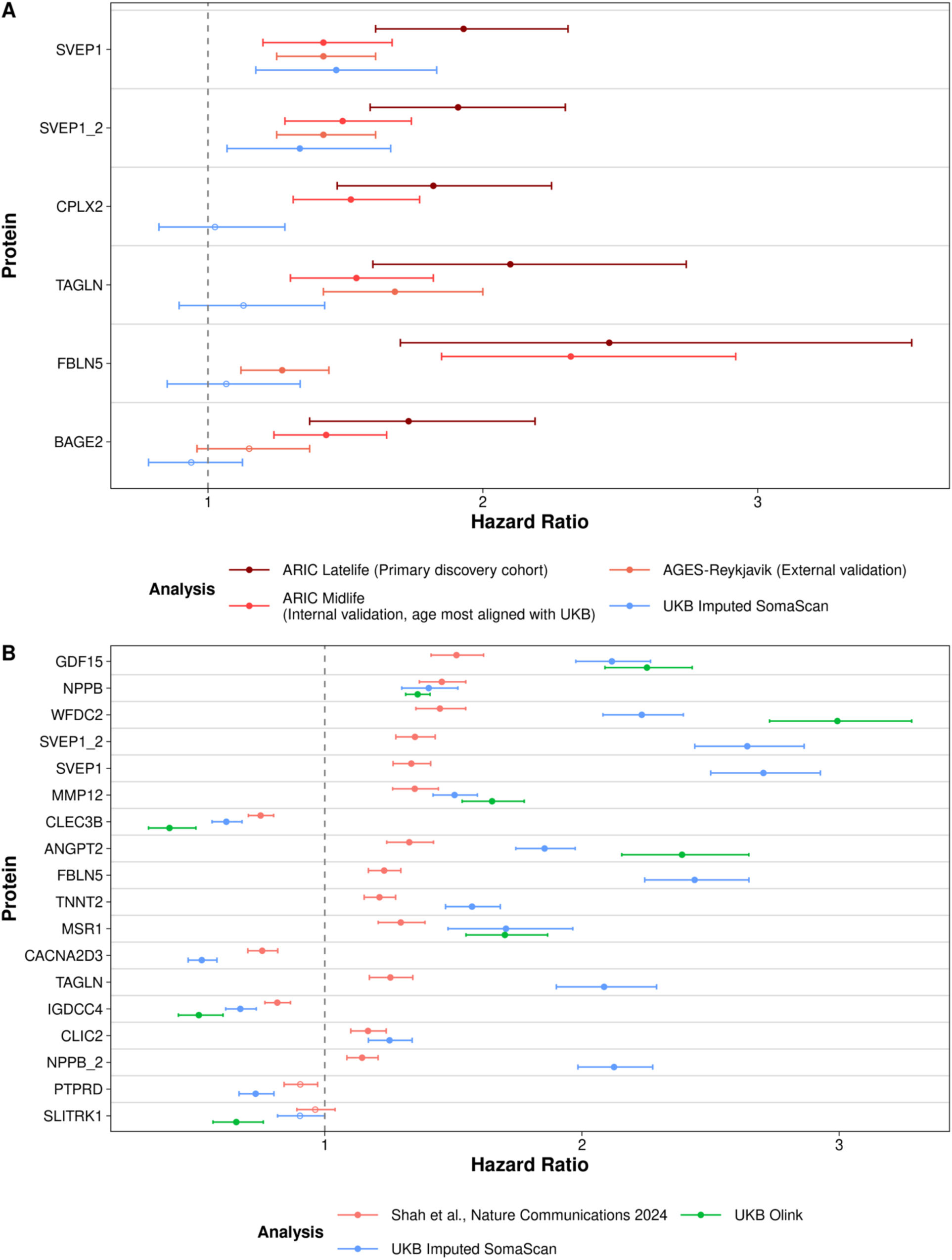
Baseline protein levels (imputed SomaScan) and the hazard of incident dementia and heart failure in UKB (Application 2 & 3). **A. Application 2 - Association between baseline protein level and incident dementia.** Five SomaScan exclusive proteins (6 measurements) were displayed. Points are the hazard of incident dementia associated with 1-unit increase of Olink-imputed SomaScan protein level at baseline, with 95% confidence interval. Proteins are ordered according to the original study (Walker et al., Nature Aging 2021)’s study findings in ARIC latelife cohorts, with SVEP1 being most significant findings that replicated in internal validation cohort (ARIC midlife, where its mean age aligns the best with UKB’s) and in external validation cohort using AGES-Reykjavik. We successfully replicated the most important signals / findings in the original study: SVEP1. SVEP1: sushi, von Willebrand factor type A (suffix indicates values measured by different aptamers), EGF and pentraxin domain containing 1; CPLX2: complexin 2; TAGLN: transgelin; FBLN5: fibulin 5; BAGE2: B melanoma antigen 2. Each protein was associated with incident dementia in separate multivariate models adjusted for age at enrollment, sex, white British ancestry, smoking status, eGFR-creatinine, APOE-ε4, BMI, prevalent hypertension, and diabetes, identical to the original study. **B. Application 3 - Association between baseline protein level and incident heart failure.** Points are estimated hazards of incident heart failure associated with a 1-unit increase in imputed protein or 1-SD Olink levels in with 95% confidence interval. ARIC: Atherosclerosis Risk in Communities Study; AGES-Reykjavik: The Age, Gene/Environment Susceptibility-Reykjavik; SVEP1: sushi, von Willebrand factor type A, EGF, and pentraxin domain-containing protein 1; WFDC2: WAP four-disulfide core domain protein 2; MSR1: Macrophage Scavenger Receptor 1, or CD204; SLITRK1: SLIT and NTRK-like family, member 1; FBLN5: fibulin 5; TAGLN: transgelin; ANGPT2: angiopoietin-2; GDF15: growth differentiation factor 15; MMP12: matrix metallopeptidase 12; TNNT2: troponin T2; CLEC3B: c-type lectin domain family 3 member B; NPPB: natriuretic peptide B; CLIC2: chloride intracellular channel 2; CACNA2D3: calcium voltage-gated channel auxiliary subunit alpha2delta 3; PTPRD: protein tyrosine phosphatase receptor type D; IGDCC4: immunoglobulin superfamily, DCC subclass, member 4; Proteins are ordered based on most descending significance (Bonferroni adjusted p-value) in the original study (Shah et al., Nature Communications 2024).

While SVEP1 was the primary focus of this replication given its centrality to the original discovery, several additional proteins reported by Walker et al. (CPLX2, TAGLN, FBLN5, BAGE2) did not replicate using imputed values in UKB (**Figure 6A**). We attribute this non-replication primarily to cohort-level differences between UKB and the original ARIC/AGES-Reykjavik cohorts, including demographic composition and potential differences in dementia ascertainment (ICD-code-based in UKB vs. protocol-driven cognitive assessment in ARIC/AGES-Reykjavik).

#### Application 3 - Improving replication of overlapping protein-outcome associations (Shah et al., *Nature Communications* 2024)

In addition to the recovery of platform-exclusive protein-phenotype associations, the third application of our imputed SomaScan values is yielding ‘more replicative and comparable’ results to SomaScan-based findings compared to using direct Olink measurements for overlapping proteins. Similar to our approach in ‘Application 2’, we adopted the same methodology as Shah et al. for our analysis (**Methods**)^32^. While the inclusion criteria of the UKB study population were different from ‘Application 1’ and ‘Application 2’, most of the demographic characteristics, such as mean age and BMI, remained identical (**Supplementary Table 1, Supplementary Table 11**). The incidence rate of heart failure per 100,000 person-years is 330.9, and the median follow-up time is 13.5 years (IQR: 12.7-14.3).

For 17 of 18 measurements (94%), the imputed SomaScan value was significant (Bonferroni-adjusted P<0.05). For 7 out of 8 available in both SomaScan and Olink, imputed SomaScan measurements consistently yielded estimates closer to the associations by Shah et al. than the estimates using native Olink measurements (**Figure 6B, Supplementary Table 13**). All four tier 2 proteins (CLEC3B, IGDCC4, NPPB, and SLITRK1) showed imputed SomaScan HRs closer to the original discovery HR than the corresponding Olink-measured HRs, supporting the validity of tier 2 membership; for tier 3 protein MSR1, imputed and unimputed estimates remained similar to one another but not closer to the original SomaScan estimate.

### Protein Performance Interpretation: Feature Importance by SHAP values & Protein Structures by Alphafold3

To examine whether feature importance patterns differ between proteins with a measured counterpart on the other platform (overlapping) and those that are purely imputed (platform-exclusive), we compared SHAP-based feature importance across both protein groups. In general, more strongly correlated proteins tend to rely on their counterpart and the top features tend to be more important (**Supplementary Table 14, Figure 7A**).

**Figure 7.**
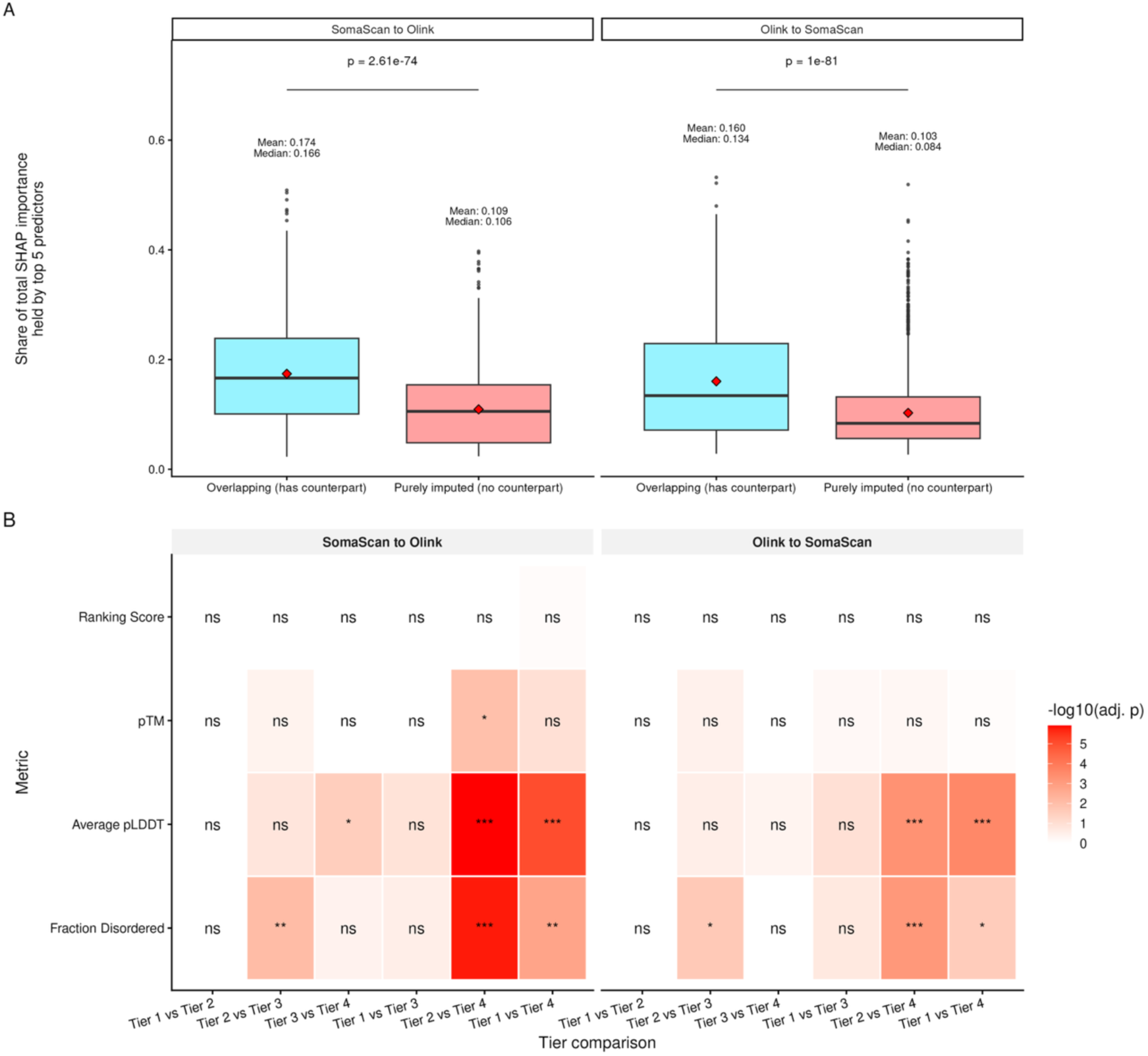
Feature Importance by SHAP, and protein structure by Alphafold3. **Panel A. ‘Concentration’ of SHAP feature importance among proteins with and without an overlapping counterpart (purely imputed).** This ‘concentration’ is quantified as the proportion of total importance held by the top 5 predictors for each protein. Boxplots show the distribution stratified by overlapping status (yes/no overlapping) and imputation direction; the red diamond marks the mean, and mean/median values are annotated above each box. Groups were compared by Wilcoxon rank-sum test (p-value shown above each facet). **Panel B. Pairwise comparisons of AlphaFold3 structural metrics (ranking score, pTM, average pLDDT, fraction disordered).** Pairwise comparisons were made using Wilcoxon rank-sum test, and p-values are adjusted for multiple testing using Bonferroni. Tile color reflects the magnitude of -log_10_ adjusted P-value. Asterisks denote significance (*: adj P <0.05, **: adj P <0.01, ***: adj P<0.001; ns = not significant). pTM: predicted template modeling; pLDDT: predicted local distance difference test.

We further investigated the reasons behind the successes in applications and observed differences in imputed correlation, focusing on FABP4 (highlighted in application 1), PILRA, and SERPINA5 (strong negative correlation) as illustrative examples. For example, the top feature contributing to both imputed correlation and model prediction accuracy of FABP4 is FABP3 in SO direction and itself in OS direction. While PILRA itself drove the correlation gain in magnitude in SO direction, its gains in OS direction were led by PILRB and NECTIN2, proteins that are highly correlated with PILRA^33,34^. Similarly for SERPINA5, SO imputation leveraged the strongly correlated SERPINA5-F2 pair to improve performance^33,35^, a feature missed in the OS direction.

Alphafold3^36^ was used to further investigate the performance difference between different protein tiers, as protein structures would directly impact the binding affinity of SomaScan aptamers and Olink antibodies. Across both directions, fraction disordered and average predicted local distance difference test (pLDDT) showed a consistent gradient with tier 4 proteins differed significantly from ‘good’ tiers such as Tiers 1 and 2 (**Figure 7B**). The structural disorder and per-residue confidence decline monotonically as reliability of the protein worsens, and the difference became significant when comparing non-adjacent tiers (**Supplementary Table 15**).

## DISCUSSION

In recent years, the exponential growth of population-scale proteomics has generated a wealth of association data. However, findings derived from one platform usually lack reproducibility when attempted on the other, one of the most frequent critiques in peer review that stifles the transition from discovery to clinical application. When non-replication occurs between characteristically different cohorts, it is often impossible to disentangle whether the failure is due to distinct environmental or biological cohort effects or simply technical platform artifacts. This uncertainty has become a rate-limiting step in the translation of epidemiological and clinical research using proteomics, and our study is motivated by this critical bottleneck.

In this study, we leveraged a machine learning-based framework to bridge the critical gap between the two dominant high-throughput proteomic platforms, SomaScan and Olink. By leveraging paired data from MESA, we constructed bidirectional imputation models that predict full proteomic panels for both platforms. Our approach is designed to addresses the systemic challenge of non-replication in large-scale proteomics mentioned above, a discrepancy that largely limits the confidence of findings and thus the number of protein candidates for potential disease biomarkers and drug targets. Our framework offers three complementary components: 1) a tiered protein reliability system that guides target prioritization or future assay optimization; 2) a bidirectional imputation to recover “exclusive” proteins; and 3) 3) improved cross-platform concordance of overlapping proteins. We demonstrated the clinical validity of this approach and its imputed values through successful replication of protein-outcome associations with BMI (application 1), incident dementia (application 2), and incident heart failure (application 3) across demographically distinct cohorts (UKB and CHS), which supports the utility of this approach for increasing replicability.

Previous evidence indicates that poor correlation between platforms could arise due to the epitope effect, or potential mismeasurement^1,25^. For medical and epidemiological researchers, the primary objective is to identify which reliable and robust associations are rooted either in high-fidelity assays or replicability across independent cohorts. Therefore, we chose the bidirectional approach rather than harmonizing two platforms’ measurements into one, as it may further deviate us from the underlying biology. The bidirectional imputation reconstructs the target signal by leveraging the high-dimensional proteome and learning the non-linear correlations between the target protein and its biological ‘neighbors.’ The objective of this reconstruction is also not to create ‘literal’ replications of the original measurements (e.g., tuning for high model performance/prediction accuracy), as the original measurements contain binding artifacts and noise. The imputation may act as a denoising step, and such imputation ensures that researchers can replicate an Olink-based discovery in a SomaScan-based cohort (or vice versa) by generating the specific counterpart values, thus validating the original signal in its native feature space. The preservation of this information would also facilitate higher replicability of epidemiological studies. One such example is SLITRK1 (tier 2, SLIT and NTRK like family member 1) from Application 3: imputed-SomaScan levels of this protein yielded a more similar point estimate than its actual Olink measures compared to the published and peer-reviewed results.

The tiered reliability system, together with the protein importance scale, offers two primary utilities for the research community. First, it enables researchers to gauge the reliability and replicability of specific findings and would potentially reduce downstream analysis costs by avoiding pursuing false positive signals. A pertinent example is the classification of GFAP (glial fibrillary acidic protein) as a tier 4 protein. This biomarker was repeatedly shown to have significant discordance between SomaScan measurement and clinical gold standard measurements in multiple independent cohorts, suggesting potential binding artifact or off-target binding. Therefore, rather than immediately pursuing such signals as valid disease markers, researchers should approach findings involving tier 4 proteins with skepticism, acknowledging a higher potential for false positives. In contrast, researchers can prioritize or pursue correlations found with tier 1 proteins as they exhibit strong cross-platform correlation independent of our imputation framework, and this was empirically validated by Application 1. While tier 1 and tier 4 have more defined measurement quality or model recoverability, tier 2 and 3 offered more insights into the moderate part and expanded the usable proteomes, as demonstrated in Application 3. Second, it offers different potential hypothesis testing opportunities for researchers with different research focuses with our protein importance annotation. While downstream analyses like protein quantitative trait loci (pQTL) could prioritize tier 1 and 2 proteins, it is also worth investigating potential contributors to the persistent discordance between platforms or between platforms and clinical gold standards for tier 3 and tier 4 proteins, especially for proteins with high scientific interest. For example, unstable structures for tier 4 proteins induced by protein-altering variants predicted by Alphafold 3 could affect one or both platforms (**Results**).

Our machine-learning-driven model was able to identify biological signals and utilize proteins’ biological neighbors even when raw measurements appeared contradictory across platforms. For instance, PILRA is known to have SomaScan measurement bias due to binding affinity (rs1859788-A, p.G78R epitope effect), with similar suspicions regarding its Olink measurement^1,37^. While the PILRA exhibited negative baseline correlations (r=-0.4), our model achieved an increase in the magnitude of negative correlation (r=-0.50) for both SO and OS directions. However, the underlying contributing features were not the same across both directions. This highlighted that while genetic variants may modify the ‘stickiness’ of probes used by one or both platforms, some biological modification was preserved and computationally retrievable. Unlike PILRA, the different gain in magnitude of negative association (|gain_SO_|=0.17, |gain_OS_|=0.05) and different feature importance indicate a different kind of modifying effect for SERPINA5.

Not limited to the overlapping targets, our framework also enables the use of non-overlapping proteins, allowing cohorts restricted to one platform to investigate targets that were inaccessible. Application 2 suggested that imputed values can act as valid proxies for absent proteomic data, enabling accessible validation in an independent and different cohort using another platform for proteomics measurement.

Our study has limitations. First, imputed values for non-overlapping proteins remain probabilistic imputations. While our use cases demonstrate utility, these predictions are subject to error and should be interpreted with appropriate caution, particularly for proteins lacking orthogonal validation. Second, validation of cross-platform imputation approaches at scale is constrained by resource scarcity, including either orthogonal validation (e.g., using ELISA or mass spectrometry) or direct validation in another cohort with paired, individual-level SomaScan & Olink measurements. This is also a broader constraint of the field, and it is precisely this scarcity that motivates future data-driven approaches to bridge various measuring platforms.

Third, our use of histogram-based gradient boosting still minimizes for the loss function rather than the biological mechanism. While it captures non-linear relationships, it does not physically adjust for binding artifacts, and other methods could also be used to explore this topic. Similarly, our validation metrics (i.e., correlation, imputation gain, and tier assignment) are constructed to measure similarities between the two platforms rather than the proximity to underlying protein levels; developing metrics capable of measuring such proximity instead of platform similarities or predictive performances remains an open methodological challenge and an important direction of future work. Last but not least, our validation applications focused exclusively on observational phenotype associations (BMI, dementia, heart failure); future work should investigate whether this imputation framework similarly improves the replicability of protein quantitative trait loci (pQTL) discovery and fine-mapping across platforms, given that genetic association analyses may be differentially sensitive to platform-specific measurement artifacts compared to phenotypic associations.

To mitigate these, we validated our findings through extensive replication of peer-reviewed phenotypes in independent cohorts, and benchmarked our tier system against available orthogonal methods.

In summary, our study helps bridge the two main proteomic measuring platforms, SomaScan and Olink, improving the replicability for proteins with adequate cross-platform fidelity. By providing a public reference for protein reliability and a method to impute datasets computationally, we contribute to dismantling the barriers imposed by platform differences and potentially accelerating the translation of proteomic discoveries.

## METHODS

### Study Cohorts

#### Multi-Ethnic Study of Atherosclerosis

The Multi-Ethnic Study of Atherosclerosis (MESA) is a prospective longitudinal cohort study that recruited 6,814 men and women aged 45 to 84 years from six communities in the United States during 2000-2002^13^. The participants were free of clinically apparent cardiovascular disease at baseline and included individuals of White, Black, Hispanic, and Chinese descent. For the current analysis, plasma protein levels were measured from samples collected at Exam 1 of 5,325 participants. Proteomic profiling was performed using both the SomaScan v4.1 Panel and the Olink Explore 3072 platform.

#### Cardiovascular Health Study

The Cardiovascular Health Study (CHS) is a population-based prospective cohort study initiated to identify risk factors for the development and progression of coronary heart disease and stroke in older adults^16,17^. The study originally recruited 5,201 men and women aged 65 years and older from four U.S. communities between 1989 and 1990, with an additional cohort of 687 African American participants recruited in 1992–1993. Circulating plasma protein concentrations were assessed using the SomaScan v4.0 platform. Measurements were performed on EDTA plasma samples collected from 3,173 participants at the 1992-1993 visit, which is the baseline for the African American cohort and year 3 for the original cohort.

#### UK Biobank

The UK Biobank (UKB) is a prospective cohort study that recruited approximately 500,000 participants from the UK during 2006-2010^14,15^. This study restricted the study population to the subset of 53,018 participants with available baseline (instance 0) plasma protein measurements. The circulating plasma protein concentration was measured using Olink Explore 3072, covering up to 2923 unique protein assays per sample. Details on UKB Olink data processing have been described elsewhere^3^.

### Preparation of Protein Data

#### Data Processing of SomaScan

SomaScan utilizes SOMAmer (aptamers) reagents that bind to the target protein, and after stringent washing that removes non-specific bindings, the SOMAmer that is specifically bound to the protein is released and quantified on a standard microarray chip via fluorescent signal and returned as the value in relative fluorescent units (RFU)^5^. SomaScan data showed no missing values in either cohort. Different sets of SOMAmers targeting the same protein were kept as distinct measurements and denoted with suffixes (e.g., protein_2/3) to preserve reagent-specific reads.

Different versions of SomaScan utilize different technical processing protocols, such as reagents, well volumes, etc. As CHS utilized the SomaLogic Panel v4.0, which includes around 5,000 proteins, while MESA used the SomaScan v4.1 panel, which includes approximately 7,000 proteins, MESA data were lifted from the v4.1 panel to the v4.0 panel in this study using the SomaDataIO R package (v6.3.0)^18^. SomaDataIO is an official and open-source R package provided by SomaLogic / BioTools, Inc. to allow users to load, manipulate, and analyze data generated by SomaScan proteomic measurements. The lifting process is a linear transformation of the raw RFU data columns between panel versions for each protein assay. The details of the lifting algorithm are well documented in the SomaDataIO R package documentation and GitHub website^18^. After lifting, the RFU values were log2 transformed, then further standardized to approximate the standard distribution of Olink NPX values.

#### Data Processing of Olink

Olink raw values were measured using PEA, and further normalized into Normalized Protein eXpression (NPX) that is proportional to the initial concentration of the plasma protein^6^. For proteins with a very low abundance or concentration in plasma, too few successful binding events would not be able to generate signals higher than the platform’s established limits of detection (LOD) threshold, and hence were marked with missing values. Proteins (columns) and participants (rows) having more than 10% missing values were excluded per quality control, and we leveraged K-nearest neighbor (KNN, R package ‘impute’ v1.62.0)^38^ imputation with 10 nearest neighbor (K=10) to impute the remaining missingness in the dataset.

Six proteins were repeatedly measured across different Olink panels, and the final protein value was chosen based on the highest target protein detectability and the largest number of NPX data points exceeding the LOD per UKB Olink processing documentation (UKB Category 1839, Resource 4654)^3^.

### Imputation of SomaScan and Olink

#### Histogram-based Gradient Boosting Regressor Algorithm

We employed a histogram-based gradient boosting regressor from the Scikit-Learn package to construct bidirection predictive model trained on the SomaScan and Olink measurements from MESA, aiming to predict SomaScan measurements from existing Olink measurements (Olink → SomaScan (OS)) and predict Olink measurements using existing SomaScan measurements (SomaScan → Olink (SO)) for cohorts (CHS, UKB) with single-platform proteomics data, using scikit-learn’s default squared-error loss and default hyperparameters.

Gradient boosting (GB) is a machine learning ensemble method that builds a predictive model. It trains a series of decision trees where each tree is explicitly designed to correct the residual errors of the previous tree^39^. This additive process minimizes a loss function, increasing the predictive ability of the model. Histogram-based gradient boosting (HGB) is a variation of this approach, optimizing computation efficiency to process large numbers of samples and features^40,41^. Instead of evaluating split points for each feature, HGB bins similar continuous input features and reduces computation complexity. It is selected for imputation as the discordance between these platforms is often non-linear and driven by different binding kinetics of the platforms. Not only can GB model non-linear relationships between proteins, but it is also robust to outliers and noise, which are common in high-throughput proteomic data due to batch effects or technical noise during and post normalization. By using these decision trees, the model can capture local variation between proteins that the global scaling methods like normalization would miss. This approach is designed to overcome issues of non-replication, therefore allowing accessible and cost-effective hypothesis testing across diverse cohorts and technologies.

For each protein measurement in the destination platform, the entire set of protein measurements from the base platform was used as input features for the histogram-based gradient boosting regressor. For the SomaScan-to-Olink direction, each of the Olink proteins was separately predicted by using all 4,972 SomaScan proteins as predictors; this yields a total of 2,871 models, including both overlapping proteins and platform-exclusive proteins. Similarly, for the Olink-to-SomaScan direction, each of the SomaScan proteins was separately predicted by using 2,871 Olink proteins.

### Training, Internal & External Validation

The MESA cohorts were separated into 75% training (N = 3,993) and 25% hold-out internal validation (N=1,332) sets. Baseline covariates reflecting general health status were compared between training and hold-out sets, including categorical variables (e.g., diabetes, hypertension, smoking status, self-reported cancer history) and continuous variables (age, BMI, eGFR) using appropriate methods (Fisher Exact for N_subgroup min_<6, Chisq for categorical, Wilcoxon for continuous variables). We calculated the baseline correlation of the overlapping proteins (N = 1,737) between SomaScan measurement and Olink measurements in the training dataset and post-imputation correlation in the hold-out internal validation set with the Pearson correlation value. In the internal validation set, the correlation between imputed protein levels and actual protein levels is also calculated using Pearson correlation coefficient. For non-overlapping proteins of each platform, only the model performance criteria (Pearson correlation) is available. Post-imputation cross-platform correlation was also reported for demographic subgroups (sex: male/female; race/ethnicity: white/black/Hispanic/Chinese).

Following model training on the MESA cohort, the resulting predictive models were applied to two independent external validation cohorts (CHS and UKB). The primary objective was to assess the cross-platform performance of the imputation models for cohorts with single-platform measurement in the setting of replicating previous proteomic studies.

### Alternative Modeling Benchmark (ElasticNet and LightGBM)

To benchmark HGB against alternative modeling approaches, we additionally trained elastic net (glmnet v5.0, R) and LightGBM models using the same predictors and train/test split for all three models, using default hyperparameters except for elastic net’s regularization parameter λ, which was selected via cv.glmnet. We quantified pairwise tier-membership concordance using Cohen’s κ, each method’s association with an independent mass-spectrometry validation dataset (Sissala et al., *Communications Chemistry* 2025) using η², and pairwise differences in correlation with this ground truth using Steiger’s Z test (cocor v1.1.4, R).

### Correlation Gain

We define baseline correlation as the Pearson r between log-2 transformed, scaled SomaScan RFU and Olink NPX values within the MESA training subset. Post-imputation correlation was similarly calculated in the MESA test subset, comparing machine-learning-imputed values against their measured counterparts (e.g., imputed SomaScan versus measured Olink). The resulting correlation gain represents the difference between these post-imputation and baseline Pearson r values. We additionally reported Spearman’s *ρ* as a supplementary, rank-based correlation metric given its insensitivity to outliers.

### Tiered Protein Reliability System

Protein tiers were defined based on the baseline and post-imputation Pearson correlation coefficients (r): 1) tier 1 proteins were categorized if they had both baseline and post-imputation r ≥0.7; 2) tier 4: both baseline and post-imputation r<0.1.

After defining tier 1 and 4 proteins, the remaining proteins were further categorized based on their post-imputation correlation. If the post-imputation correlation ≥0.4, it will be in tier 2; otherwise (r <0.4), it will be in tier 3 (**Supplementary Table 7**). To quantify sampling uncertainty in post-imputation cross-platform correlation and assess the stability of tier assignments under resampling, we performed bootstrapping with 1,000 resamples in the MESA test set (N = 1,332) and re-assigned tiers based on bootstrapped correlations. We visualized proportion of proteins with ≥5% alternative tiers within each originally assigned tiers and the 95% bootstrapped confidence interval of each proteins’ post-imputation correlation.

In addition to the conventionally used strength of correlation thresholds (0.1, 0.4, 0.7)^23^, we tested how change in these thresholds would affect the tiering assignment. We re-assigned protein tiers for every 0.02 correlation change in the ± 0.1 Pearson correlation range, and calculated the proportion of proteins assigned to a tier under various thresholds.

### Model Calibration and Uncertainty Intervals

Calibration slopes and intercepts, root mean square errors (RMSEs), and variance preservation (SD ratios) were calculated by regressing measured values (outcome) on imputed values (predictor) in the MESA hold-out test set (N=1,332). Prediction uncertainty intervals were quantified using split conformal prediction^22^, which produces distribution free prediction intervals using the calibration set (here, the hold-out test set).

### Protein Localization and Importance

The importance of a protein is evaluated through 2 main criteria, clinical relevance and overall scientific attention, across two sources: the Human Protein Atlas (HPA)^26,27^ and MalaCards, the human disease database from the GeneCards Suite^28^. HPA highlights a protein’s established druggability and clinical relevance by indicating whether this protein is an “FDA-approved drug target” or a “potential drug target” in its “Protein Class” section. On the other hand, we quantify the overall scientific attention of a protein by the sum of relevant publications of that gene symbol or protein name in the MalaCards database. A previous publication had described the data mining and database-building process of MalaCards in detail^28^. In Brief, MalaCards searched the protein and/or gene name in databases like DISEASE^29,30^, Novoseek, or PubMed to establish the association between a protein and/or gene with a specific publication. We curated the information from MalaCards using GeneALaCart and calculated the aggregated sum of publications associated with a protein or gene symbol.

We interpret the protein importance as the following: 1) FDA-approved drug targets or potential drug targets have more clinical relevance than proteins that have no available information in this criterion, and 2) the higher the total number of publications, the more overall scientific attention it receives and hence the more important it is to the research community.

Raw protein localization was obtained through HPA protein class^26^. For 5 proteins (LILRA3, MENT, LEG1, DDX58, KIR2DL2) with missing location information in HPA, UniProt subcellular location was obtained^42^. We further quantified the localization of proteins into 3 major classes: secreted or plasma, membrane, or intracellular.

### Orthogonal Validation of the Tiered Reliability System

We compared our tier assignments against independently collected and derived MS-Olink correlations reported by Sissala et al. (Communications Chemistry, 2025)^8^, which quantified concordance between Olink and mass spectrometry. For proteins with available comparisons (N=821), we examined the subgroup difference between MS-Olink correlation and our assigned tiers using pairwise Dunn’s test to examine whether our tier assignment reflects the quality of protein measurement. P-values were adjusted for multiple testing using Bonferroni.

### Application of Imputed Protein Measurements

#### Application 1 – Tier 1 Protein for Consistent Associations (BMI as example)

We associate all the proteins with Body Mass Index (BMI), a common continuous phenotype in both CHS and UKB, using all proteins from their actual protein measurements (CHS: SomaScan, UKB: Olink) and tier 1 proteins of the actual measurements and imputed measurements, controlling for age, sex, and race. For CHS and UKB, the predictive modeling utilized elastic net regression using the “glmnet” R package to assess the association between the protein features and BMI individually. To ensure optimal performance of feature selection and elastic net, the mixing parameter *a* and regularization parameter λ were tuned using 10-fold cross-validation within the training split (80%) only. Then the model’s final performance was evaluated in the 20% hold-out test set. Unpenalized covariates (age, sex, and categorical race) were included and assigned a penalty factor of 0. The model performance is measured by the variance of BMI explained by model adjusted by non-zero model predictors (adjusted R^2^). The feature importance of each protein was then quantified using the permutation feature importance (PFI) on the test set, where the importance of a protein to the model prediction was measured by the drop in R^2^ after randomly shuffling that protein’s values. Adjusted R^2^ (1 − (1−R^2^)(n−1)/(n−p−1)) accounts for both the number of predictors (p) and sample size (n), and was reported alongside raw R^2^ given that the full-panel and tier1-only models differ substantially in predictor counts (1,737 vs. 334) and the UKB and CHS cohorts differ substantially in sample size (41,185 vs. 3,153).

#### Application 2 – Enabling Replication of Non-overlapping Proteins (Walker et al., *Nature Aging* 2021)

We also conducted a replication proteomic study of the findings reported by Walker et al., which utilized the SomaScan v4.0 platform. Separate from the tier membership informed analysis, the primary goal of this replication is to demonstrate the performance of non-overlapping proteins between SomaScan and Olink, a common obstacle faced when replicating study findings across cohorts. Briefly, Walker et al. leverage Atherosclerosis Risk in Communities (ARIC) cohort’s data from visit 5 (late-life) as discovery, and validated their findings internally in ARIC midlife (visit 3) and externally in the AGES-Reykjavik study. They utilized Cox proportional hazard regression model to examine the association between the standardized level of proteins (SomaScan) and the incidence of dementia, controlling for age, sex, race-center, education (**Supplementary Table 11**), APOE-ε4, eGFR-creatinine^43^, BMI, prevalent cases of diabetes, hypertension, and smoking status, as their primary analysis methods. To ensure methodological consistency, we harmonized definitions of covariates, exposures, and outcomes definition with the original analysis. Due to the absence of protocol-based dementia and cognitive assessment in UKB, we approximated dementia phenotype with available ICD-9 and ICD-10 codes from self-report and HESIN records.

UKB is more phenotypically similar to ARIC midlife in terms of mean age. Therefore, the goal of replication is to replicate the found associations in ARIC midlife replication in UKB using imputed SomaScan measures. Prior to the association analysis using Cox proportional hazard model, KNN imputation (k=10) was used to impute missing covariates.

#### Application 3 - Improving Cross-platform Protein Association Replication. (Shah et al., *Nature Communications* 2024)

Additionally, we performed another replication study of Shah et al.’s 2024 study on proteomic association with incidence of heart failure^32^. In summary, their study cohort also utilized ARIC midlife and late-life visits, coupled with an external validation cohort of the Trøndelag Health (HUNT) Study. Shah et al. also leveraged Cox proportional hazard model to assess the association between each individual protein and incidence of Heart Failure, adjusting for age, sex, BMI, eGFR based on creatinine (CKD-EPI), race, sex, smoking status, and prevalent cases of coronary artery disease, diabetes, atrial fibrillation, and hypertension. Likewise, we performed the replication analyses with the same set of covariates, exposures, and phenotypes (ICD codes).

### Feature Importance

Feature importance was calculated using SHAP TreeExplainer with tree-path-dependent perturbation. For each feature, we calculated the mean and standard deviation of the absolute SHAP value across the test set. Features were then ranked in descending order of mean importance, and the first K features (K=1,2,3,4,5,7,10,…,50) were used to retrain the model to assess their contribution to model performance (imputed vs. actual value) and to imputed correlation (e.g., SomaScan vs. SomaScan-imputed-Olink in overlapping proteins).

### Alphafold 3

We used AlphaFold 3 to predict protein structures as further validations. The canonical FASTA sequences were retrieved from UniProt and then submitted to the Alphafold 3 server, performing multiple runs for each protein. For each outputted JSON files, features like ranking score, pTM, average pLDDT, chain pTM mean, chain pair iPTM mean, chain paired PAE min mean, and fraction disordered are outputted. Methods in calculating these features have been described in the Alphafold3 literature^36^. For each protein, we averaged the metrics and analyzed them to identify structural factors linked to tier stratification.

## Data Availability

Downloadable protein information data can be found on the Human Protein Atlas website (https://www.proteinatlas.org/about/download). MalaCards for genes and proteins for academic use can be used with the GeneALaCart tool within the GeneCards Suite at https://genealacart.genecards.org/. Individual-level data from the Trans-Omics for Precision Medicine (TOPMed) program are available upon submission of a research proposal to the Data Access Committee through the Database of Genotypes and Phenotypes (dbGaP) at https://dbgap.ncbi.nlm.nih.gov/. Individual-level data from the UK Biobank are available to registered researchers via the UK Biobank Access Management System (https://www.ukbiobank.ac.uk) with an approved application; this study was conducted under application number 7089.

## Code and Model Availability

GitHub: https://github.com/linkeli742/somascan_olink_cross-platform_imputation; Hugging Face: https://huggingface.co/AntonyT1207/soma-olink-crosswalk-models.

## Acknowledgement

Molecular data for the Trans-Omics in Precision Medicine (TOPMed) program was supported by the National Heart, Lung and Blood Institute (NHLBI). Core support including centralized genomic read mapping and genotype calling, along with variant quality metrics and filtering were provided by the TOPMed Informatics Research Center (3R01HL-117626-02S1; contract HHSN268201800002I). Core support including phenotype harmonization, data management, sample-identity QC, and general program coordination were provided by the TOPMed Data Coordinating Center (R01HL-120393; U01HL-120393; contract HHSN268201800001I). We gratefully acknowledge the studies and participants who provided biological samples and data for TOPMed. “Multi-Ethnic Study of Atherosclerosis (MESA)” (phs001416) Phenotype harmonization, data management, sample-identity QC, and general study coordination, were provided by the TOPMed Data Coordinating Center (3R01HL-120393-02S1), and TOPMed MESA Multi-Omics (HHSN2682015000031/HSN26800004). The MESA projects are conducted and supported by the National Heart, Lung, and Blood Institute (NHLBI) in collaboration with MESA investigators. Support for MESA is provided by contracts 75N92025D00022, 75N92020D00001, HHSN268201500003I, N01-HC-95159, 75N92025D00026, 75N92020D00005, N01-HC-95160, 75N92020D00002, N01-HC-95161, 75N92025D00024, 75N92020D00003, N01-HC-95162, 75N92025D00027, 75N92020D00006, N01-HC-95163, 75N92025D00025, 75N92020D00004, N01-HC-95164, 75N92025D00028, 75N92020D00007, N01-HC-95165, N01-HC-95166, N01-HC-95167, N01-HC-95168, N01-HC-95169, UL1-TR-000040, UL1-TR-001079, UL1-TR-001420, UL1TR001881, and R01HL105756. The authors thank the MESA participants and the MESA investigators and staff for their valuable contributions. A full list of participating MESA investigators and institutions can be found at http://www.mesa-nhlbi.org. This Cardiovascular Health Study (CHS, phs001368) research was supported by NHLBI contracts HHSN268201200036C, HHSN268200800007C, HHSN268201800001C, N01HC55222, N01HC85079, N01HC85080, N01HC85081, N01HC85082, N01HC85083, N01HC85086, 75N92021D00006; and NHLBI grants U01HL080295, R01HL087652, R01HL103612, R01HL105756, R01HL120393, U01HL130114, R01HL144483, and R01HL172803 with additional contribution from the National Institute of Neurological Disorders and Stroke (NINDS). Additional support was provided through R01AG023629 from the National Institute on Aging (NIA). A full list of principal CHS investigators and institutions can be found at CHS-NHLBI.org. The lead authors want to thank A.J.B. for his insights and support during a critical turning point of this research.

## Author Contribution declaration

LL, AA, ID, ZY, AP conceived and designed the study. LL, YT, QZ performed the computational and statistical analyses. RT, KT, BY, CB, RD, RD, MT, JB, TA, BP, UT, JC, SR, JR, PG, RG contributed TOPMed cohort data. MH, SF, WH, AC provided technical support and resources. LL wrote the manuscript with input from AA, ZY, GP, JN, LM, and PN. AA, PN, and ZY supervised the study. ZY acquired funding. All authors reviewed and approved the final manuscript.

## Supplementary Figures

**Supplementary Figure 1.**
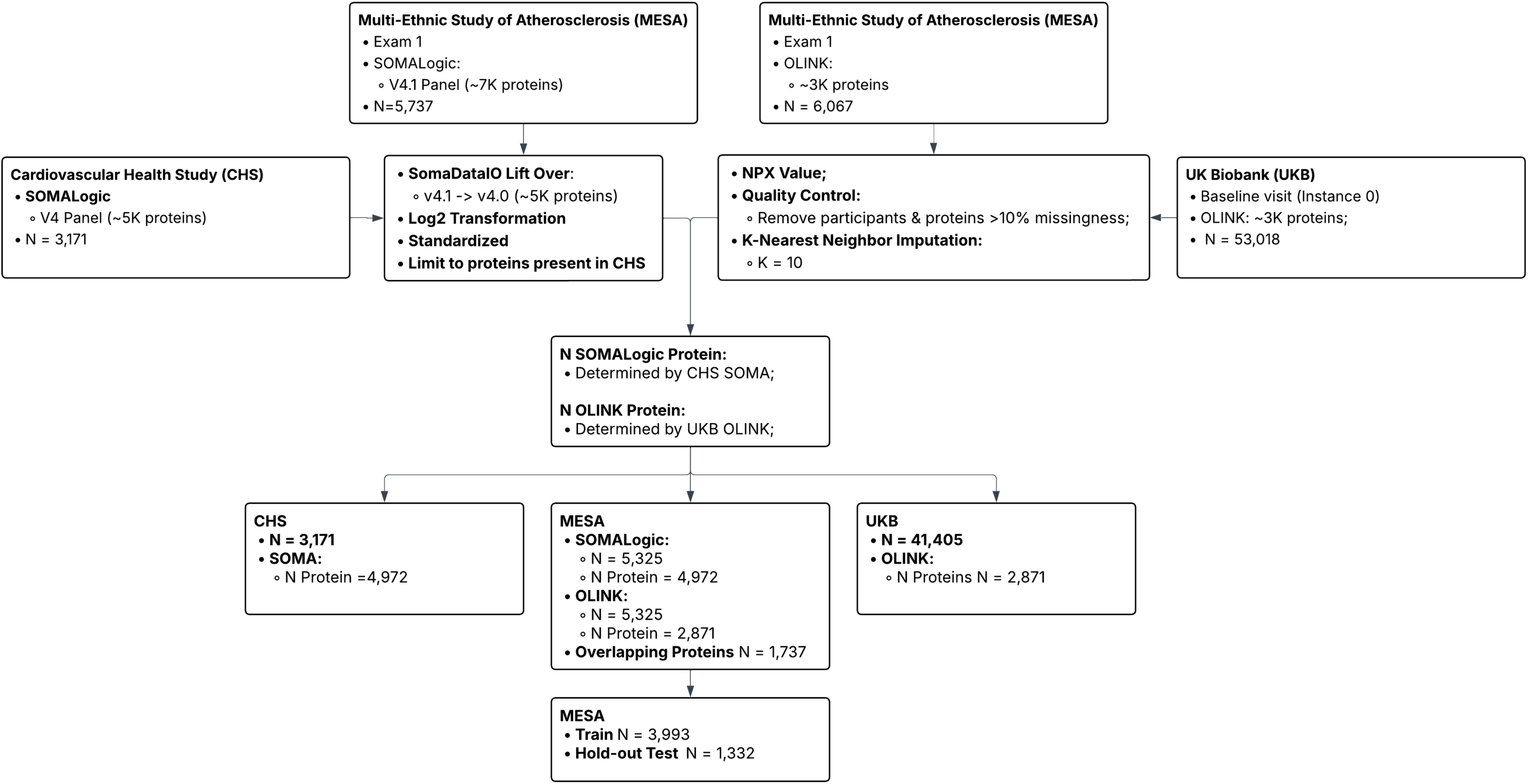
Consort diagram of sample and protein quality control in three cohorts: MESA, UKB and CHS. MESA: Multi-Ethnic Study of Atherosclerosis; UKB: UK Biobank; CHS: Cardiovascular Health Study.

**Supplementary Figure 2.**
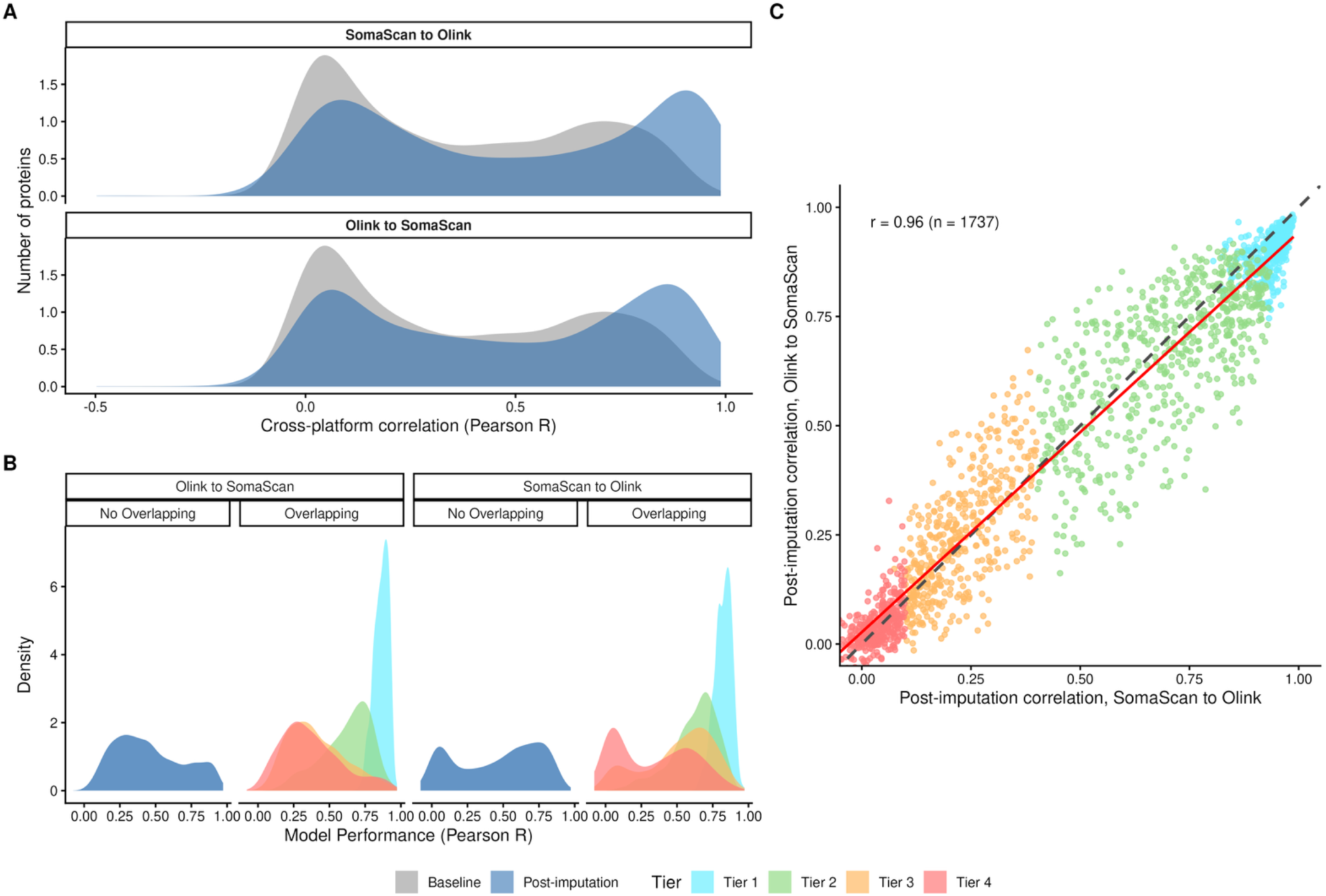
Shift in cross-platform correlation and model predictive performance (Pearson correlation) in the MESA held-out test set (N=1,332). **Panel A. Density plot of baseline and post-imputation crossplatform correlation stratified by imputation direction measured by Pearson correlation; Panel B. Density plot of model predictive performance stratified by imputation direction and overlapping status of the full panel proteins (SomaScan N=4,972; Olink N=2,871), colored by assigned reliability tiers (overlapping proteins only); Panel C. Concordance between the post-imputation correlation of overlapping proteins (N=1,737) of two imputation direction.** Each dot represents one protein. The black dashed line is x=y, meaning the post-imputation correlation is identical in both directions. The proteins/dots above the dashed line have higher post-imputation correlation in Olink to SomaScan direction, vice versa. The red line represents the fitted line using y ∼ x geom_smooth function, and the concordance between the two post-imputation correlation is measured by Pearson correlation. Tier 1 and Tier 4 proteins have more similar post-imputation correlation between 2 imputation directions compared to Tier 2 and 3 proteins. MESA: Multi-Ethnic Study of Atherosclerosis.

**Supplementary Figure 3.**
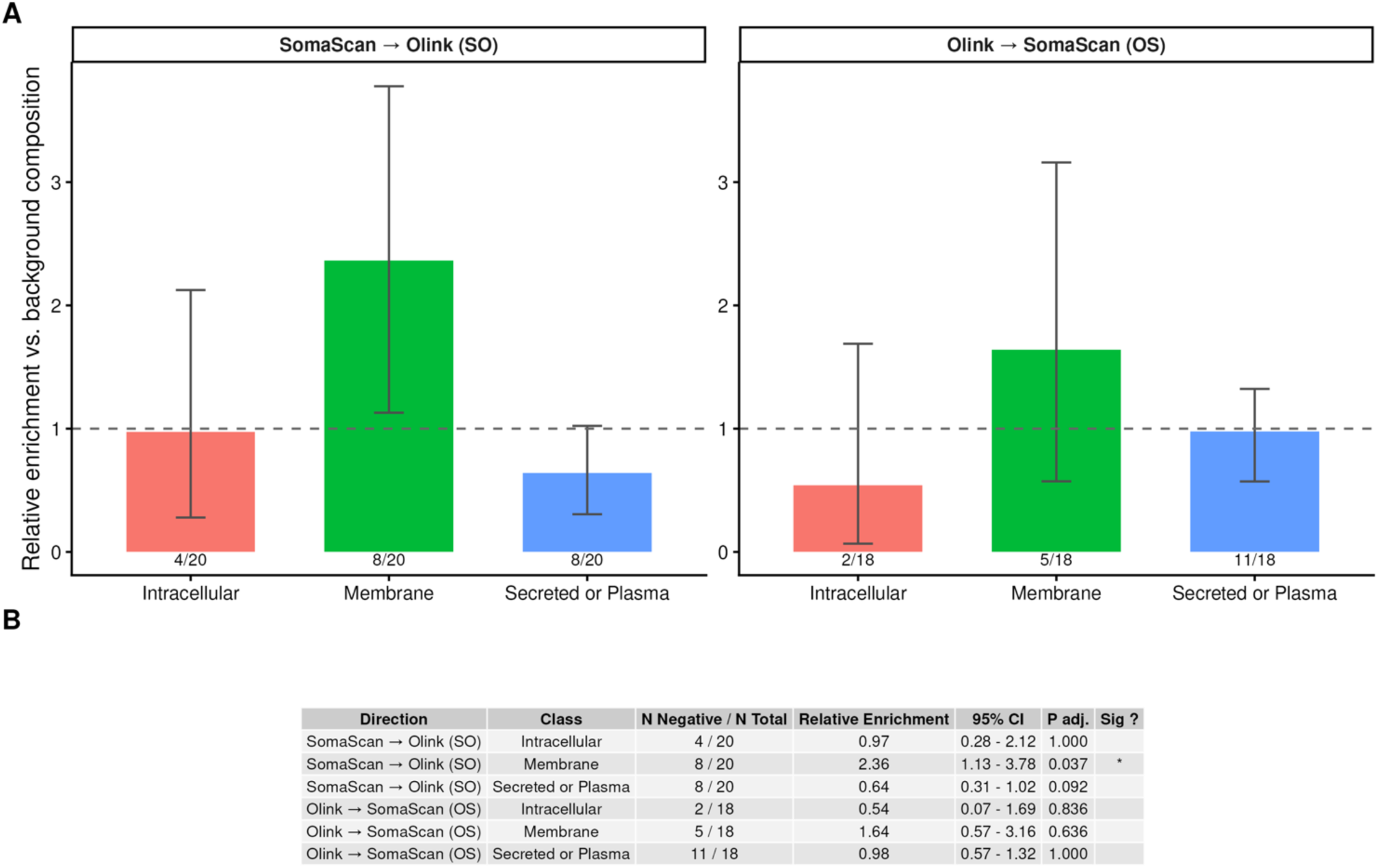
Proteins (N_SO_=20, N_OS_=18) with meaningful negative gain (gain < -0.05) by localization. **Panel A**. Relative enrichment in each direction is calculated as: Relative enrichment = (proportion of the negative-gain set in a class) / (proportion of all 1,737 overlapping proteins in that class). Dashed line: no enrichment (ratio = 1). 95% Confidence intervals were calculated using exact Binomial test. *: adjusted P<0.001

**Supplementary Figure 4.**
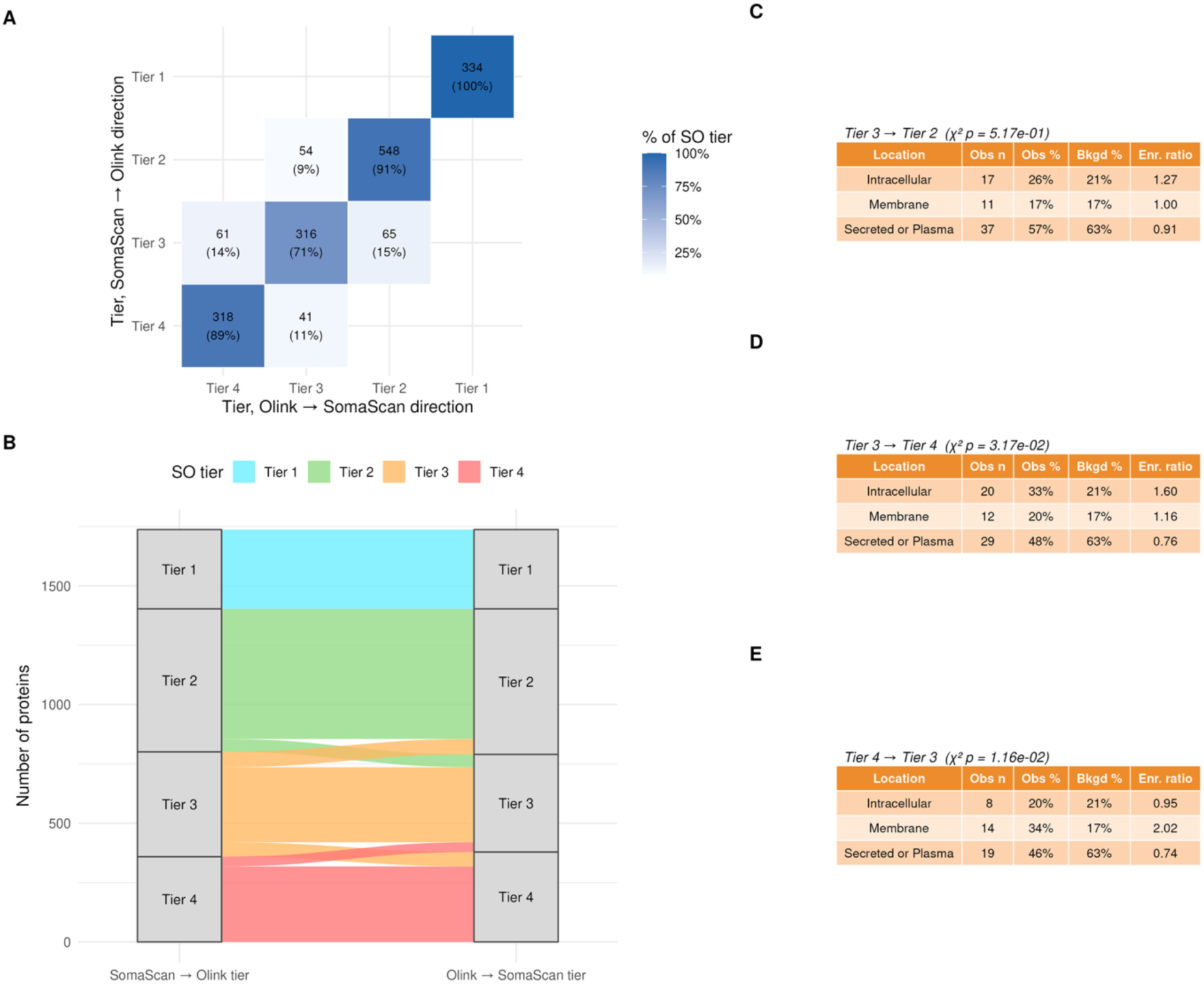
Shift in tiers by direction and protein localization enrichment for the overlapping proteins (N=1,737). **Panel A. Confusion matrix of tier assignments across 2 imputation directions for overlapping proteins.** Cell values show protein count and the row-wise (based on tier assignment in SO direction) percentage. **Panel B. Sankey Diagram of change in tiers between the imputation directions. Panel C-E. Protein localization class (Intracellular, Membrane, Plasma) composition among proteins with tier changes. (C)Tier 3 to 2. (D) Tier 3 to 4, and (E)Tier 4 to 3**. The proportion of proteins within each localization class is compared with their background frequency using Chisq test to determine if one localization class was enriched for a specific switch in tiers.

**Supplementary Figure 5.**
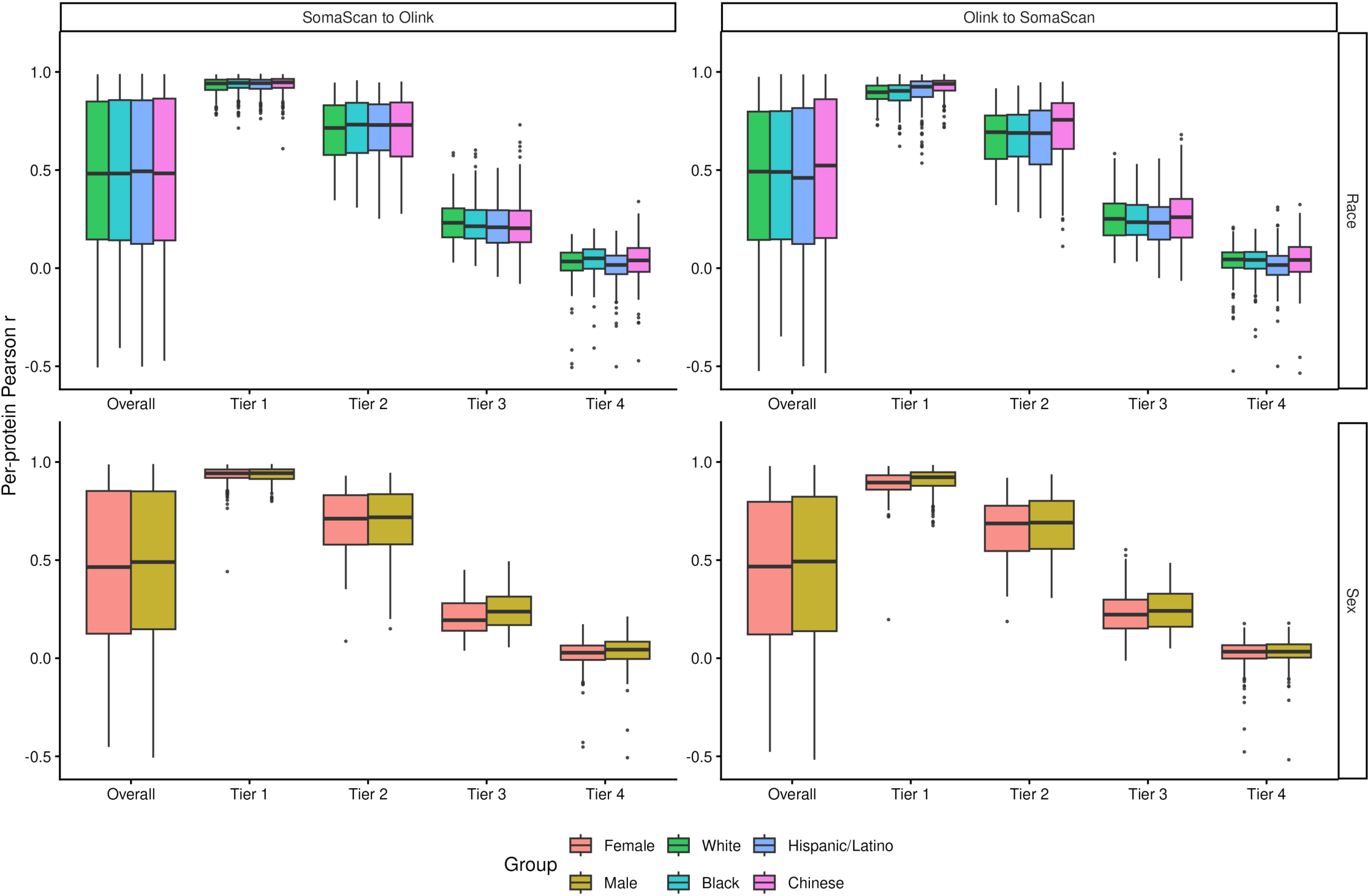
Post-imputation Pearson r of overlapping proteins(N=1,737) by MESA subgroups in test set (N=1,332). Boxplot for post-imputation correlation for each tier for the full test set and stratified by demographic subgroups is used. Demographic subgroups of sex (Male, Female) and race/ethnicity (White, Black, Hispanic/Latino, Chinese) were used. MESA: Multi-Ethnic Study of Atherosclerosis.

**Supplementary Figure 6.**
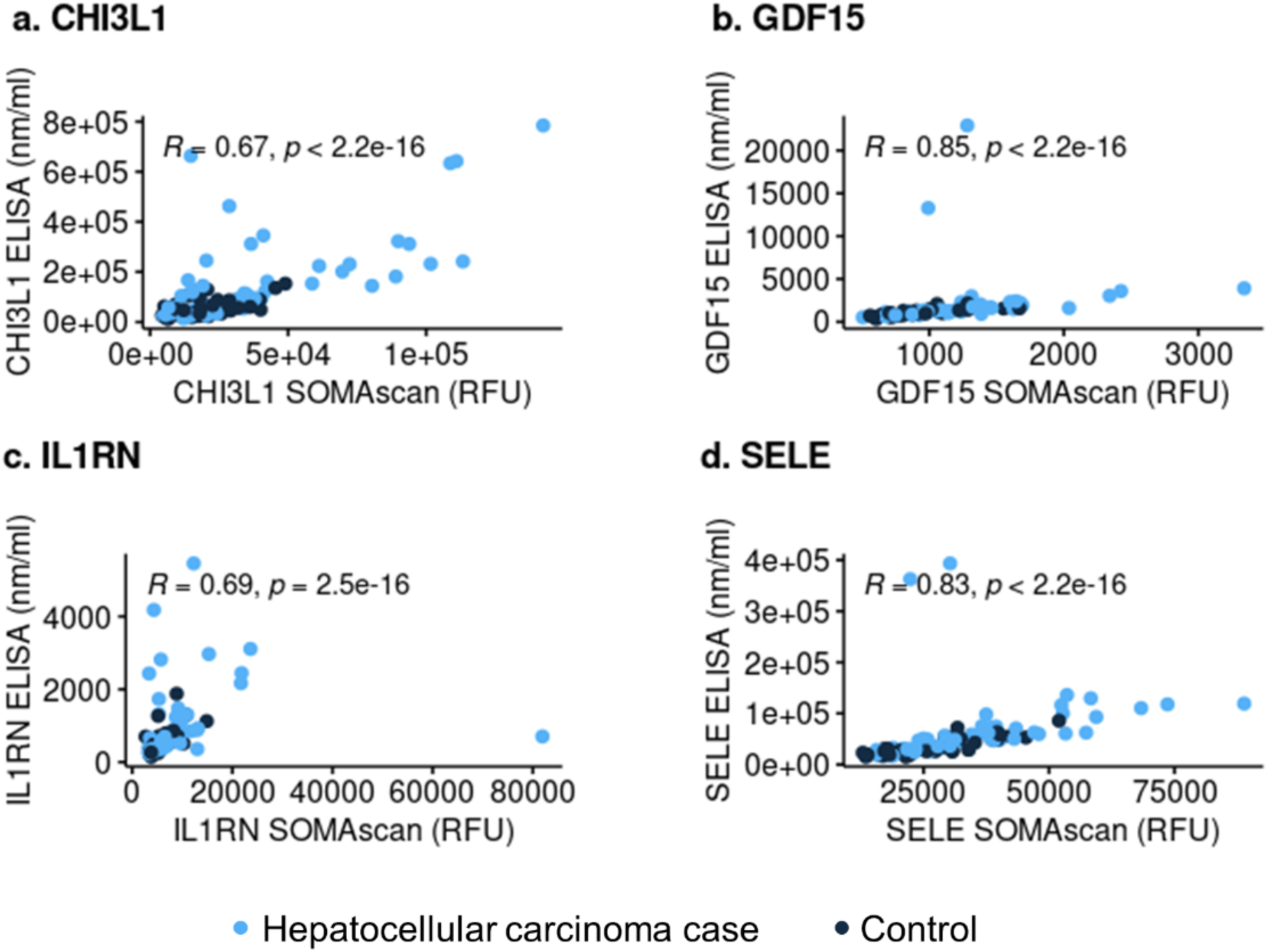
Spearman correlations between the SomaScan and ELISA protein levels in the Nurses’ Health Study and Health Professionals Follow-up Study (N=54) Re-use with permission from lead author Xinyuan Zhang. Figure also published as Supplemental Figure 2 in Zhang et al., Journal of the National Cancer Institute 2024, doi: 10.1093/jnci/djae079.

**Supplementary Figure 7.**
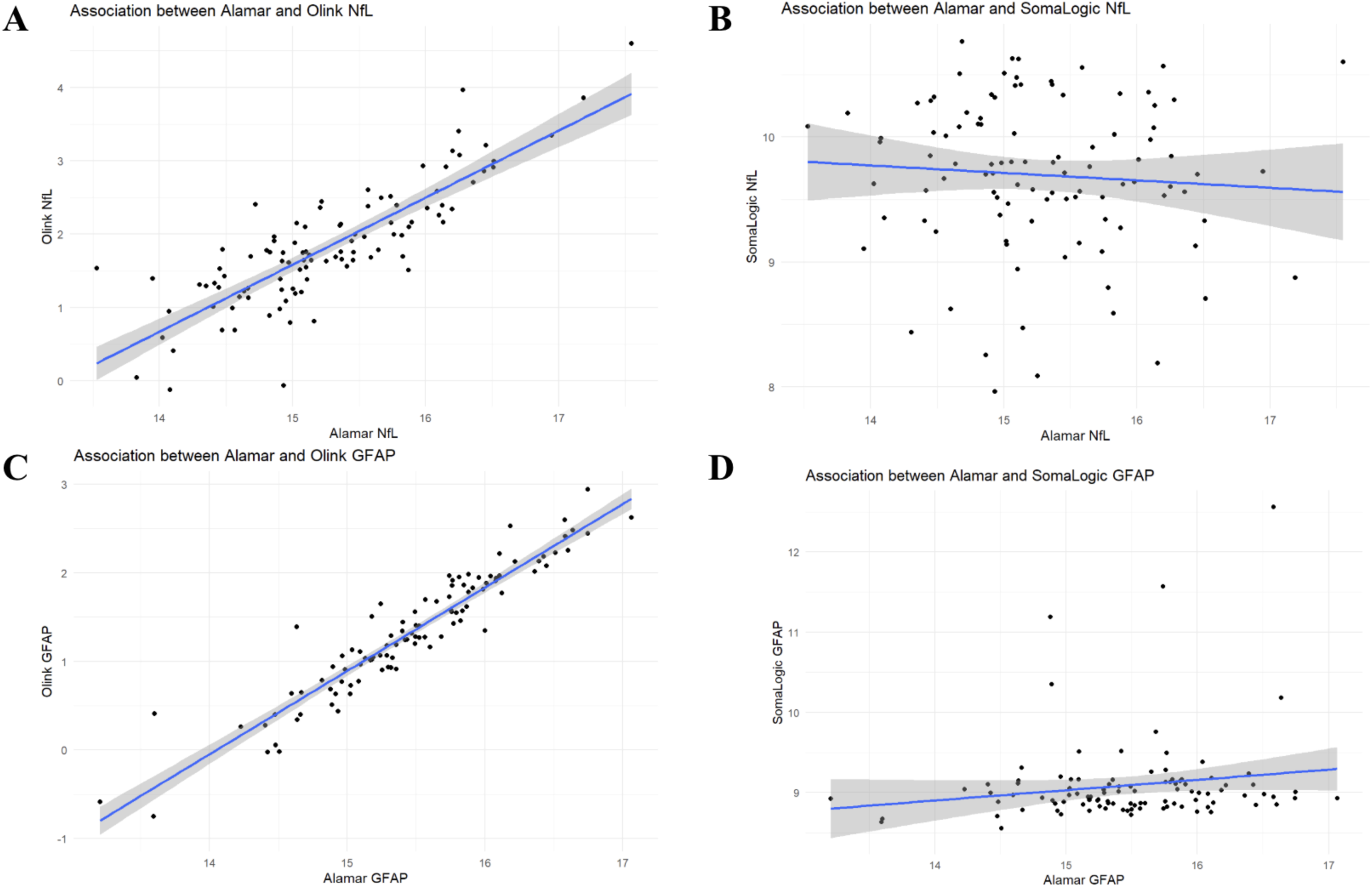
Spearman Correlation between Alamar, SomaScan and Olink measurements in Atherosclerosis Risk in Communities (N=101). Correlation calculated between Alamar and SomaScan and Olink (N∼101). The Simoa (ELISA based, gold standard) had a high correlation with Alamar assays (>0.9 after excluding 1 outlier) NfL: neurofilament light polypeptide; GFAP: glial fibrillary acidic protein;

**Supplementary Figure 8.**
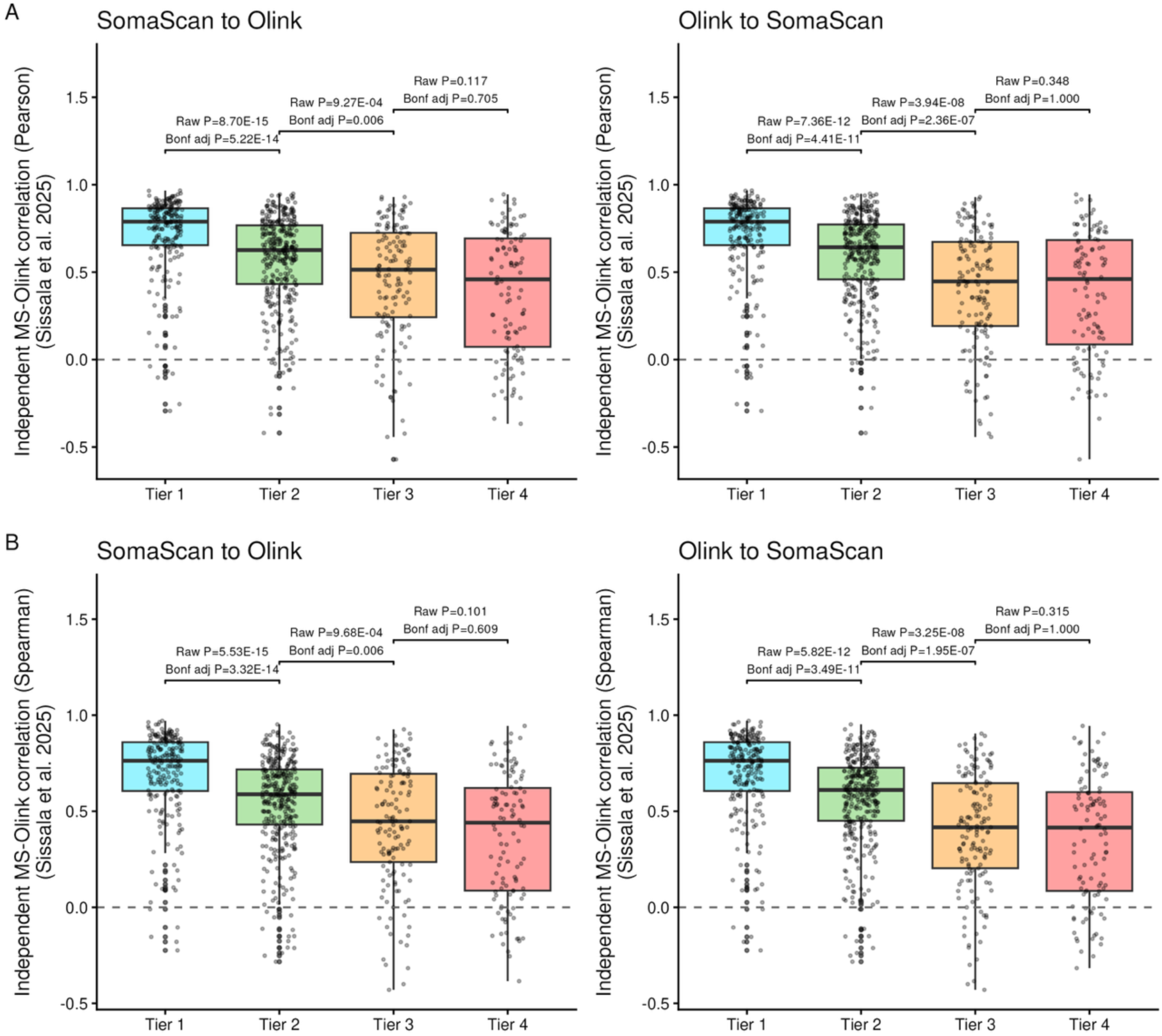
Bidirection tier assignment compared to MS-Olink correlation in Sissala et al., 2025. Panel A. Boxplots for distribution of independent MS-Olink correlation (Pearson correlation) from Sissala et al., 2025 stratified by our tier list. Panel B. Boxplots for distribution of independent MS-Olink correlation (Spearman correlation) from Sissala et al., 2025 stratified by our tier list. Dunn’s test was used to test the pair-wise MS-Olink correlation between assigned tiers.

**Supplementary Figure 9.**
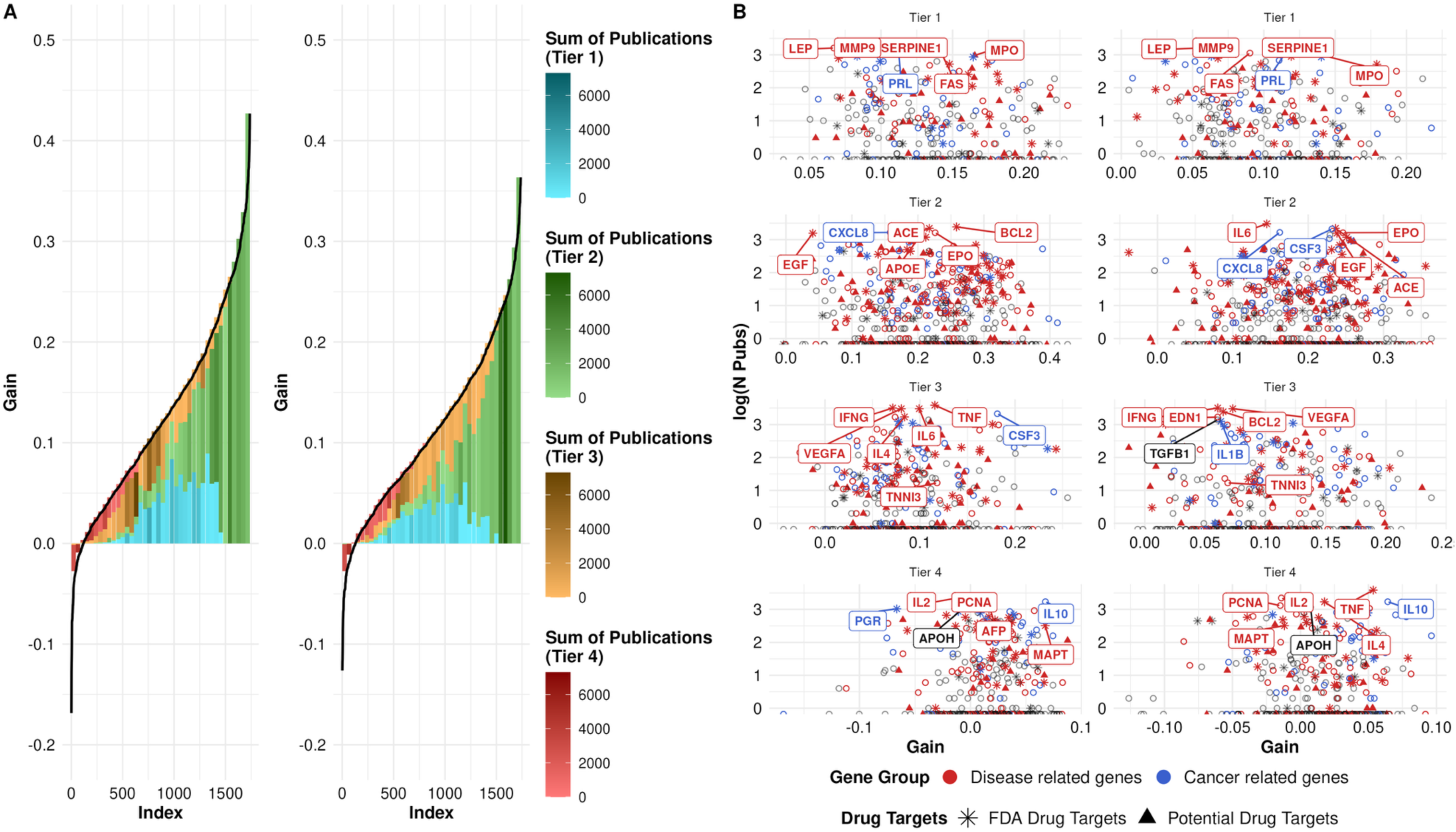
Protein importance of 1,737 overlapping proteins. **Panel A. Binned importance by correlation gain (left: SomaScan to Olink; Right: Olink to SomaScan)**; The Y-axis illustrates the gain in correlation (post-imputation Pearson correlation minus baseline Pearson correlation) for 1,737 overlapping proteins between the two platforms. X-axis represents the protein index, ranked in ascending order of the gain. All proteins were binned into 220 bins based on their gain for better visual demonstration of importance. Importance is defined by the sum of publications of a given protein through text mining provided by GeneCards. Within individual bins, the size of each color block (each protein tier) is proportional to the proportion of: (number of proteins in that tier / total number of proteins in the bin); the larger the proportion, the more proteins in the corresponding tier compared to other tiers present in the bin. In each bin, the aggregated number of publications of all proteins grouped by tiers is calculated: the darker the color, the more total publication proteins have and hence more important the proteins are. **Panel B. Scatter plot of log(number of publications) stratified by tiers (left: SomaScan to Olink; right: Olink to SomaScan)**. Each point represents one protein, colored by gene group (disease or cancer association), and shaped by druggability (FDA-approved or potential drug target). Y-axis denotes the log(number of publications) obtained using GeneALaCart.

## Supplementary Tables

**Table S1.**
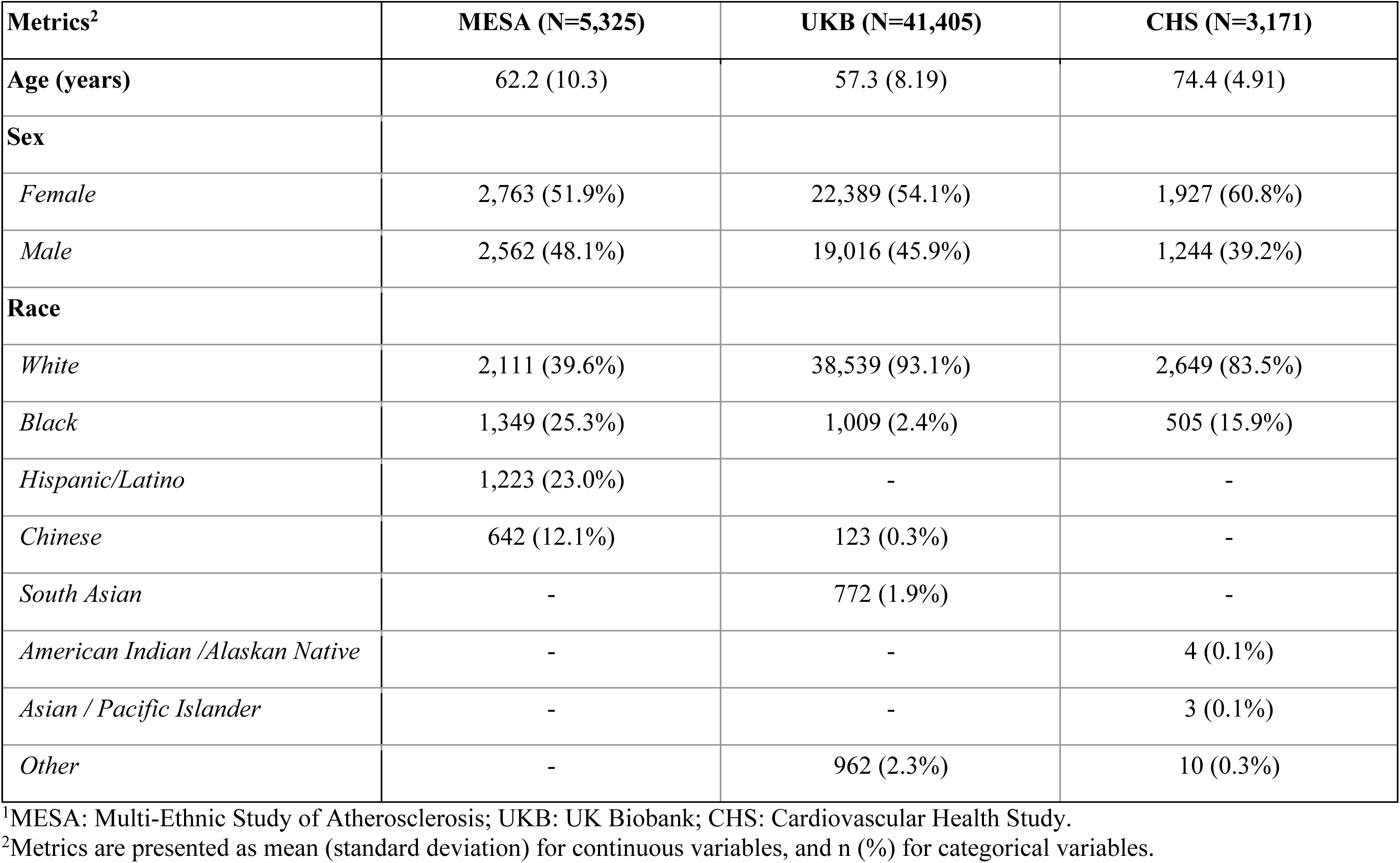
Demographics of the 3 study cohorts (MESA, UKB, CHS)^1^.

**Table S2.**
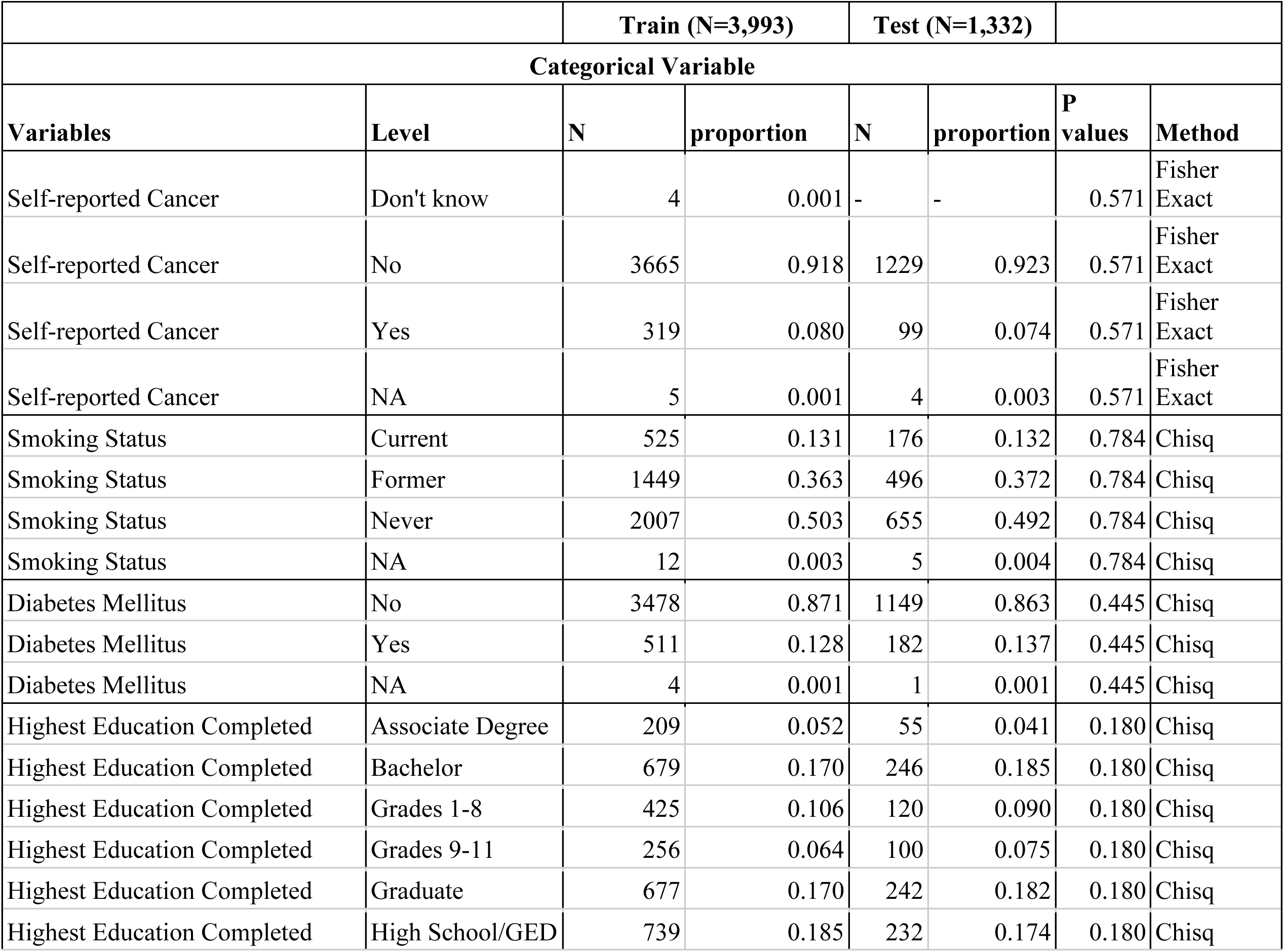
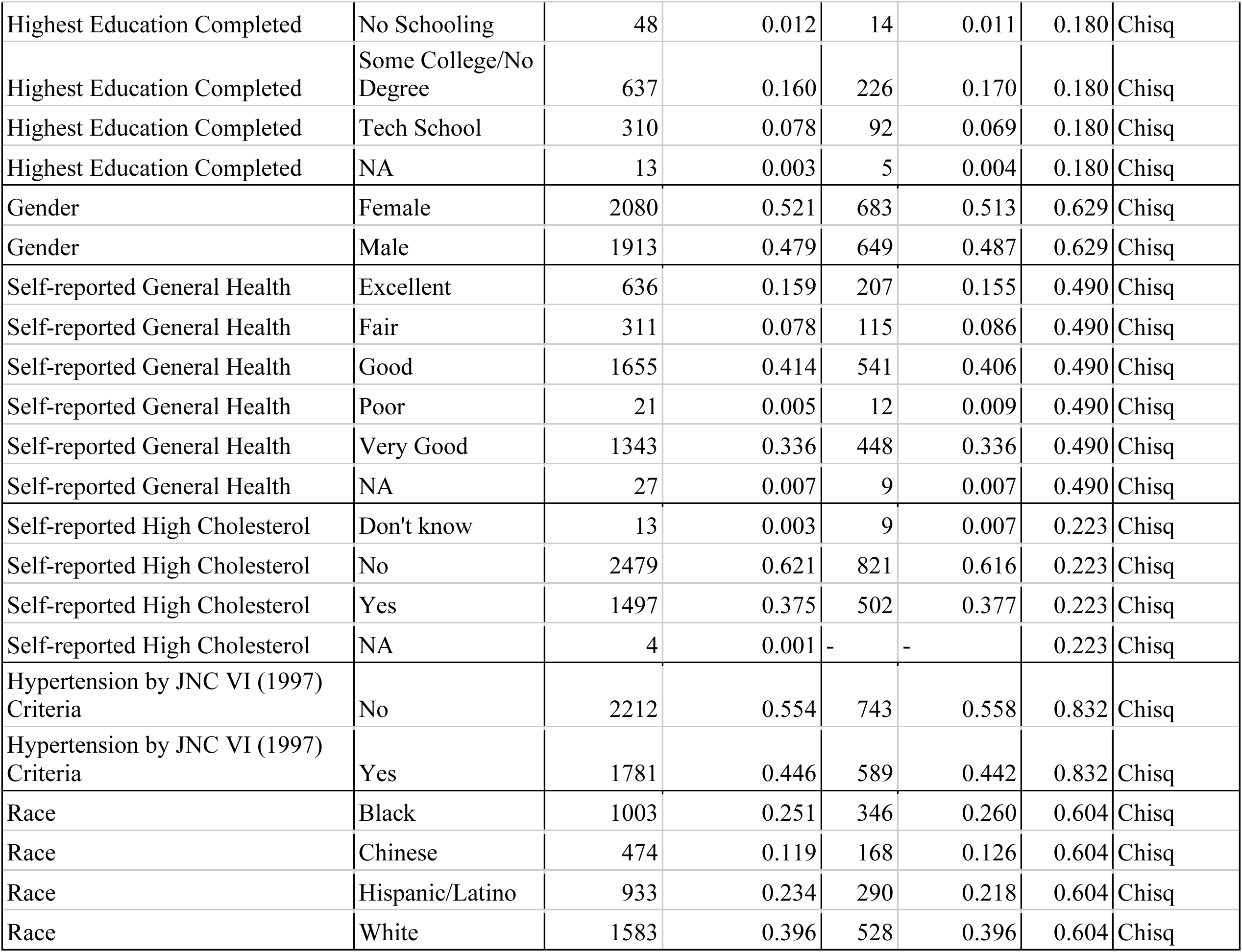
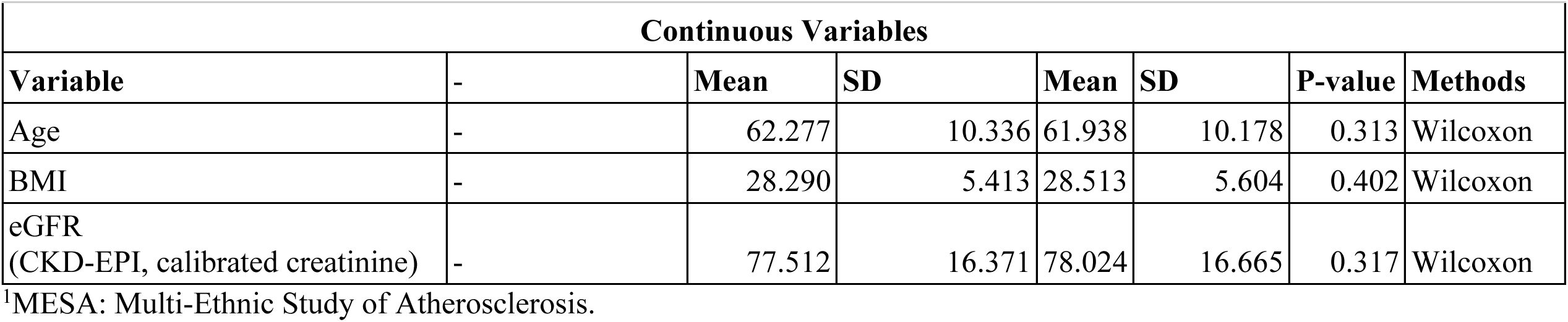
Comparison of health status between MESA1 train (N=3,993) and test (N=1,332) sets.

**Table S3.**
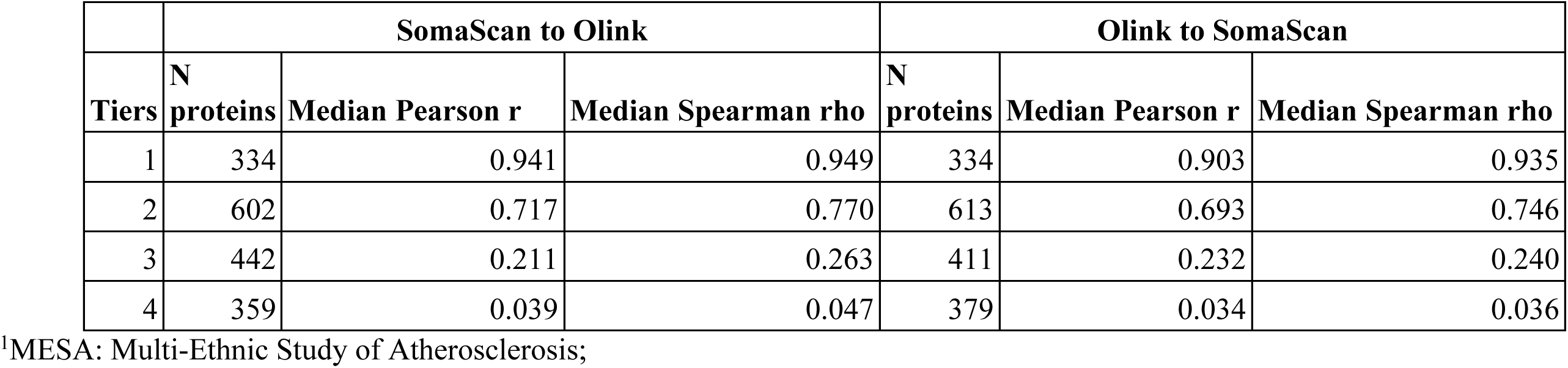
Post-imputation Pearson and Spearman correlation of 1,737 proteins in MESA1 test set (N=1,332).

**Table S4.**
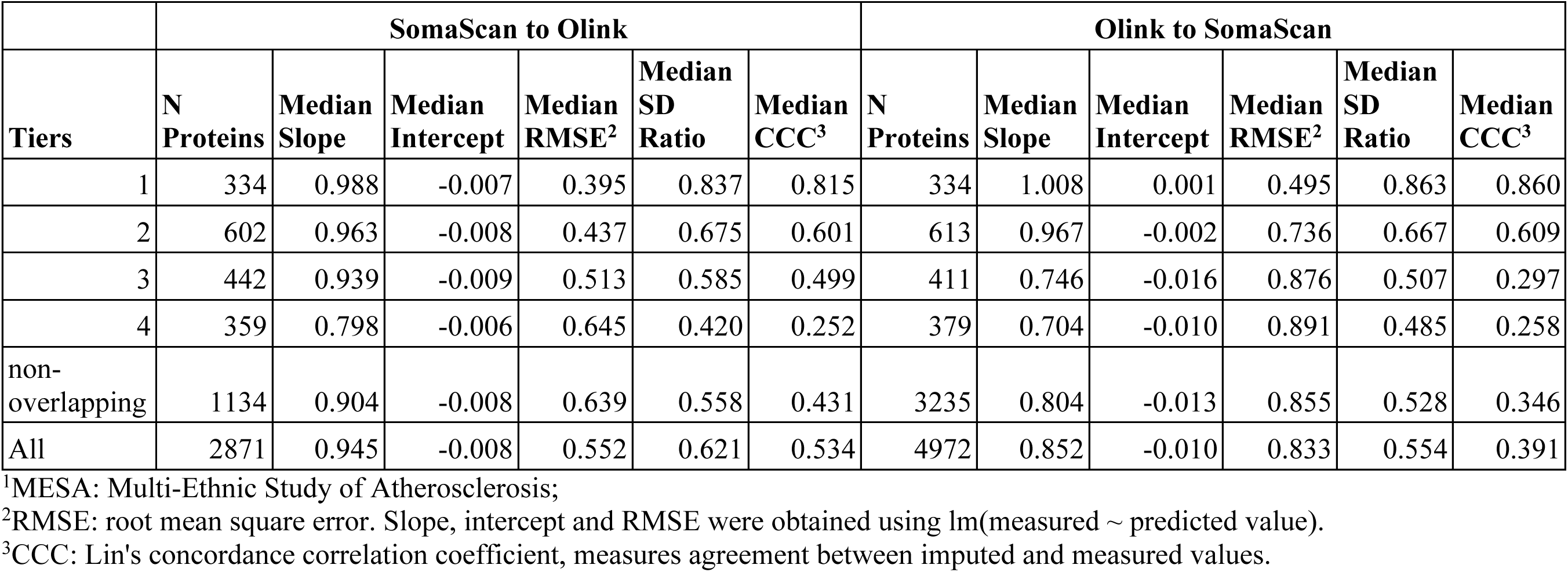
Calibration of SomaScan (N=4,972) and Olink (N=2,871) proteins in the MESA1 test set (N=1,332).

**Table S5.**
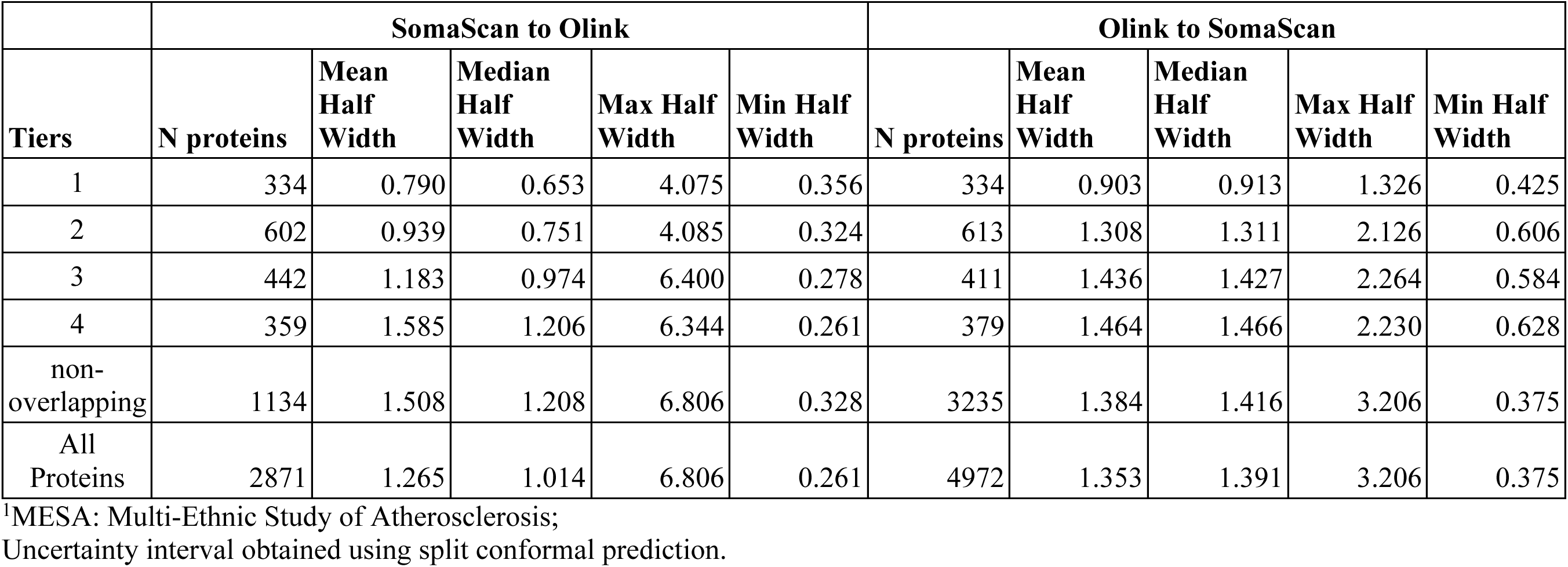
Uncertainty interval using split conformal prediction for SomaScan (N=4,972) and Olink (N=2,871) proteins in the MESA1 test set (N=1,332).

**Table S6.**
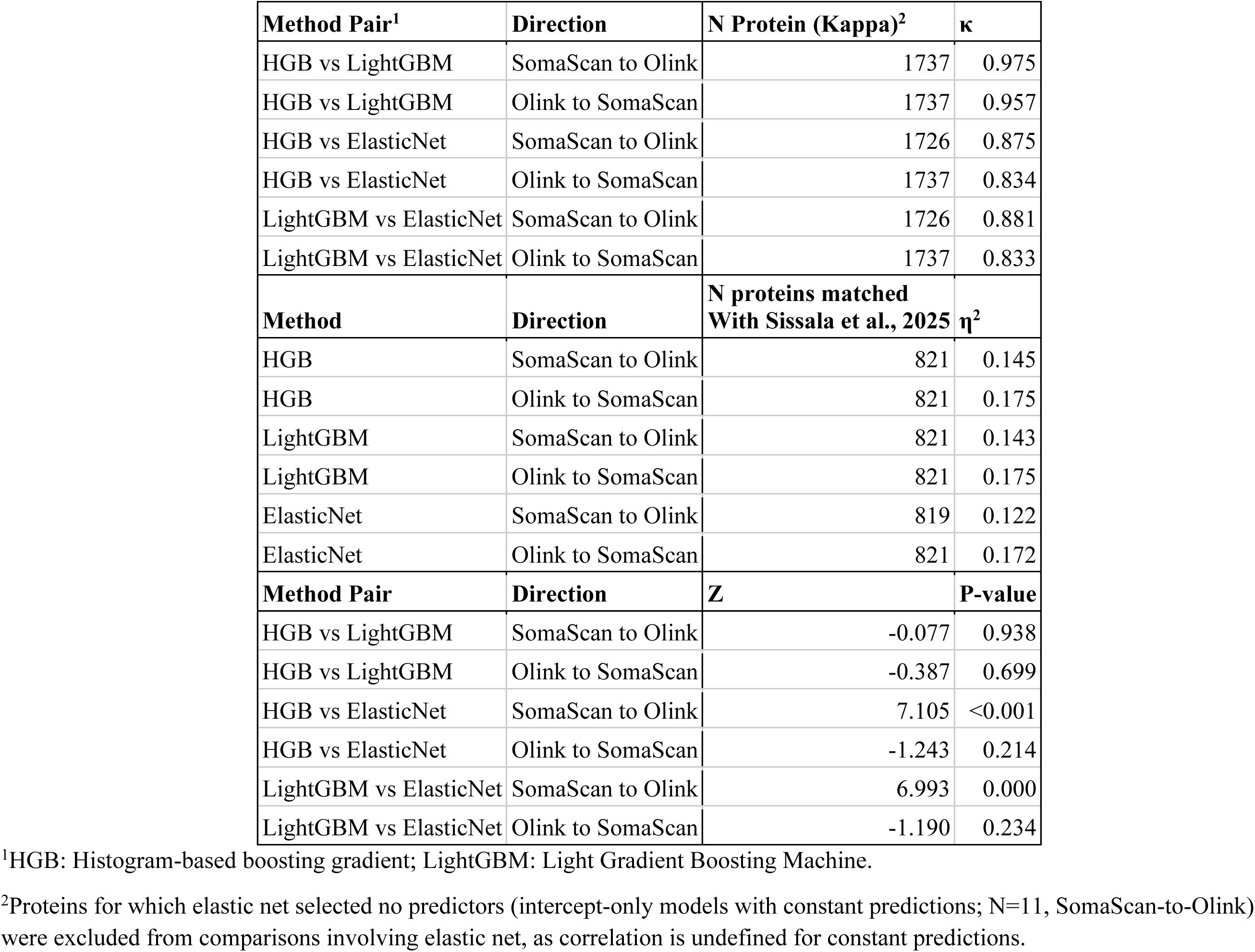
Benchmarking HGB against LightGBM and ElasticNet.

**Table S7.**
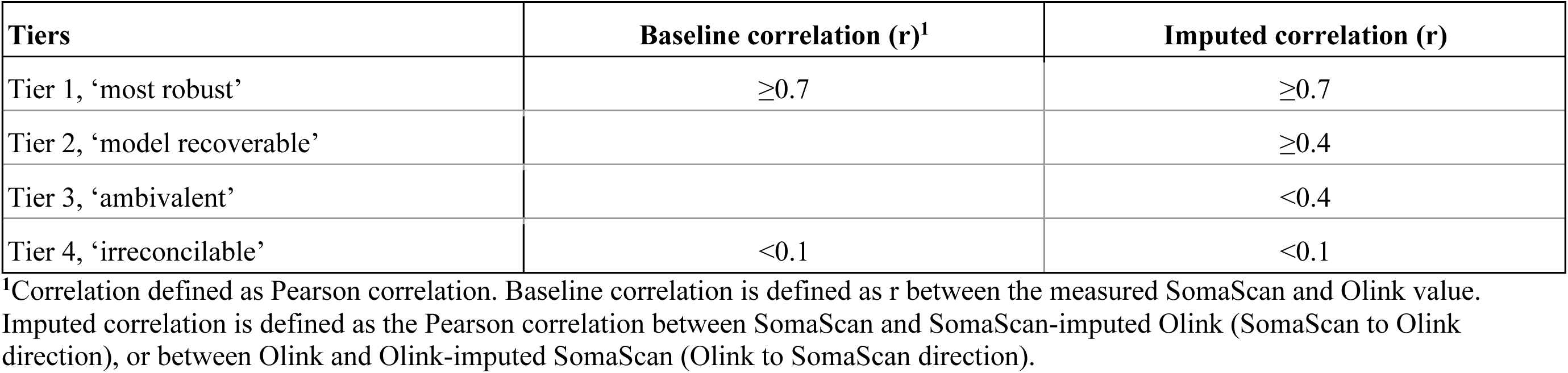
Protein Tier membership thresholds.

**Table S8.**
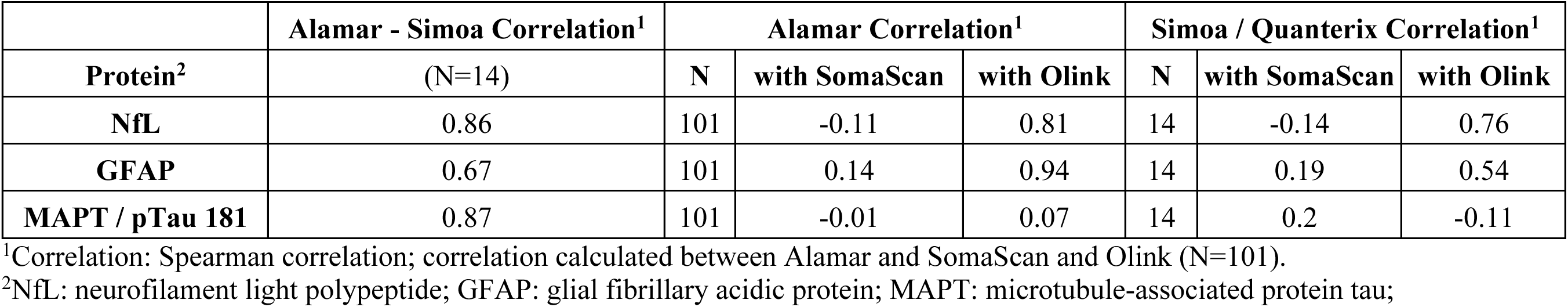
Alamar validation MAPT, GFAP, and NfL in the Atherosclerosis Risk in Communities Study.

**Table S9.**
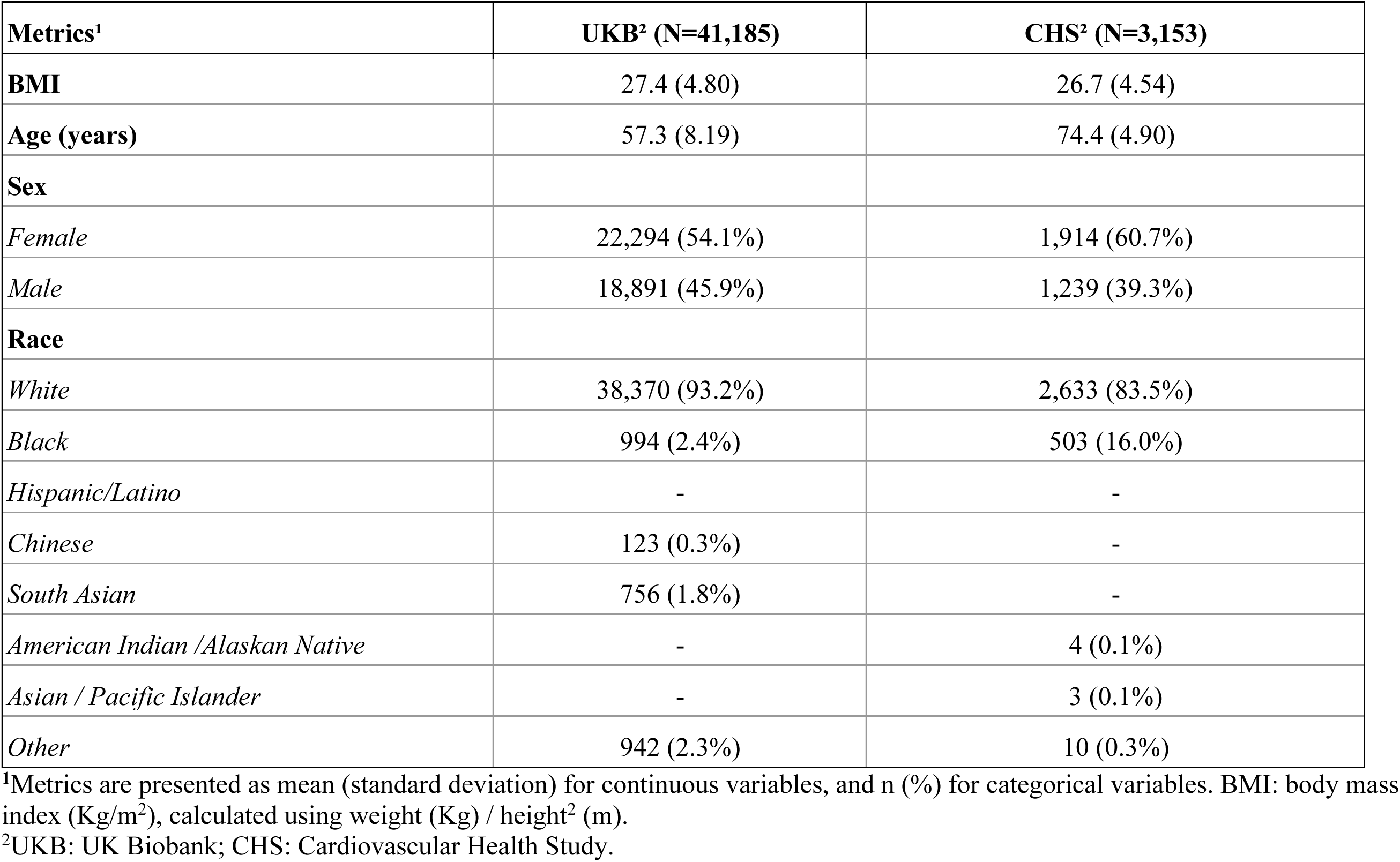
Demographic characteristics of BMI analytical cohorts.

**Table S10.**
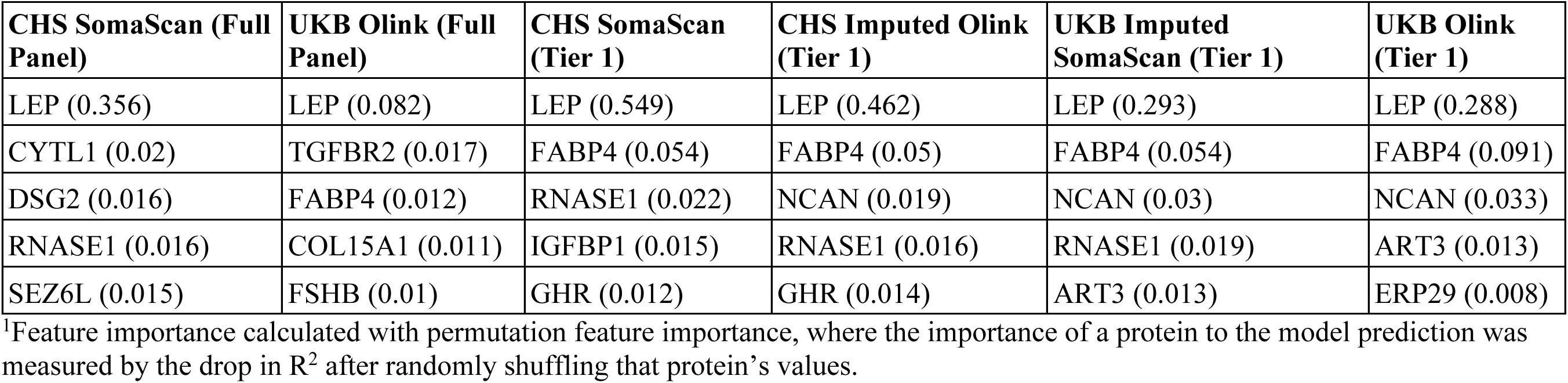
Application 1 - feature importance1 of proteins in BMI prediction.

**Table S11.**
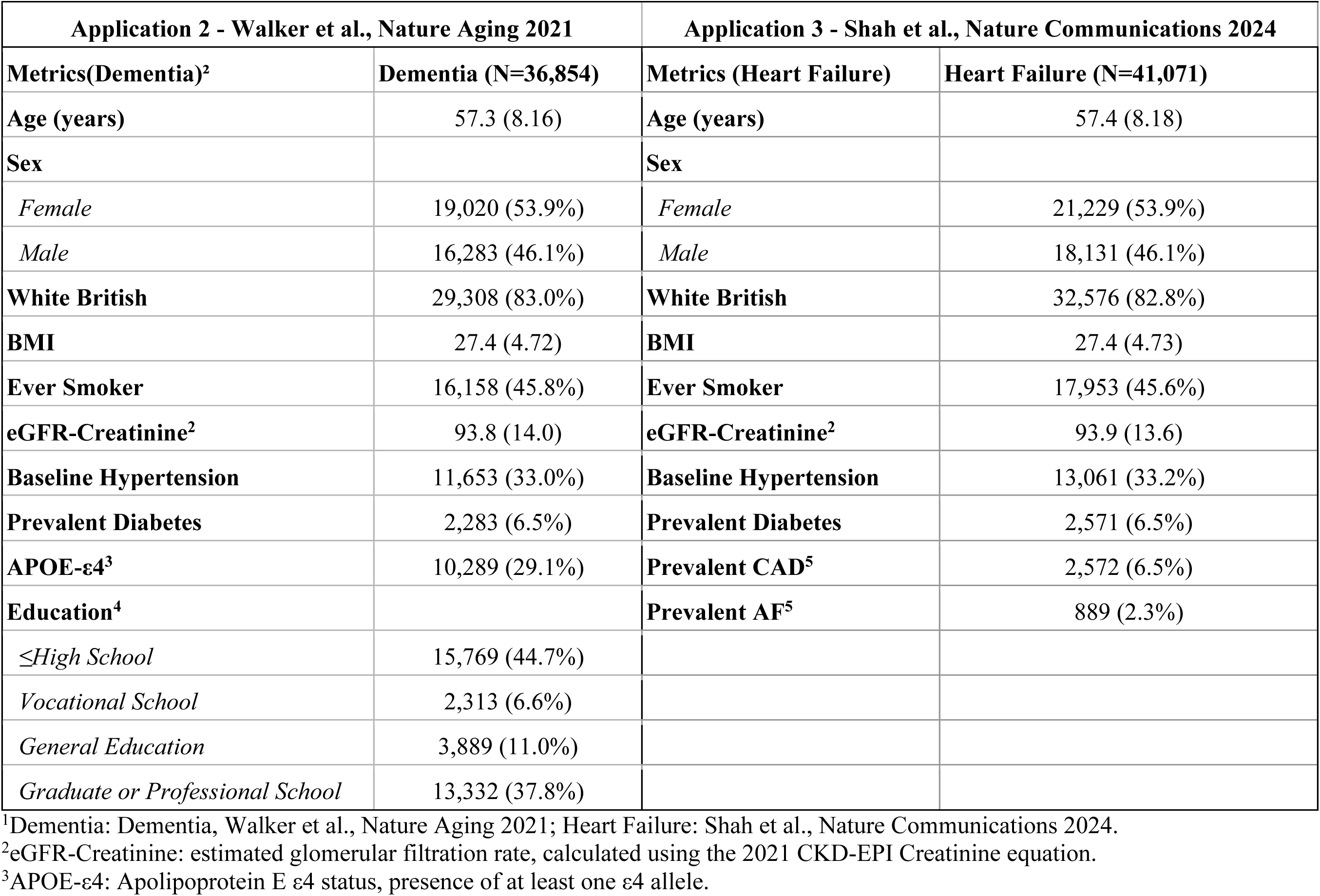
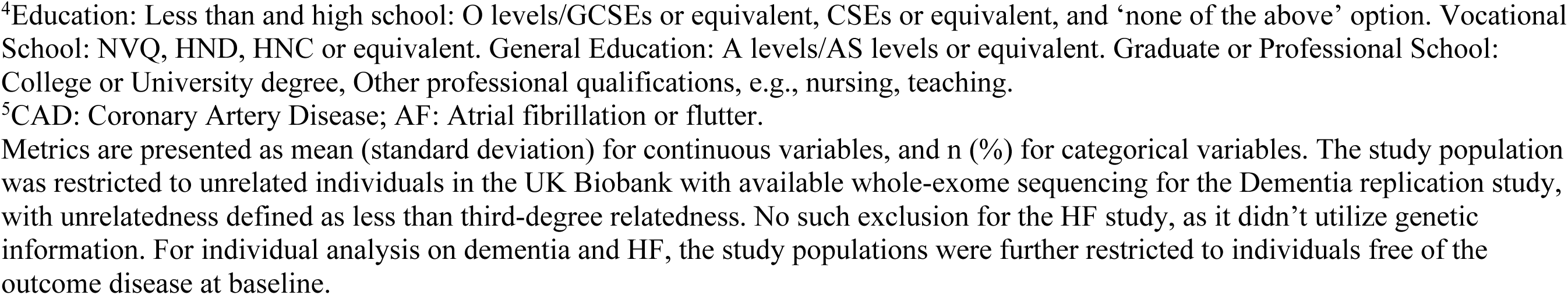
Characteristics of the study population in the UK Biobank in replication studies.

**Table S12.**
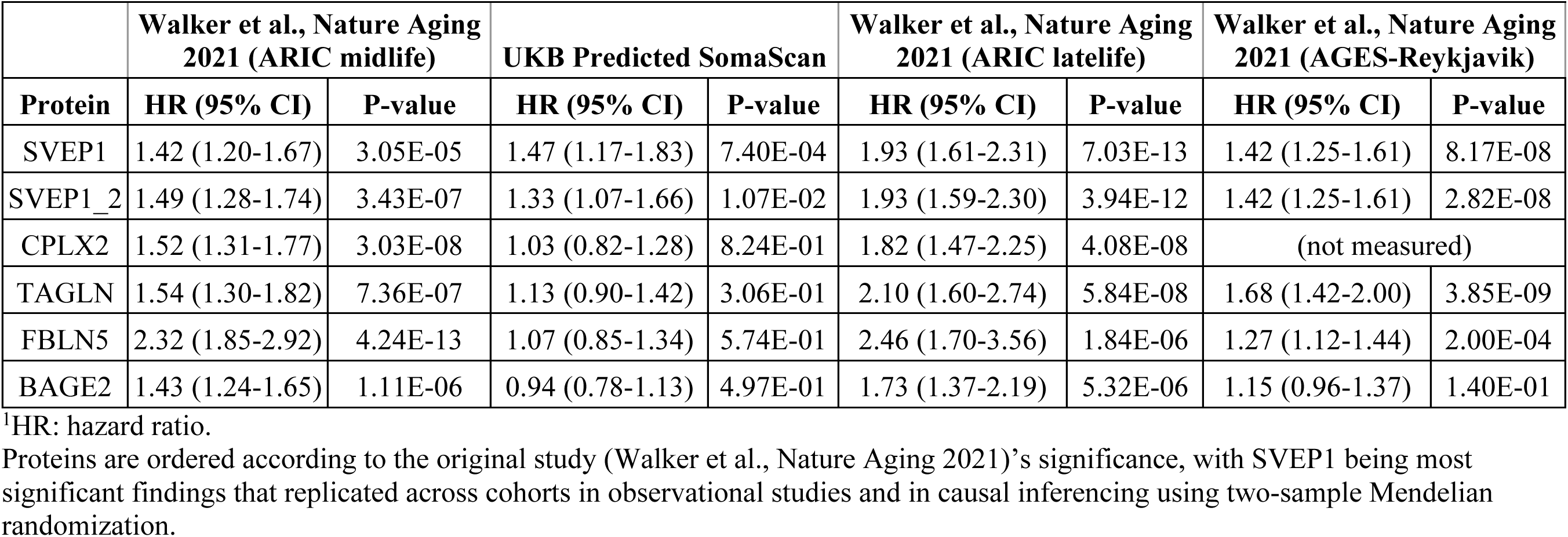
Application 2 - Replication Results of Walker et al., Nature Aging 2021.

**Table S13.**
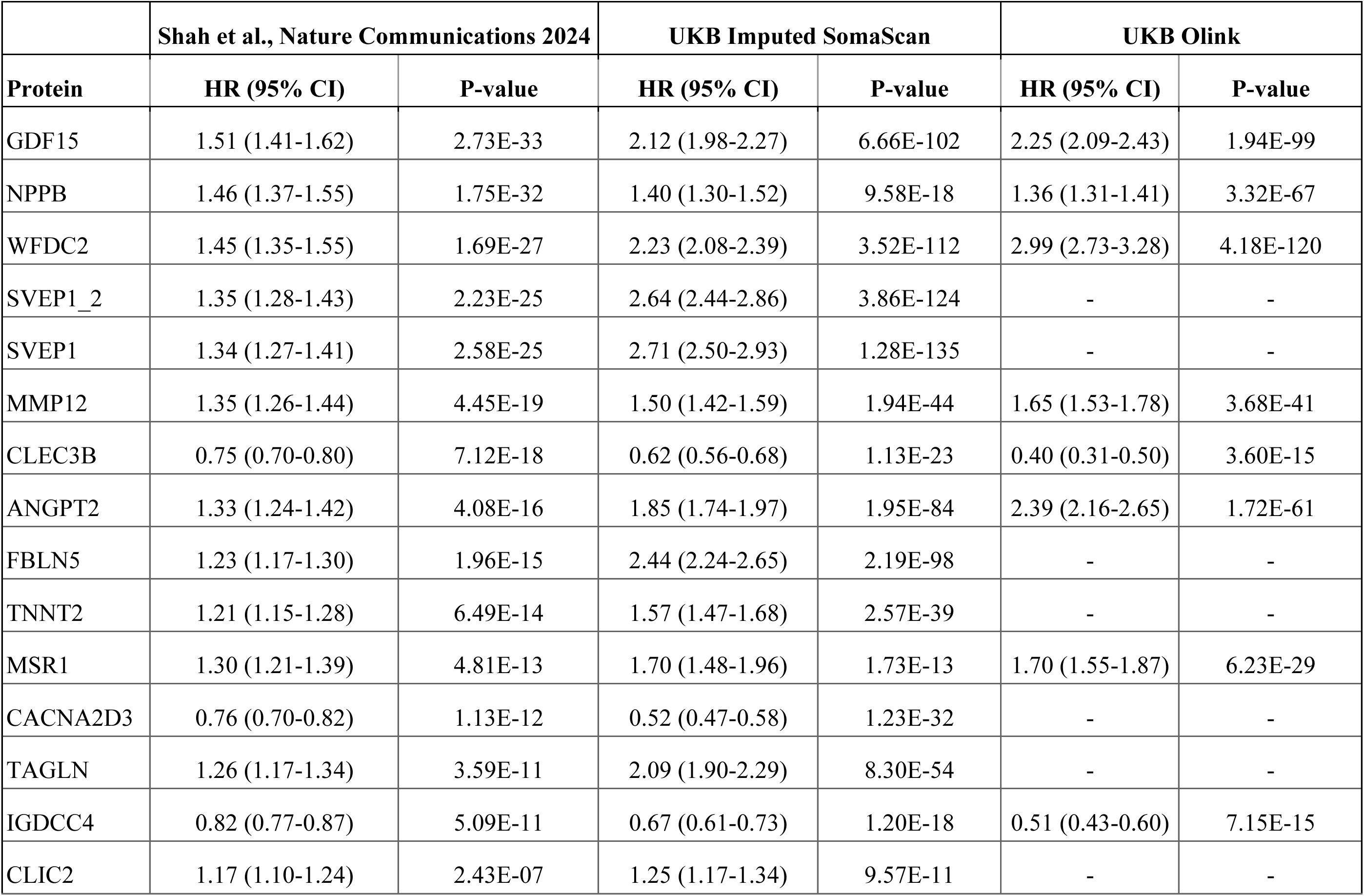
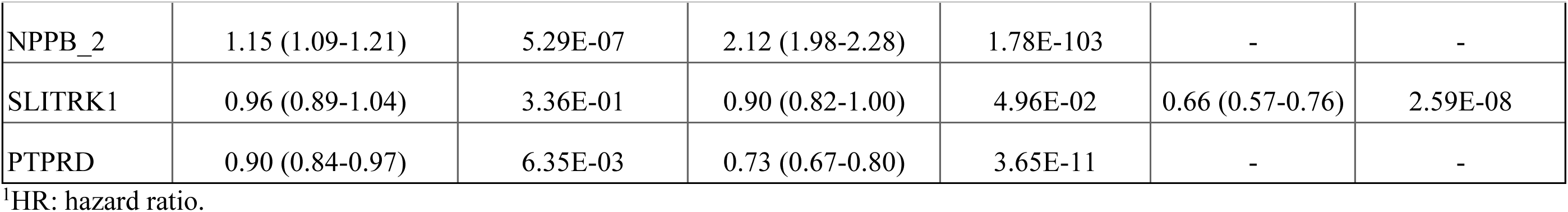
Replication Results of Application 3 - Shah et al., Nature Communications 2024.

**Table S14.**
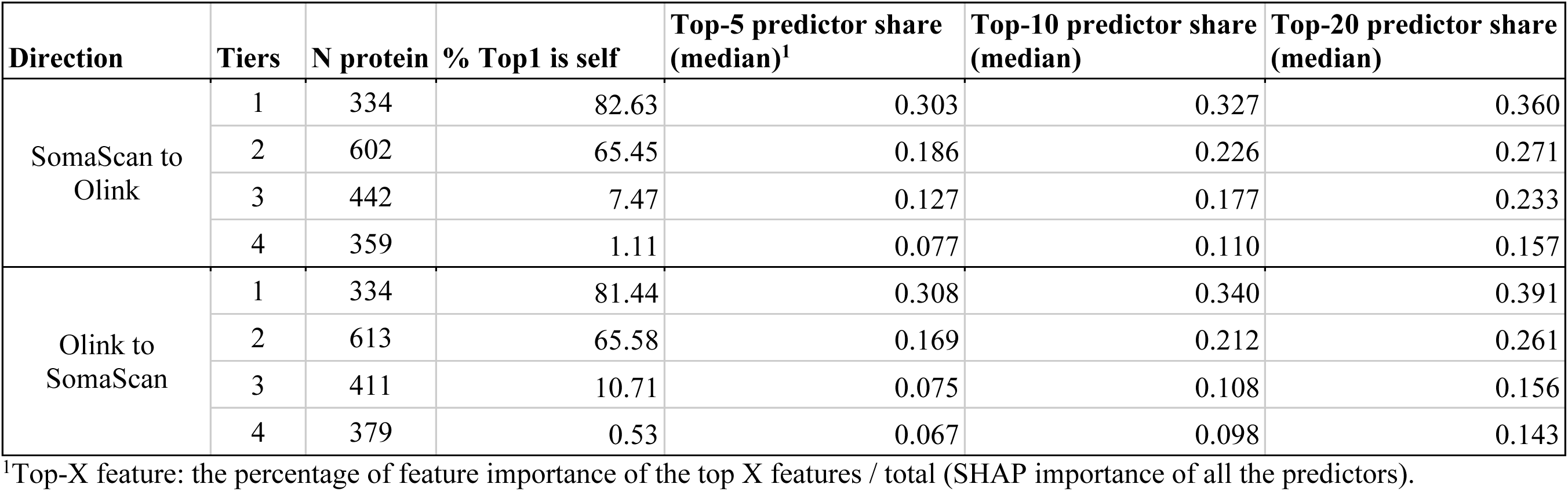
Summary of SHAP-based feature importance in the overlapping proteins (N=1,737) in MESA test set (N=1,332)

**Table S15.**
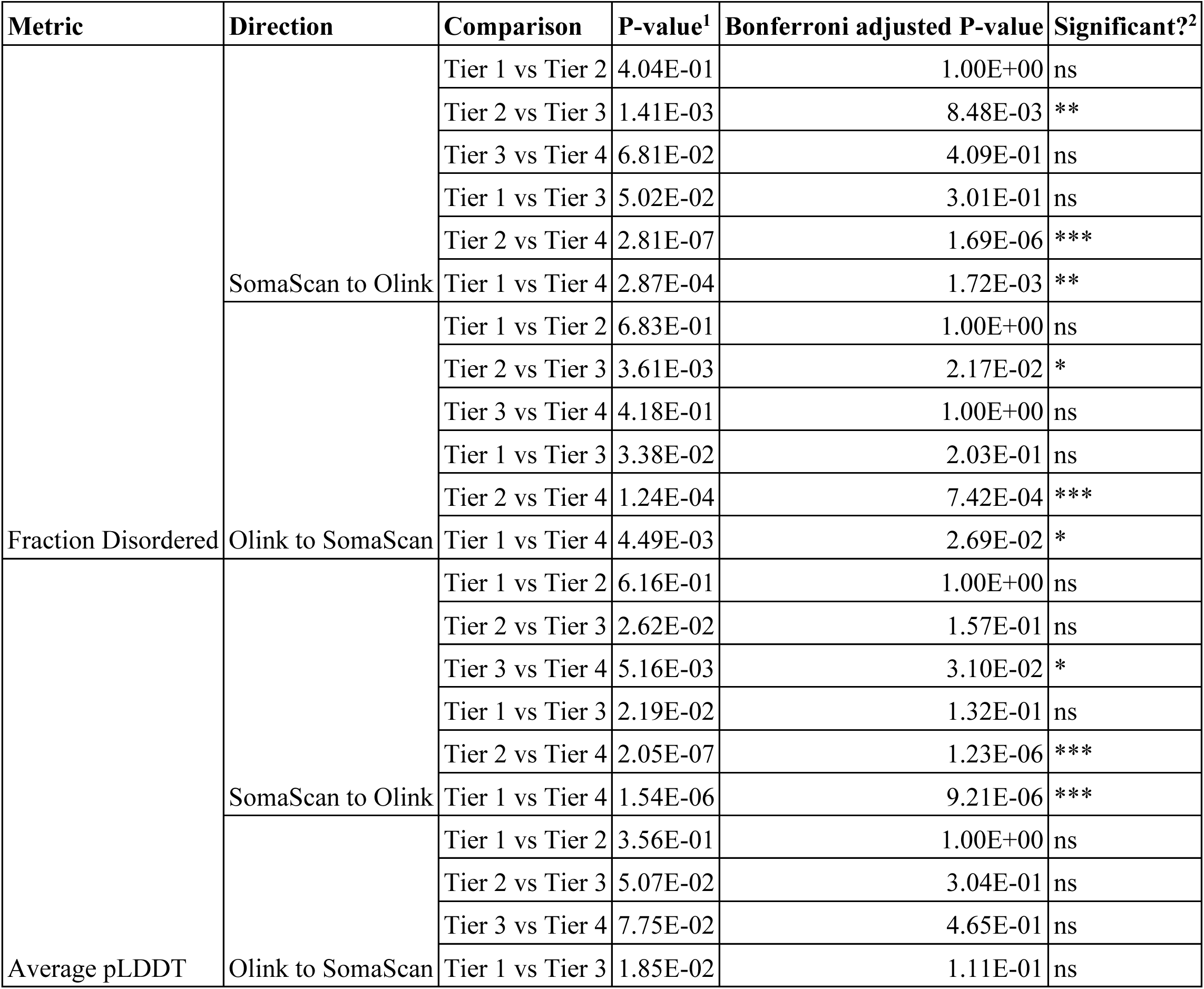
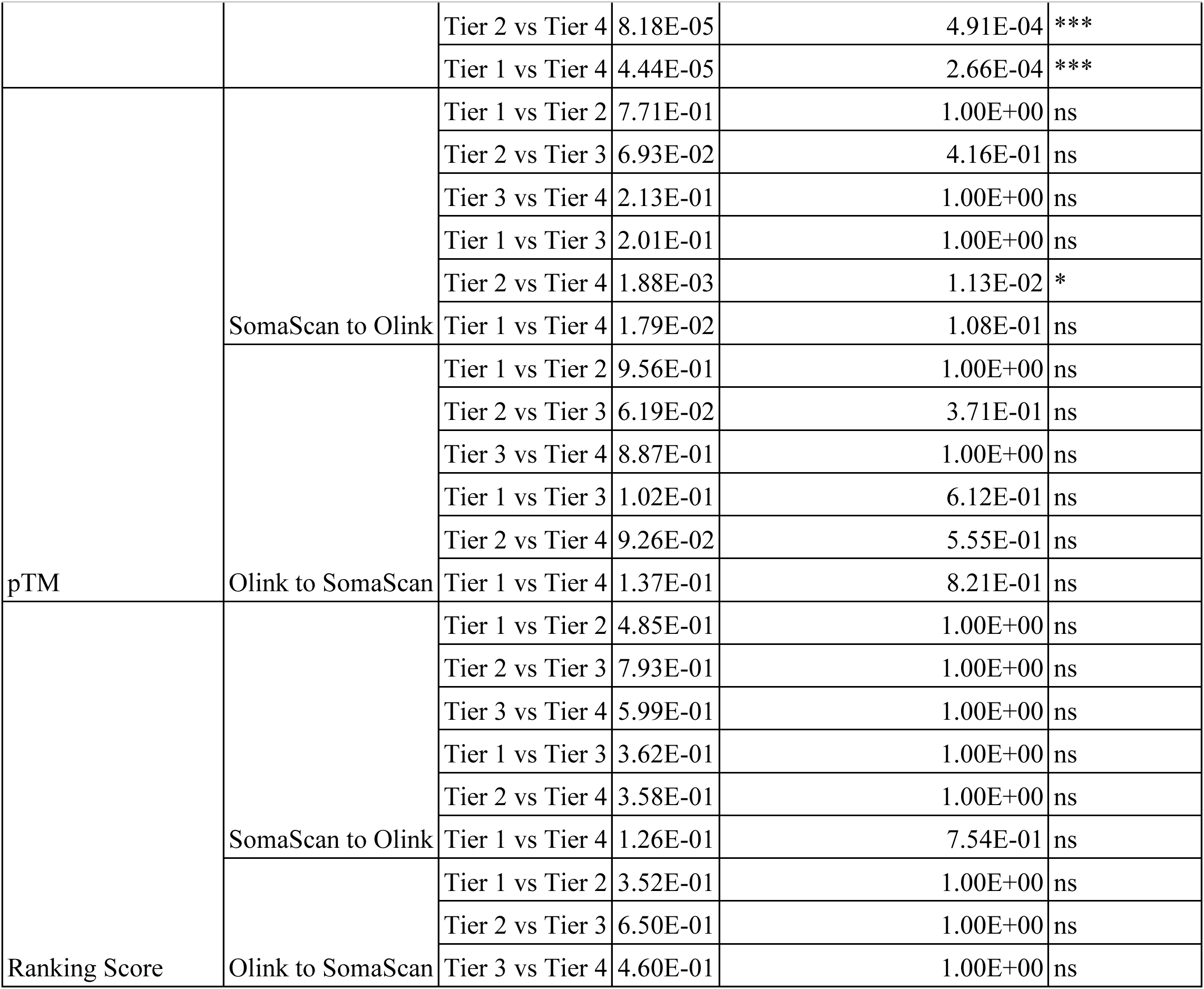
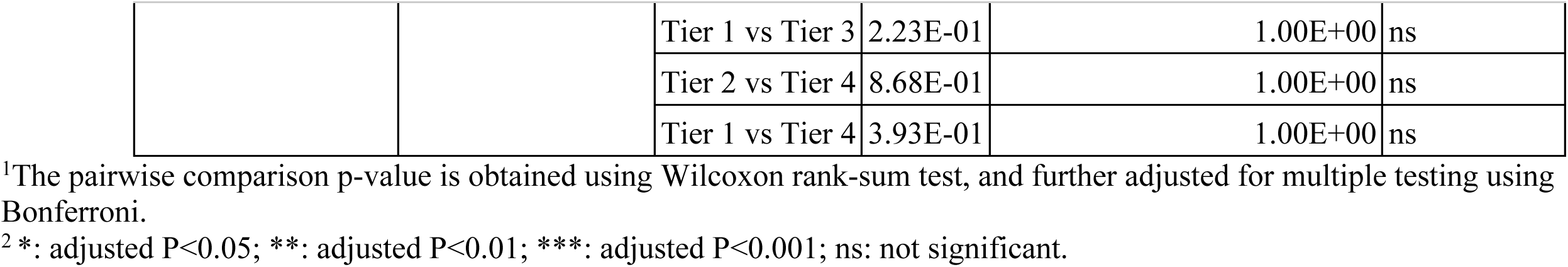
Direction- and pairwise-comparison for all Alphafold 3 metrics.

| Metric | Direction | Comparison | P-value<sup>1</sup> | Bonferroni adjusted P-value | Significant?<sup>2</sup> |
|---|---|---|---|---|---|
| Fraction Disordered | SomaScan to Olink | Tier 1 vs Tier 2 | 4.04E-01 | 1.00E+00 | ns |
|  |  | Tier 2 vs Tier 3 | 1.41E-03 | 8.48E-03 | ** |
|  |  | Tier 3 vs Tier 4 | 6.81E-02 | 4.09E-01 | ns |
|  |  | Tier 1 vs Tier 3 | 5.02E-02 | 3.01E-01 | ns |
|  |  | Tier 2 vs Tier 4 | 2.81E-07 | 1.69E-06 | *** |
|  |  | Tier 1 vs Tier 4 | 2.87E-04 | 1.72E-03 | ** |
|  | Olink to SomaScan | Tier 1 vs Tier 2 | 6.83E-01 | 1.00E+00 | ns |
|  |  | Tier 2 vs Tier 3 | 3.61E-03 | 2.17E-02 | * |
|  |  | Tier 3 vs Tier 4 | 4.18E-01 | 1.00E+00 | ns |
|  |  | Tier 1 vs Tier 3 | 3.38E-02 | 2.03E-01 | ns |
|  |  | Tier 2 vs Tier 4 | 1.24E-04 | 7.42E-04 | *** |
|  |  | Tier 1 vs Tier 4 | 4.49E-03 | 2.69E-02 | * |
| Average pLDDT | SomaScan to Olink | Tier 1 vs Tier 2 | 6.16E-01 | 1.00E+00 | ns |
|  |  | Tier 2 vs Tier 3 | 2.62E-02 | 1.57E-01 | ns |
|  |  | Tier 3 vs Tier 4 | 5.16E-03 | 3.10E-02 | * |
|  |  | Tier 1 vs Tier 3 | 2.19E-02 | 1.32E-01 | ns |
|  |  | Tier 2 vs Tier 4 | 2.05E-07 | 1.23E-06 | *** |
|  |  | Tier 1 vs Tier 4 | 1.54E-06 | 9.21E-06 | *** |
|  | Olink to SomaScan | Tier 1 vs Tier 2 | 3.56E-01 | 1.00E+00 | ns |
|  |  | Tier 2 vs Tier 3 | 5.07E-02 | 3.04E-01 | ns |
|  |  | Tier 3 vs Tier 4 | 7.75E-02 | 4.65E-01 | ns |
|  |  | Tier 1 vs Tier 3 | 1.85E-02 | 1.11E-01 | ns |
|  |  | Tier 2 vs Tier 4 | 8.18E-05 | 4.91E-04 | *** |
|  |  | Tier 1 vs Tier 4 | 4.44E-05 | 2.66E-04 | *** |
| pTM | SomaScan to Olink | Tier 1 vs Tier 2 | 7.71E-01 | 1.00E+00 | ns |
|  |  | Tier 2 vs Tier 3 | 6.93E-02 | 4.16E-01 | ns |
|  |  | Tier 3 vs Tier 4 | 2.13E-01 | 1.00E+00 | ns |
|  |  | Tier 1 vs Tier 3 | 2.01E-01 | 1.00E+00 | ns |
|  |  | Tier 2 vs Tier 4 | 1.88E-03 | 1.13E-02 | * |
|  |  | Tier 1 vs Tier 4 | 1.79E-02 | 1.08E-01 | ns |
|  | Olink to SomaScan | Tier 1 vs Tier 2 | 9.56E-01 | 1.00E+00 | ns |
|  |  | Tier 2 vs Tier 3 | 6.19E-02 | 3.71E-01 | ns |
|  |  | Tier 3 vs Tier 4 | 8.87E-01 | 1.00E+00 | ns |
|  |  | Tier 1 vs Tier 3 | 1.02E-01 | 6.12E-01 | ns |
|  |  | Tier 2 vs Tier 4 | 9.26E-02 | 5.55E-01 | ns |
|  |  | Tier 1 vs Tier 4 | 1.37E-01 | 8.21E-01 | ns |
| Ranking Score | SomaScan to Olink | Tier 1 vs Tier 2 | 4.85E-01 | 1.00E+00 | ns |
|  |  | Tier 2 vs Tier 3 | 7.93E-01 | 1.00E+00 | ns |
|  |  | Tier 3 vs Tier 4 | 5.99E-01 | 1.00E+00 | ns |
|  |  | Tier 1 vs Tier 3 | 3.62E-01 | 1.00E+00 | ns |
|  |  | Tier 2 vs Tier 4 | 3.58E-01 | 1.00E+00 | ns |
|  |  | Tier 1 vs Tier 4 | 1.26E-01 | 7.54E-01 | ns |
|  | Olink to SomaScan | Tier 1 vs Tier 2 | 3.52E-01 | 1.00E+00 | ns |
|  |  | Tier 2 vs Tier 3 | 6.50E-01 | 1.00E+00 | ns |
|  |  | Tier 3 vs Tier 4 | 4.60E-01 | 1.00E+00 | ns |
|  |  | Tier 1 vs Tier 3 | 2.23E-01 | 1.00E+00 | ns |
|  |  | Tier 2 vs Tier 4 | 8.68E-01 | 1.00E+00 | ns |
|  |  | Tier 1 vs Tier 4 | 3.93E-01 | 1.00E+00 | ns |

<sup>1</sup>The pairwise comparison p-value is obtained using Wilcoxon rank-sum test, and further adjusted for multiple testing using Bonferroni.
<sup>2</sup> \*: adjusted P<0.05; \*\*: adjusted P<0.01; \*\*\*: adjusted P<0.001; ns: not significant.

