## Supplementary tables and figures for "Machine learning cross-platform proteomic imputation enables protein quality scoring and replication of epidemiological associations"

### Supplementary Figures

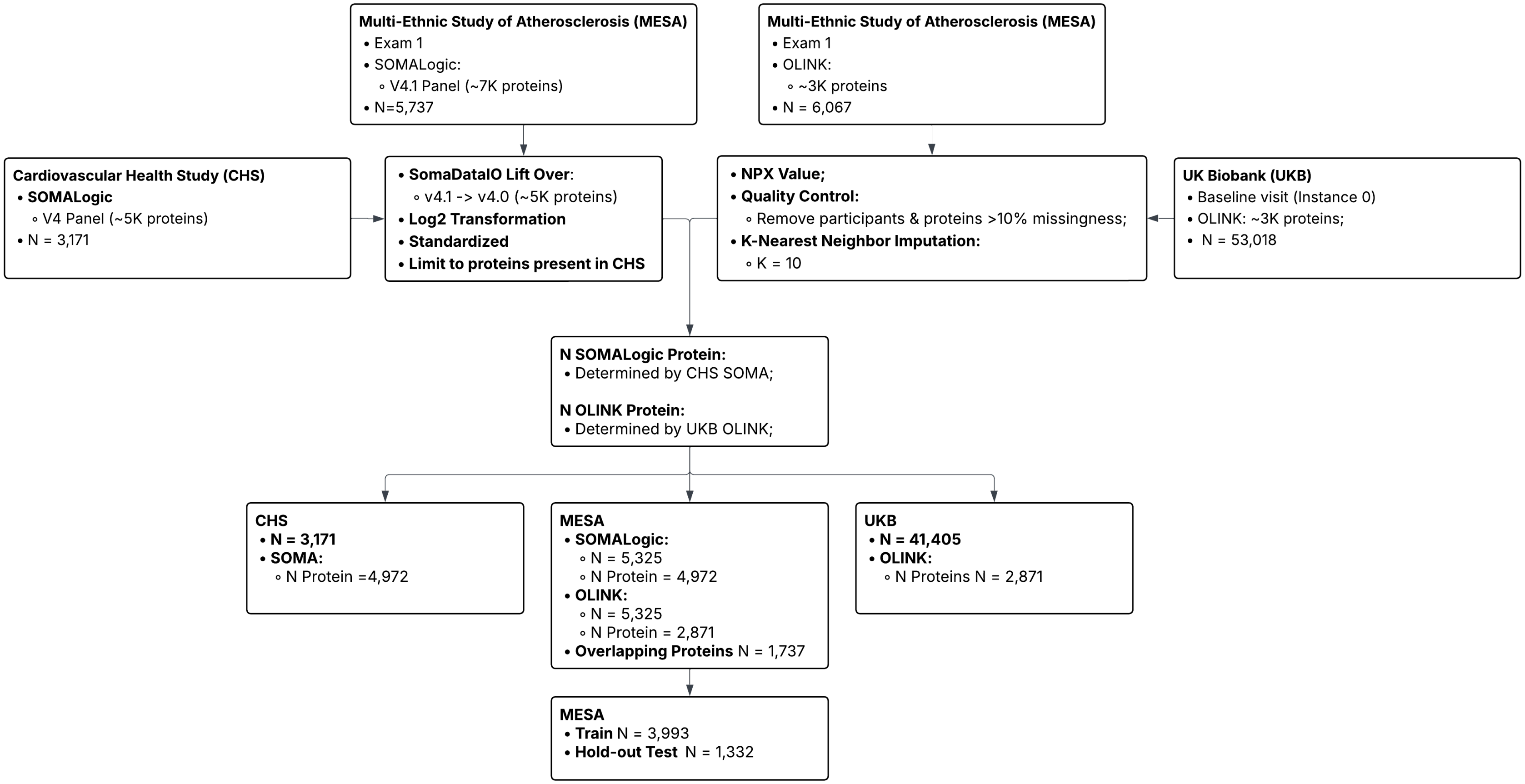


**Supplementary Figure 1. Consort diagram of sample and protein quality control in three cohorts: MESA, UKB and CHS.** MESA: Multi-Ethnic Study of Atherosclerosis; UKB: UK Biobank; CHS: Cardiovascular Health Study.

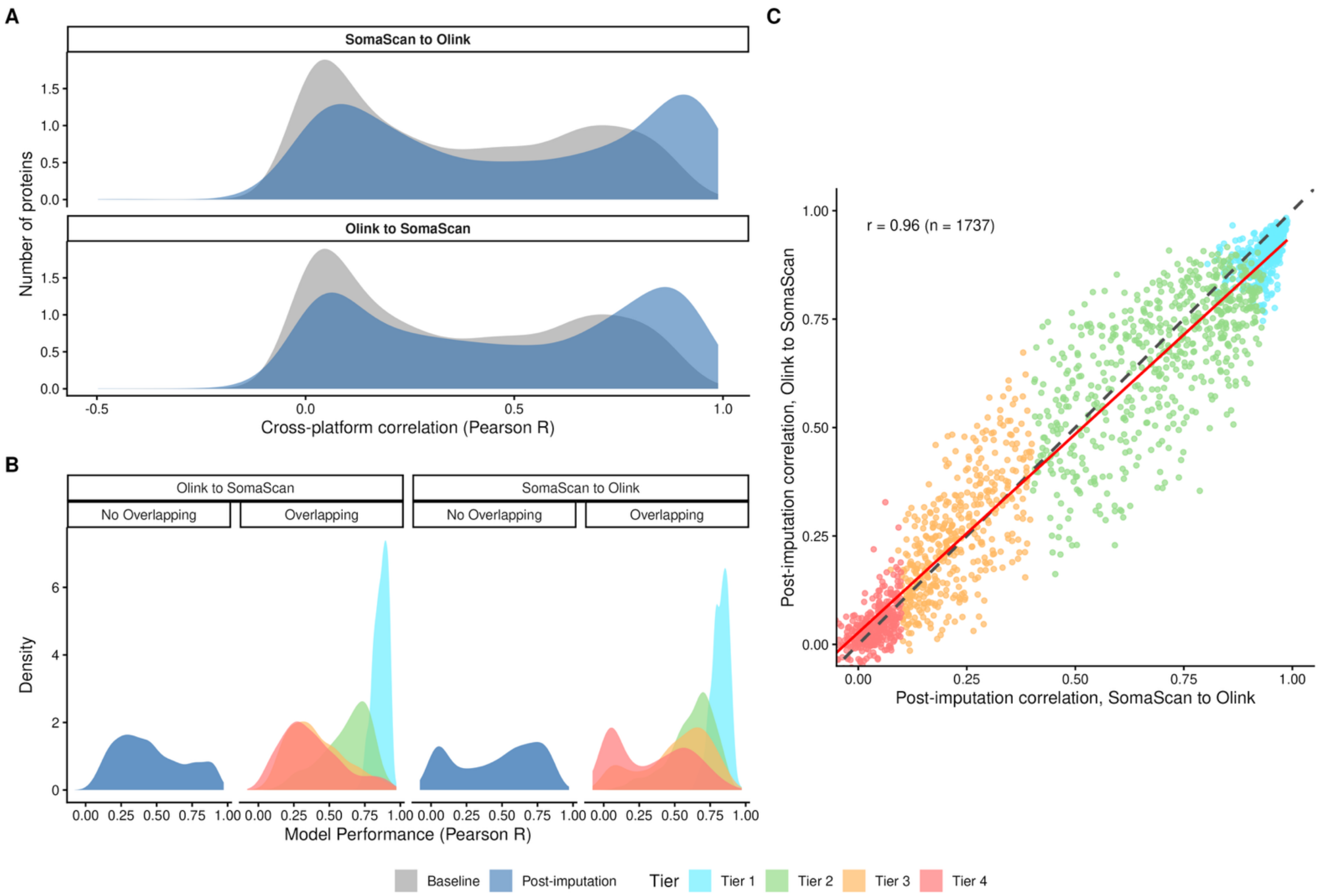


**Supplementary Figure 2. Shift in cross-platform correlation and model predictive performance (Pearson correlation) in the MESA held-out test set (N=1,332).**
**Panel A. Density plot of baseline and post-imputation crossplatform correlation stratified by imputation direction measured by Pearson correlation; Panel B. Density plot of model predictive performance stratified by imputation direction and overlapping status of the full panel proteins (SomaScan N=4,972; Olink N=2,871), colored by assigned reliability tiers (overlapping proteins only); Panel C. Concordance between the post-imputation correlation of overlapping proteins (N=1,737) of two imputation direction.** Each dot represents one protein. The black dashed line is x=y, meaning the post-imputation correlation

is identical in both directions. The proteins/dots above the dashed line have higher post-imputation correlation in Olink to SomaScan direction, vice versa. The red line represents the fitted line using y ~ x geom_smooth function, and the concordance between the two post-imputation correlation is measured by Pearson correlation. Tier 1 and Tier 4 proteins have more similar post-imputation correlation between 2 imputation directions compared to Tier 2 and 3 proteins.
MESA: Multi-Ethnic Study of Atherosclerosis.

A

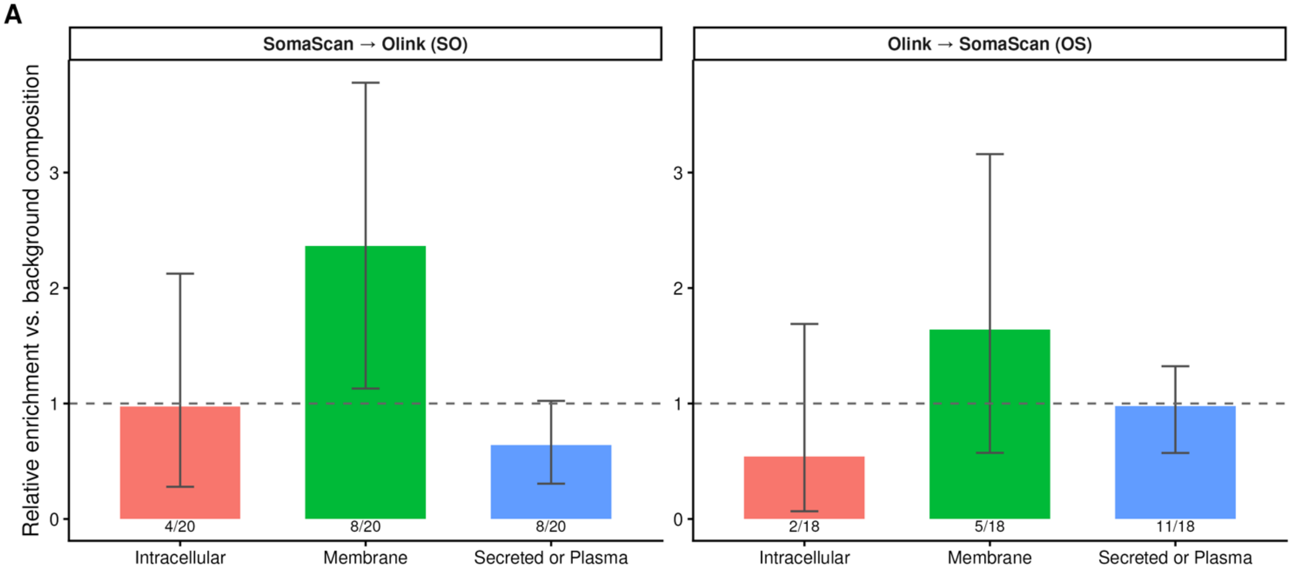


B

| Direction | Class | N Negative / N Total | Relative Enrichment | 95% CI | P adj. | Sig ? |
|---|---|---|---|---|---|---|
| SomaScan → Olink (SO) | Intracellular | 4 / 20 | 0.97 | 0.28 - 2.12 | 1.000 | |
| SomaScan → Olink (SO) | Membrane | 8 / 20 | 2.36 | 1.13 - 3.78 | 0.037 | * |
| SomaScan → Olink (SO) | Secreted or Plasma | 8 / 20 | 0.64 | 0.31 - 1.02 | 0.092 | |
| Olink → SomaScan (OS) | Intracellular | 2 / 18 | 0.54 | 0.07 - 1.69 | 0.836 | |
| Olink → SomaScan (OS) | Membrane | 5 / 18 | 1.64 | 0.57 - 3.16 | 0.636 | |
| Olink → SomaScan (OS) | Secreted or Plasma | 11 / 18 | 0.98 | 0.57 - 1.32 | 1.000 | |

**Supplementary Figure 3. Proteins ($N_{SO}$=20, $N_{OS}$=18) with meaningful negative gain (gain < -0.05) by localization.**
**Panel A.** Relative enrichment in each direction is calculated as: Relative enrichment = (proportion of the negative-gain set in a class) / (proportion of all 1,737 overlapping proteins in that class). Dashed line: no enrichment (ratio = 1). 95% Confidence intervals were calculated using exact Binomial test.
*: adjusted P<0.05, **: adjusted P<0.01, *** adjusted P<0.001

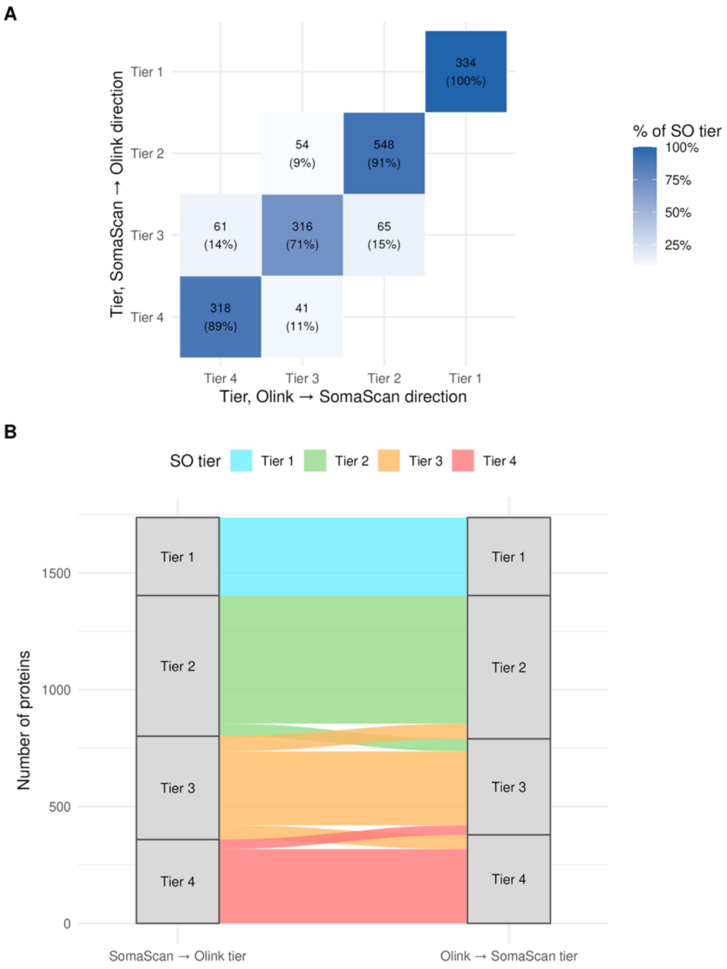


C

*Tier 3 → Tier 2 ($\chi^2$ p = 5.17e-01)*

| Location | Obs n | Obs % | Bkgd % | Enr. ratio |
|---|---|---|---|---|
| Intracellular | 17 | 26% | 21% | 1.27 |
| Membrane | 11 | 17% | 17% | 1.00 |
| Secreted or Plasma | 37 | 57% | 63% | 0.91 |

D

*Tier 3 → Tier 4 ($\chi^2$ p = 3.17e-02)*

| Location | Obs n | Obs % | Bkgd % | Enr. ratio |
|---|---|---|---|---|
| Intracellular | 20 | 33% | 21% | 1.60 |
| Membrane | 12 | 20% | 17% | 1.16 |
| Secreted or Plasma | 29 | 48% | 63% | 0.76 |

E

*Tier 4 → Tier 3 ($\chi^2$ p = 1.16e-02)*

| Location | Obs n | Obs % | Bkgd % | Enr. ratio |
|---|---|---|---|---|
| Intracellular | 8 | 20% | 21% | 0.95 |
| Membrane | 14 | 34% | 17% | 2.02 |
| Secreted or Plasma | 19 | 46% | 63% | 0.74 |

**Supplementary Figure 4. Shift in tiers by direction and protein localization enrichment for the overlapping proteins (N=1,737). Panel A. Confusion matrix of tier assignments across 2 imputation directions for overlapping proteins.** Cell values show protein count and the row-wise (based on tier assignment in SO direction) percentage. **Panel B. Sankey Diagram of change in tiers between the imputation directions. Panel C-E. Protein localization class (Intracellular, Membrane, Plasma) composition among proteins with tier changes. (C)Tier 3 to 2. (D) Tier 3 to 4, and (E)Tier 4 to 3.** The proportion of proteins within each localization class is compared with their background frequency using Chisq test to determine if one localization class was enriched for a specific switch in tiers.

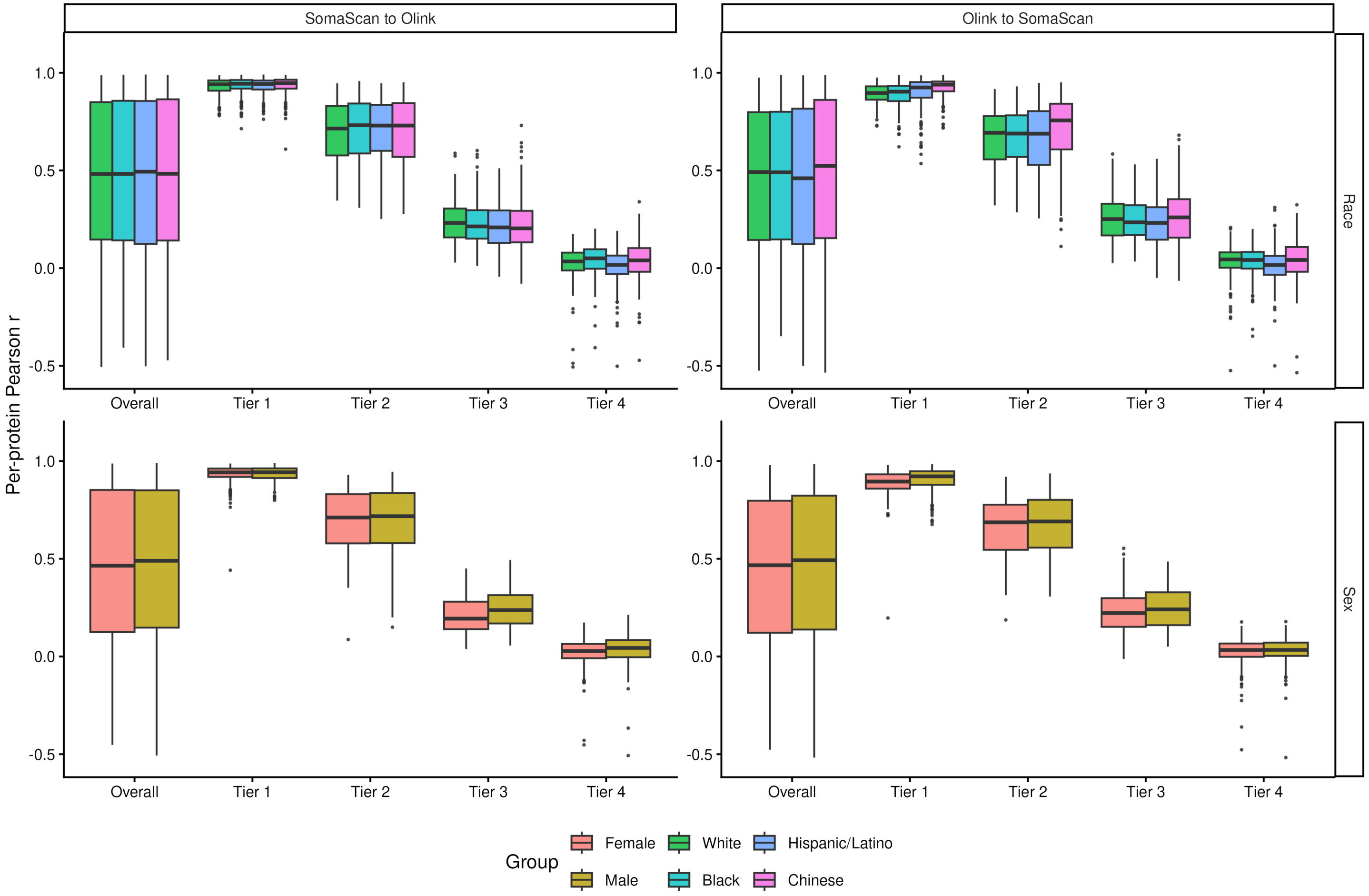


**Supplementary Figure 5. Post-imputation Pearson r of overlapping proteins(N=1,737) by MESA subgroups in test set (N=1,332).**

Boxplot for post-imputation correlation for each tier for the full test set and stratified by demographic subgroups is used. Demographic subgroups of sex (Male, Female) and race/ethnicity (White, Black, Hispanic/Latino, Chinese) were used.

MESA: Multi-Ethnic Study of Atherosclerosis.

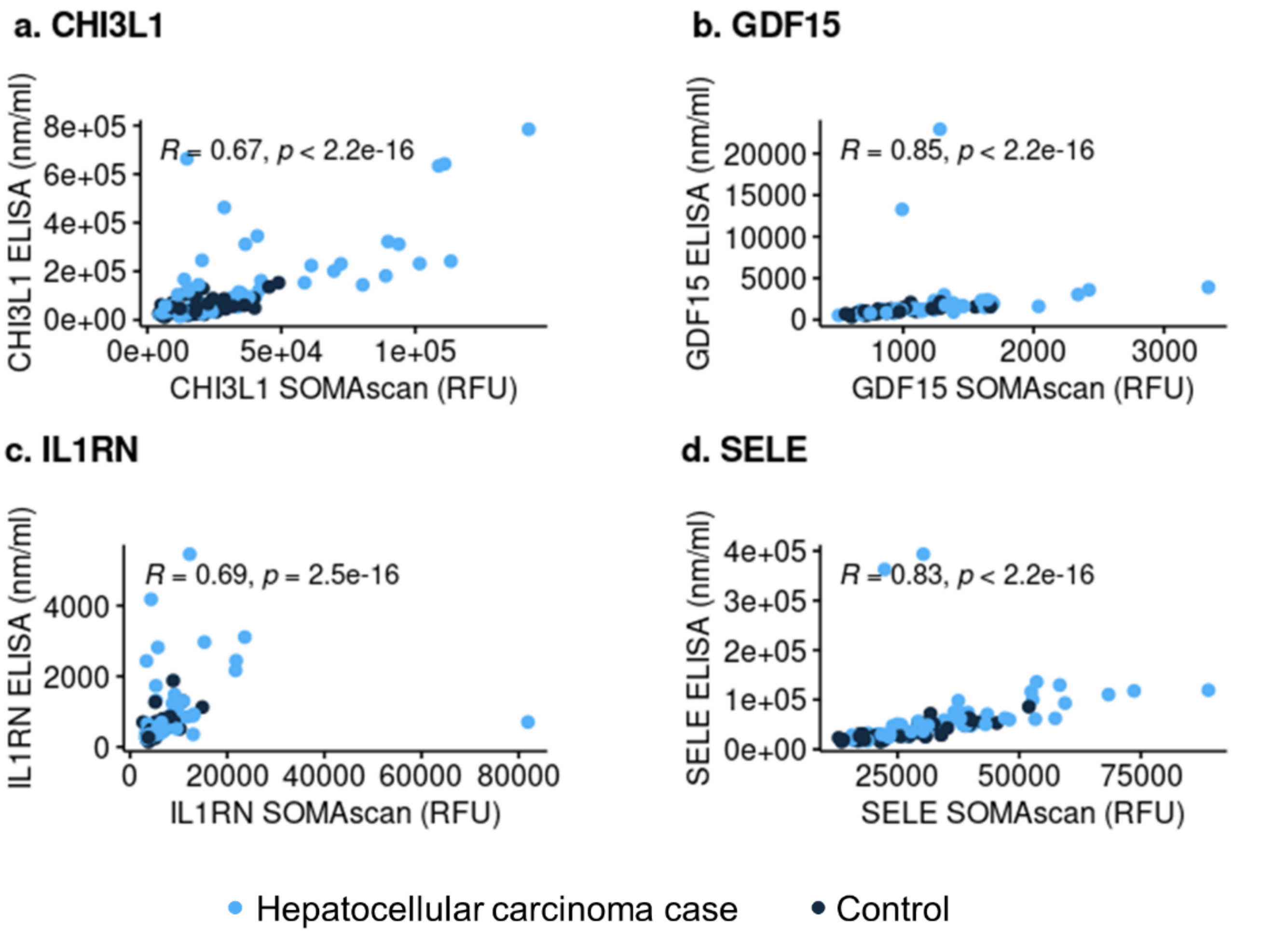


**Supplementary Figure 6. Spearman correlations between the SomaScan and ELISA protein levels in the Nurses' Health Study and Health Professionals Follow-up Study (N=54)**
Re-use with permission from lead author Xinyuan Zhang. Figure also published as Supplemental Figure 2 in Zhang et al., Journal of the National Cancer Institute 2024, doi: 10.1093/jnci/djae079.

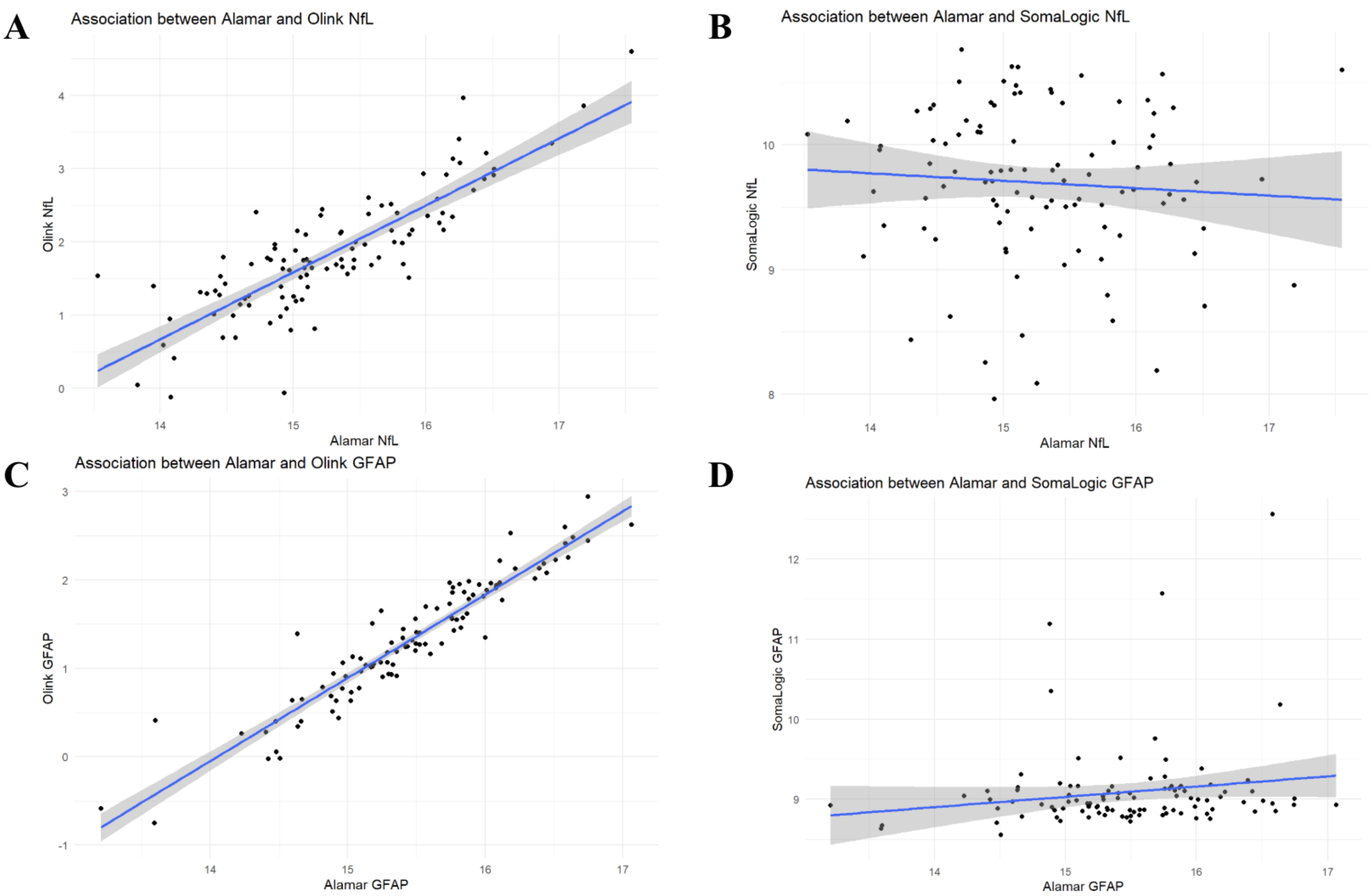


**Supplementary Figure 7. Spearman Correlation between Alamar, SomaScan and Olink measurements in Atherosclerosis Risk in Communities (N=101).**

Correlation calculated between Alamar and SomaScan and Olink (N~101). The Simoa (ELISA based, gold standard) had a high correlation with Alamar assays (>0.9 after excluding 1 outlier)

NfL: neurofilament light polypeptide; GFAP: glial fibrillary acidic protein;

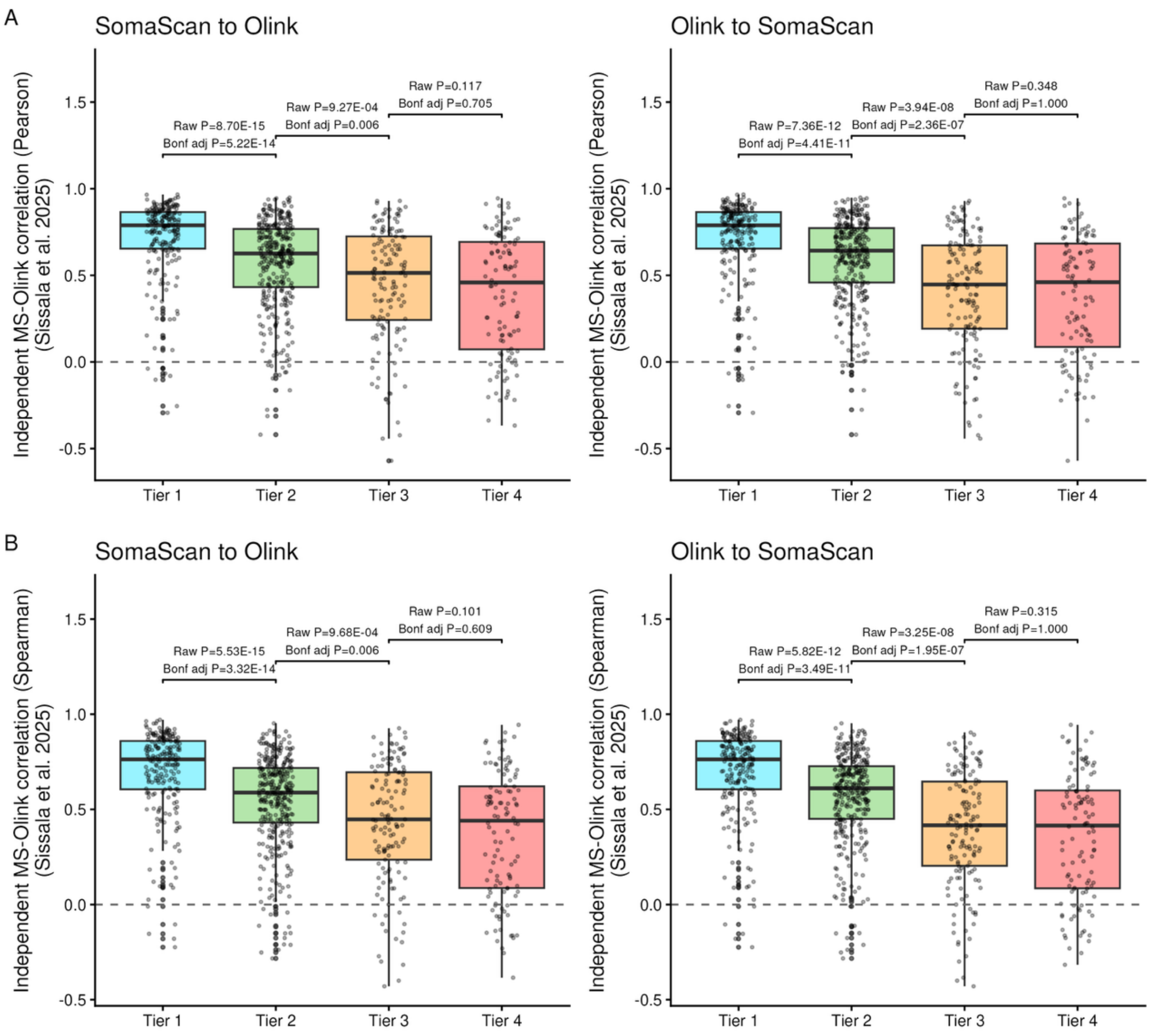


**Supplementary Figure 8. Bidirection tier assignment compared to MS-Olink correlation in Sissala et al., 2025.**
Panel A. Boxplots for distribution of independent MS-Olink correlation (Pearson correlation) from Sissala et al., 2025 stratified by our tier list. Panel B. Boxplots for distribution of independent MS-Olink correlation (Spearman correlation) from Sissala et al., 2025 stratified by our tier list. Dunn's test was used to test the pair-wise MS-Olink correlation between assigned tiers.

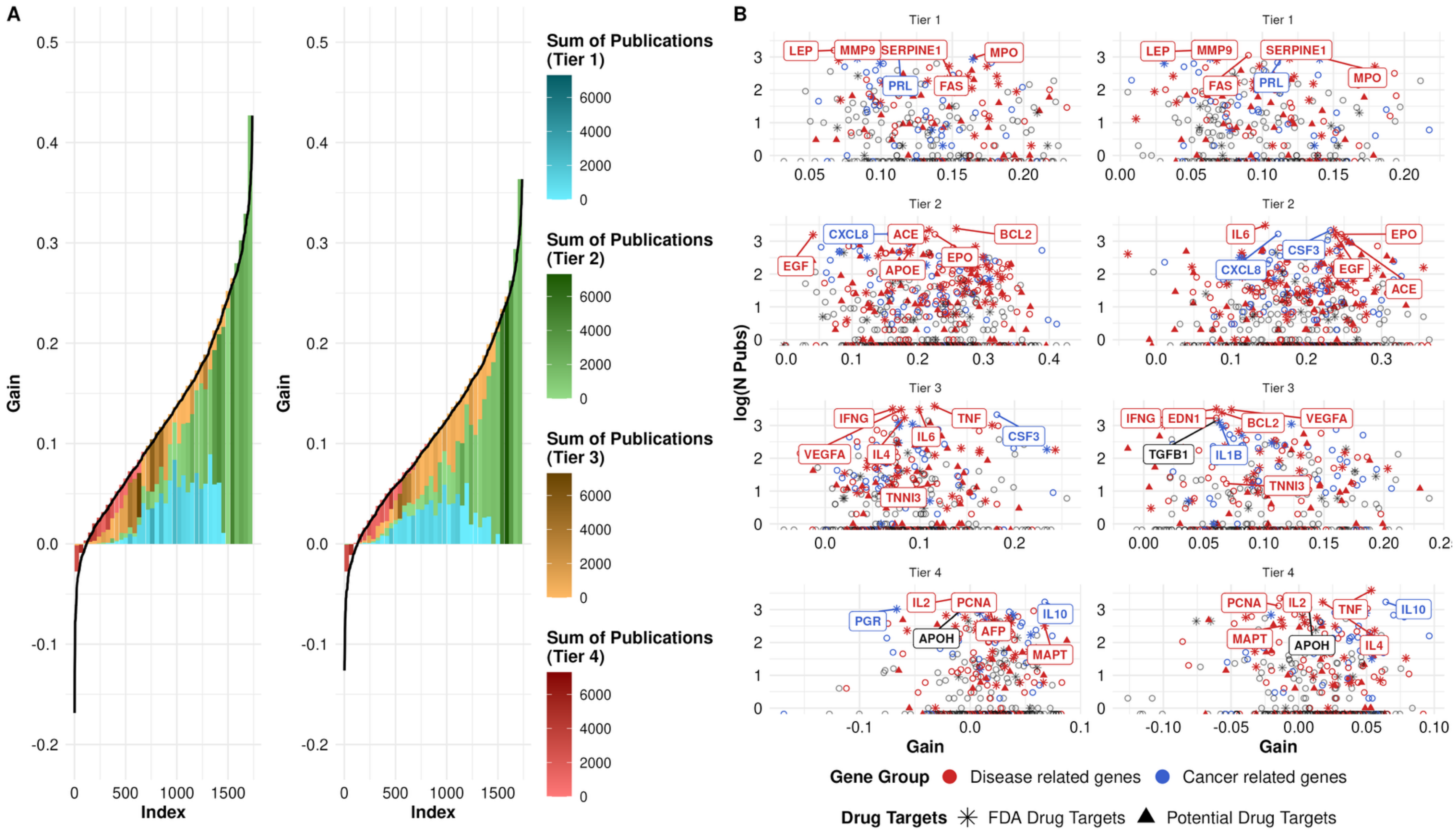


**Supplementary Figure 9. Protein importance of 1,737 overlapping proteins.**
**Panel A. Binned importance by correlation gain (left: SomaScan to Olink; Right: Olink to SomaScan);** The Y-axis illustrates the gain in correlation (post-imputation Pearson correlation minus baseline Pearson correlation) for 1,737 overlapping proteins between the two platforms. X-axis represents the protein index, ranked in ascending order of the gain. All proteins were binned into 220 bins based on their gain for better visual demonstration of importance. Importance is defined by the sum of publications of a given protein through text mining provided by GeneCards. Within individual bins, the size of each color block (each protein tier) is proportional to the proportion of: (number of proteins in that tier / total number of proteins in the bin); the larger the proportion, the more proteins in

the corresponding tier compared to other tiers present in the bin. In each bin, the aggregated number of publications of all proteins grouped by tiers is calculated: the darker the color, the more total publication proteins have and hence more important the proteins are. **Panel B. Scatter plot of log(number of publications) stratified by tiers (left: SomaScan to Olink; right: Olink to SomaScan).** Each point represents one protein, colored by gene group (disease or cancer association), and shaped by druggability (FDA-approved or potential drug target). Y-axis denotes the log(number of publications) obtained using GeneALaCart.

**Supplementary Tables**

**Supplementary Table 1. Demographics of the 3 study cohorts (MESA, UKB, CHS)[1]**

| **Metrics[2]** | **MESA (N=5,325)** | **UKB (N=41,405)** | **CHS (N=3,171)** |
|---|---|---|---|
| **Age (years)** | 62.2 (10.3) | 57.3 (8.19) | 74.4 (4.91) |
| **Sex** | | | |
| *Female* | 2,763 (51.9%) | 22,389 (54.1%) | 1,927 (60.8%) |
| *Male* | 2,562 (48.1%) | 19,016 (45.9%) | 1,244 (39.2%) |
| **Race** | | | |
| *White* | 2,111 (39.6%) | 38,539 (93.1%) | 2,649 (83.5%) |
| *Black* | 1,349 (25.3%) | 1,009 (2.4%) | 505 (15.9%) |
| *Hispanic/Latino* | 1,223 (23.0%) | - | - |
| *Chinese* | 642 (12.1%) | 123 (0.3%) | - |
| *South Asian* | - | 772 (1.9%) | - |
| *American Indian /Alaskan Native* | - | - | 4 (0.1%) |
| *Asian / Pacific Islander* | - | - | 3 (0.1%) |
| *Other* | - | 962 (2.3%) | 10 (0.3%) |

[1]MESA: Multi-Ethnic Study of Atherosclerosis; UKB: UK Biobank; CHS: Cardiovascular Health Study.
[2]Metrics are presented as mean (standard deviation) for continuous variables, and n (%) for categorical variables.

**Supplementary Table 2. Comparison of health status between MESA[1] train (N=3,993) and test (N=1,332) sets.**

| | | Train (N=3,993) | | Test (N=1,332) | | | |
|---|---|---|---|---|---|---|---|
| Categorical Variable | | | | | | | |
| **Variables** | **Level** | **N** | **proportion** | **N** | **proportion** | **P values** | **Method** |
| Self-reported Cancer | Don't know | 4 | 0.001 | - | - | 0.571 | Fisher Exact |
| Self-reported Cancer | No | 3665 | 0.918 | 1229 | 0.923 | 0.571 | Fisher Exact |
| Self-reported Cancer | Yes | 319 | 0.080 | 99 | 0.074 | 0.571 | Fisher Exact |
| Self-reported Cancer | NA | 5 | 0.001 | 4 | 0.003 | 0.571 | Fisher Exact |
| Smoking Status | Current | 525 | 0.131 | 176 | 0.132 | 0.784 | Chisq |
| Smoking Status | Former | 1449 | 0.363 | 496 | 0.372 | 0.784 | Chisq |
| Smoking Status | Never | 2007 | 0.503 | 655 | 0.492 | 0.784 | Chisq |
| Smoking Status | NA | 12 | 0.003 | 5 | 0.004 | 0.784 | Chisq |
| Diabetes Mellitus | No | 3478 | 0.871 | 1149 | 0.863 | 0.445 | Chisq |
| Diabetes Mellitus | Yes | 511 | 0.128 | 182 | 0.137 | 0.445 | Chisq |
| Diabetes Mellitus | NA | 4 | 0.001 | 1 | 0.001 | 0.445 | Chisq |
| Highest Education Completed | Associate Degree | 209 | 0.052 | 55 | 0.041 | 0.180 | Chisq |
| Highest Education Completed | Bachelor | 679 | 0.170 | 246 | 0.185 | 0.180 | Chisq |
| Highest Education Completed | Grades 1-8 | 425 | 0.106 | 120 | 0.090 | 0.180 | Chisq |
| Highest Education Completed | Grades 9-11 | 256 | 0.064 | 100 | 0.075 | 0.180 | Chisq |
| Highest Education Completed | Graduate | 677 | 0.170 | 242 | 0.182 | 0.180 | Chisq |
| Highest Education Completed | High School/GED | 739 | 0.185 | 232 | 0.174 | 0.180 | Chisq |

| | | | | | | | |
|---|---|---|---|---|---|---|---|
| Highest Education Completed | No Schooling | 48 | 0.012 | 14 | 0.011 | 0.180 | Chisq |
| Highest Education Completed | Some College/No Degree | 637 | 0.160 | 226 | 0.170 | 0.180 | Chisq |
| Highest Education Completed | Tech School | 310 | 0.078 | 92 | 0.069 | 0.180 | Chisq |
| Highest Education Completed | NA | 13 | 0.003 | 5 | 0.004 | 0.180 | Chisq |
| Gender | Female | 2080 | 0.521 | 683 | 0.513 | 0.629 | Chisq |
| Gender | Male | 1913 | 0.479 | 649 | 0.487 | 0.629 | Chisq |
| Self-reported General Health | Excellent | 636 | 0.159 | 207 | 0.155 | 0.490 | Chisq |
| Self-reported General Health | Fair | 311 | 0.078 | 115 | 0.086 | 0.490 | Chisq |
| Self-reported General Health | Good | 1655 | 0.414 | 541 | 0.406 | 0.490 | Chisq |
| Self-reported General Health | Poor | 21 | 0.005 | 12 | 0.009 | 0.490 | Chisq |
| Self-reported General Health | Very Good | 1343 | 0.336 | 448 | 0.336 | 0.490 | Chisq |
| Self-reported General Health | NA | 27 | 0.007 | 9 | 0.007 | 0.490 | Chisq |
| Self-reported High Cholesterol | Don't know | 13 | 0.003 | 9 | 0.007 | 0.223 | Chisq |
| Self-reported High Cholesterol | No | 2479 | 0.621 | 821 | 0.616 | 0.223 | Chisq |
| Self-reported High Cholesterol | Yes | 1497 | 0.375 | 502 | 0.377 | 0.223 | Chisq |
| Self-reported High Cholesterol | NA | 4 | 0.001 | - | - | 0.223 | Chisq |
| Hypertension by JNC VI (1997) Criteria | No | 2212 | 0.554 | 743 | 0.558 | 0.832 | Chisq |
| Hypertension by JNC VI (1997) Criteria | Yes | 1781 | 0.446 | 589 | 0.442 | 0.832 | Chisq |
| Race | Black | 1003 | 0.251 | 346 | 0.260 | 0.604 | Chisq |
| Race | Chinese | 474 | 0.119 | 168 | 0.126 | 0.604 | Chisq |
| Race | Hispanic/Latino | 933 | 0.234 | 290 | 0.218 | 0.604 | Chisq |
| Race | White | 1583 | 0.396 | 528 | 0.396 | 0.604 | Chisq |

| Continuous Variables | | | | | | | |
|---|---|---|---|---|---|---|---|
| **Variable** | - | **Mean** | **SD** | **Mean** | **SD** | **P-value** | **Methods** |
| Age | - | 62.277 | 10.336 | 61.938 | 10.178 | 0.313 | Wilcoxon |
| BMI | - | 28.290 | 5.413 | 28.513 | 5.604 | 0.402 | Wilcoxon |
| eGFR (CKD-EPI, calibrated creatinine) | - | 77.512 | 16.371 | 78.024 | 16.665 | 0.317 | Wilcoxon |

[1]MESA: Multi-Ethnic Study of Atherosclerosis.

**Supplementary Table 3. Post-imputation Pearson and Spearman correlation of 1,737 proteins in MESA[1] test set (N=1,332).**

| | SomaScan to Olink | | | Olink to SomaScan | | |
|---|---|---|---|---|---|---|
| **Tiers** | **N proteins** | **Median Pearson r** | **Median Spearman rho** | **N proteins** | **Median Pearson r** | **Median Spearman rho** |
| 1 | 334 | 0.941 | 0.949 | 334 | 0.903 | 0.935 |
| 2 | 602 | 0.717 | 0.770 | 613 | 0.693 | 0.746 |
| 3 | 442 | 0.211 | 0.263 | 411 | 0.232 | 0.240 |
| 4 | 359 | 0.039 | 0.047 | 379 | 0.034 | 0.036 |

[1]MESA: Multi-Ethnic Study of Atherosclerosis;

**Supplementary Table 4. Calibration of SomaScan (N=4,972) and Olink (N=2,871) proteins in the MESA[1] test set (N=1,332).**

| | SomaScan to Olink | | | | | | Olink to SomaScan | | | | | |
|---|---|---|---|---|---|---|---|---|---|---|---|---|
| **Tiers** | **N Proteins** | **Median Slope** | **Median Intercept** | **Median RMSE[2]** | **Median SD Ratio** | **Median CCC[3]** | **N Proteins** | **Median Slope** | **Median Intercept** | **Median RMSE[2]** | **Median SD Ratio** | **Median CCC[3]** |
| 1 | 334 | 0.988 | -0.007 | 0.395 | 0.837 | 0.815 | 334 | 1.008 | 0.001 | 0.495 | 0.863 | 0.860 |
| 2 | 602 | 0.963 | -0.008 | 0.437 | 0.675 | 0.601 | 613 | 0.967 | -0.002 | 0.736 | 0.667 | 0.609 |
| 3 | 442 | 0.939 | -0.009 | 0.513 | 0.585 | 0.499 | 411 | 0.746 | -0.016 | 0.876 | 0.507 | 0.297 |
| 4 | 359 | 0.798 | -0.006 | 0.645 | 0.420 | 0.252 | 379 | 0.704 | -0.010 | 0.891 | 0.485 | 0.258 |
| non-overlapping | 1134 | 0.904 | -0.008 | 0.639 | 0.558 | 0.431 | 3235 | 0.804 | -0.013 | 0.855 | 0.528 | 0.346 |
| All | 2871 | 0.945 | -0.008 | 0.552 | 0.621 | 0.534 | 4972 | 0.852 | -0.010 | 0.833 | 0.554 | 0.391 |

[1]MESA: Multi-Ethnic Study of Atherosclerosis;
[2]RMSE: root mean square error. Slope, intercept and RMSE were obtained using lm(measured ~ predicted value).
[3]CCC: Lin's concordance correlation coefficient, measures agreement between imputed and measured values.

**Supplementary Table 5. Uncertainty interval using split conformal prediction for SomaScan (N=4,972) and Olink (N=2,871) proteins in the MESA[1] test set (N=1,332).**

| | SomaScan to Olink | | | | | Olink to SomaScan | | | | |
|---|---|---|---|---|---|---|---|---|---|---|
| **Tiers** | **N proteins** | **Mean Half Width** | **Median Half Width** | **Max Half Width** | **Min Half Width** | **N proteins** | **Mean Half Width** | **Median Half Width** | **Max Half Width** | **Min Half Width** |
| 1 | 334 | 0.790 | 0.653 | 4.075 | 0.356 | 334 | 0.903 | 0.913 | 1.326 | 0.425 |
| 2 | 602 | 0.939 | 0.751 | 4.085 | 0.324 | 613 | 1.308 | 1.311 | 2.126 | 0.606 |
| 3 | 442 | 1.183 | 0.974 | 6.400 | 0.278 | 411 | 1.436 | 1.427 | 2.264 | 0.584 |
| 4 | 359 | 1.585 | 1.206 | 6.344 | 0.261 | 379 | 1.464 | 1.466 | 2.230 | 0.628 |
| non-overlapping | 1134 | 1.508 | 1.208 | 6.806 | 0.328 | 3235 | 1.384 | 1.416 | 3.206 | 0.375 |
| All Proteins | 2871 | 1.265 | 1.014 | 6.806 | 0.261 | 4972 | 1.353 | 1.391 | 3.206 | 0.375 |

[1]MESA: Multi-Ethnic Study of Atherosclerosis;
Uncertainty interval obtained using split conformal prediction.

**Supplementary Table 6. Benchmarking HGB against LightGBM and ElasticNet**

| **Method Pair[1]** | **Direction** | **N Protein (Kappa)[2]** | **κ** |
|---|---|---|---|
| HGB vs LightGBM | SomaScan to Olink | 1737 | 0.975 |
| HGB vs LightGBM | Olink to SomaScan | 1737 | 0.957 |
| HGB vs ElasticNet | SomaScan to Olink | 1726 | 0.875 |
| HGB vs ElasticNet | Olink to SomaScan | 1737 | 0.834 |
| LightGBM vs ElasticNet | SomaScan to Olink | 1726 | 0.881 |
| LightGBM vs ElasticNet | Olink to SomaScan | 1737 | 0.833 |
| **Method** | **Direction** | **N proteins matched With Sissala et al., 2025** | **$\eta^2$** |
| HGB | SomaScan to Olink | 821 | 0.145 |
| HGB | Olink to SomaScan | 821 | 0.175 |
| LightGBM | SomaScan to Olink | 821 | 0.143 |
| LightGBM | Olink to SomaScan | 821 | 0.175 |
| ElasticNet | SomaScan to Olink | 819 | 0.122 |
| ElasticNet | Olink to SomaScan | 821 | 0.172 |
| **Method Pair** | **Direction** | **Z** | **P-value** |
| HGB vs LightGBM | SomaScan to Olink | -0.077 | 0.938 |
| HGB vs LightGBM | Olink to SomaScan | -0.387 | 0.699 |
| HGB vs ElasticNet | SomaScan to Olink | 7.105 | <0.001 |
| HGB vs ElasticNet | Olink to SomaScan | -1.243 | 0.214 |
| LightGBM vs ElasticNet | SomaScan to Olink | 6.993 | 0.000 |
| LightGBM vs ElasticNet | Olink to SomaScan | -1.190 | 0.234 |

[1]HGB: Histogram-based boosting gradient; LightGBM: Light Gradient Boosting Machine.

[2]Proteins for which elastic net selected no predictors (intercept-only models with constant predictions; N=11, SomaScan-to-Olink) were excluded from comparisons involving elastic net, as correlation is undefined for constant predictions.

**Supplementary Table 7. Protein Tier membership thresholds.**

| **Tiers** | **Baseline correlation (r)[1]** | **Imputed correlation (r)** |
|---|---|---|
| Tier 1, ‘most robust’ | ≥0.7 | ≥0.7 |
| Tier 2, ‘model recoverable’ | | ≥0.4 |
| Tier 3, ‘ambivalent’ | | <0.4 |
| Tier 4, ‘irreconcilable’ | <0.1 | <0.1 |

[1]Correlation defined as Pearson correlation. Baseline correlation is defined as r between the measured SomaScan and Olink value. Imputed correlation is defined as the Pearson correlation between SomaScan and SomaScan-imputed Olink (SomaScan to Olink direction), or between Olink and Olink-imputed SomaScan (Olink to SomaScan direction).

**Supplementary Table 8. Alamar validation MAPT, GFAP, and NfL in the Atherosclerosis Risk in Communities Study.**

| | **Alamar - Simoa Correlation[1]** | **Alamar Correlation[1]** | | | **Simoa / Quanterix Correlation[1]** | | |
|---|---|---|---|---|---|---|---|
| **Protein[2]** | (N=14) | **N** | **with SomaScan** | **with Olink** | **N** | **with SomaScan** | **with Olink** |
| **NfL** | 0.86 | 101 | -0.11 | 0.81 | 14 | -0.14 | 0.76 |
| **GFAP** | 0.67 | 101 | 0.14 | 0.94 | 14 | 0.19 | 0.54 |
| **MAPT / pTau 181** | 0.87 | 101 | -0.01 | 0.07 | 14 | 0.2 | -0.11 |

[1]Correlation: Spearman correlation; correlation calculated between Alamar and SomaScan and Olink (N=101).
[2]NfL: neurofilament light polypeptide; GFAP: glial fibrillary acidic protein; MAPT: microtubule-associated protein tau;

**Supplementary Table 9. Demographic characteristics of BMI analytical cohorts.**

| Metrics[1] | UKB[2] (N=41,185) | CHS[2] (N=3,153) |
|---|---|---|
| **BMI** | 27.4 (4.80) | 26.7 (4.54) |
| **Age (years)** | 57.3 (8.19) | 74.4 (4.90) |
| **Sex** | | |
| *Female* | 22,294 (54.1%) | 1,914 (60.7%) |
| *Male* | 18,891 (45.9%) | 1,239 (39.3%) |
| **Race** | | |
| *White* | 38,370 (93.2%) | 2,633 (83.5%) |
| *Black* | 994 (2.4%) | 503 (16.0%) |
| *Hispanic/Latino* | - | - |
| *Chinese* | 123 (0.3%) | - |
| *South Asian* | 756 (1.8%) | - |
| *American Indian /Alaskan Native* | - | 4 (0.1%) |
| *Asian / Pacific Islander* | - | 3 (0.1%) |
| *Other* | 942 (2.3%) | 10 (0.3%) |

[1]Metrics are presented as mean (standard deviation) for continuous variables, and n (%) for categorical variables. BMI: body mass index ($Kg/m^2$), calculated using weight (Kg) / $height^2$ (m).
[2]UKB: UK Biobank; CHS: Cardiovascular Health Study.

**Supplementary Table 10. Application 1 - feature importance[1] of proteins in BMI prediction.**

| CHS SomaScan (Full Panel) | UKB Olink (Full Panel) | CHS SomaScan (Tier 1) | CHS Imputed Olink (Tier 1) | UKB Imputed SomaScan (Tier 1) | UKB Olink (Tier 1) |
|---|---|---|---|---|---|
| LEP (0.356) | LEP (0.082) | LEP (0.549) | LEP (0.462) | LEP (0.293) | LEP (0.288) |
| CYTL1 (0.02) | TGFBR2 (0.017) | FABP4 (0.054) | FABP4 (0.05) | FABP4 (0.054) | FABP4 (0.091) |
| DSG2 (0.016) | FABP4 (0.012) | RNASE1 (0.022) | NCAN (0.019) | NCAN (0.03) | NCAN (0.033) |
| RNASE1 (0.016) | COL15A1 (0.011) | IGFBP1 (0.015) | RNASE1 (0.016) | RNASE1 (0.019) | ART3 (0.013) |
| SEZ6L (0.015) | FSHB (0.01) | GHR (0.012) | GHR (0.014) | ART3 (0.013) | ERP29 (0.008) |

[1]Feature importance calculated with permutation feature importance, where the importance of a protein to the model prediction was measured by the drop in $R^2$ after randomly shuffling that protein's values.

**Supplementary Table 11. Characteristics of the study population in the UK Biobank in replication studies.**

| Application 2 - Walker et al., Nature Aging 2021 | | Application 3 - Shah et al., Nature Communications 2024 | |
|---|---|---|---|
| **Metrics(Dementia)[2]** | **Dementia (N=36,854)** | **Metrics (Heart Failure)** | **Heart Failure (N=41,071)** |
| **Age (years)** | 57.3 (8.16) | **Age (years)** | 57.4 (8.18) |
| **Sex** | | **Sex** | |
| *Female* | 19,020 (53.9%) | *Female* | 21,229 (53.9%) |
| *Male* | 16,283 (46.1%) | *Male* | 18,131 (46.1%) |
| **White British** | 29,308 (83.0%) | **White British** | 32,576 (82.8%) |
| **BMI** | 27.4 (4.72) | **BMI** | 27.4 (4.73) |
| **Ever Smoker** | 16,158 (45.8%) | **Ever Smoker** | 17,953 (45.6%) |
| **eGFR-Creatinine[2]** | 93.8 (14.0) | **eGFR-Creatinine[2]** | 93.9 (13.6) |
| **Baseline Hypertension** | 11,653 (33.0%) | **Baseline Hypertension** | 13,061 (33.2%) |
| **Prevalent Diabetes** | 2,283 (6.5%) | **Prevalent Diabetes** | 2,571 (6.5%) |
| **APOE-ε4[3]** | 10,289 (29.1%) | **Prevalent CAD[5]** | 2,572 (6.5%) |
| **Education[4]** | | **Prevalent AF[5]** | 889 (2.3%) |
| *≤High School* | 15,769 (44.7%) | | |
| *Vocational School* | 2,313 (6.6%) | | |
| *General Education* | 3,889 (11.0%) | | |
| *Graduate or Professional School* | 13,332 (37.8%) | | |

[1]Dementia: Dementia, Walker et al., Nature Aging 2021; Heart Failure: Shah et al., Nature Communications 2024.
[2]eGFR-Creatinine: estimated glomerular filtration rate, calculated using the 2021 CKD-EPI Creatinine equation.
[3]APOE-ε4: Apolipoprotein E ε4 status, presence of at least one ε4 allele.

[4]Education: Less than and high school: O levels/GCSEs or equivalent, CSEs or equivalent, and ‘none of the above’ option. Vocational School: NVQ, HND, HNC or equivalent. General Education: A levels/AS levels or equivalent. Graduate or Professional School: College or University degree, Other professional qualifications, e.g., nursing, teaching.
[5]CAD: Coronary Artery Disease; AF: Atrial fibrillation or flutter.
Metrics are presented as mean (standard deviation) for continuous variables, and n (%) for categorical variables. The study population was restricted to unrelated individuals in the UK Biobank with available whole-exome sequencing for the Dementia replication study, with unrelatedness defined as less than third-degree relatedness. No such exclusion for the HF study, as it didn’t utilize genetic information. For individual analysis on dementia and HF, the study populations were further restricted to individuals free of the outcome disease at baseline.

**Supplementary Table 12. Application 2 - Replication Results of Walker et al., Nature Aging 2021.**

| | Walker et al., Nature Aging 2021 (ARIC midlife) | | UKB Predicted SomaScan | | Walker et al., Nature Aging 2021 (ARIC latelife) | | Walker et al., Nature Aging 2021 (AGES-Reykjavik) | |
|---|---|---|---|---|---|---|---|---|
| **Protein** | **HR (95% CI)** | **P-value** | **HR (95% CI)** | **P-value** | **HR (95% CI)** | **P-value** | **HR (95% CI)** | **P-value** |
| SVEP1 | 1.42 (1.20-1.67) | 3.05E-05 | 1.47 (1.17-1.83) | 7.40E-04 | 1.93 (1.61-2.31) | 7.03E-13 | 1.42 (1.25-1.61) | 8.17E-08 |
| SVEP1_2 | 1.49 (1.28-1.74) | 3.43E-07 | 1.33 (1.07-1.66) | 1.07E-02 | 1.93 (1.59-2.30) | 3.94E-12 | 1.42 (1.25-1.61) | 2.82E-08 |
| CPLX2 | 1.52 (1.31-1.77) | 3.03E-08 | 1.03 (0.82-1.28) | 8.24E-01 | 1.82 (1.47-2.25) | 4.08E-08 | (not measured) | |
| TAGLN | 1.54 (1.30-1.82) | 7.36E-07 | 1.13 (0.90-1.42) | 3.06E-01 | 2.10 (1.60-2.74) | 5.84E-08 | 1.68 (1.42-2.00) | 3.85E-09 |
| FBLN5 | 2.32 (1.85-2.92) | 4.24E-13 | 1.07 (0.85-1.34) | 5.74E-01 | 2.46 (1.70-3.56) | 1.84E-06 | 1.27 (1.12-1.44) | 2.00E-04 |
| BAGE2 | 1.43 (1.24-1.65) | 1.11E-06 | 0.94 (0.78-1.13) | 4.97E-01 | 1.73 (1.37-2.19) | 5.32E-06 | 1.15 (0.96-1.37) | 1.40E-01 |

[1]HR: hazard ratio.

Proteins are ordered according to the original study (Walker et al., Nature Aging 2021)'s significance, with SVEP1 being most significant findings that replicated across cohorts in observational studies and in causal inferencing using two-sample Mendelian randomization.

**Supplementary Table 13. Replication Results of Application 3 - Shah et al., Nature Communications 2024.**

| | Shah et al., Nature Communications 2024 | | UKB Imputed SomaScan | | UKB Olink | |
|---|---|---|---|---|---|---|
| **Protein** | **HR (95% CI)** | **P-value** | **HR (95% CI)** | **P-value** | **HR (95% CI)** | **P-value** |
| GDF15 | 1.51 (1.41-1.62) | 2.73E-33 | 2.12 (1.98-2.27) | 6.66E-102 | 2.25 (2.09-2.43) | 1.94E-99 |
| NPPB | 1.46 (1.37-1.55) | 1.75E-32 | 1.40 (1.30-1.52) | 9.58E-18 | 1.36 (1.31-1.41) | 3.32E-67 |
| WFDC2 | 1.45 (1.35-1.55) | 1.69E-27 | 2.23 (2.08-2.39) | 3.52E-112 | 2.99 (2.73-3.28) | 4.18E-120 |
| SVEP1_2 | 1.35 (1.28-1.43) | 2.23E-25 | 2.64 (2.44-2.86) | 3.86E-124 | - | - |
| SVEP1 | 1.34 (1.27-1.41) | 2.58E-25 | 2.71 (2.50-2.93) | 1.28E-135 | - | - |
| MMP12 | 1.35 (1.26-1.44) | 4.45E-19 | 1.50 (1.42-1.59) | 1.94E-44 | 1.65 (1.53-1.78) | 3.68E-41 |
| CLEC3B | 0.75 (0.70-0.80) | 7.12E-18 | 0.62 (0.56-0.68) | 1.13E-23 | 0.40 (0.31-0.50) | 3.60E-15 |
| ANGPT2 | 1.33 (1.24-1.42) | 4.08E-16 | 1.85 (1.74-1.97) | 1.95E-84 | 2.39 (2.16-2.65) | 1.72E-61 |
| FBLN5 | 1.23 (1.17-1.30) | 1.96E-15 | 2.44 (2.24-2.65) | 2.19E-98 | - | - |
| TNNT2 | 1.21 (1.15-1.28) | 6.49E-14 | 1.57 (1.47-1.68) | 2.57E-39 | - | - |
| MSR1 | 1.30 (1.21-1.39) | 4.81E-13 | 1.70 (1.48-1.96) | 1.73E-13 | 1.70 (1.55-1.87) | 6.23E-29 |
| CACNA2D3 | 0.76 (0.70-0.82) | 1.13E-12 | 0.52 (0.47-0.58) | 1.23E-32 | - | - |
| TAGLN | 1.26 (1.17-1.34) | 3.59E-11 | 2.09 (1.90-2.29) | 8.30E-54 | - | - |
| IGDCC4 | 0.82 (0.77-0.87) | 5.09E-11 | 0.67 (0.61-0.73) | 1.20E-18 | 0.51 (0.43-0.60) | 7.15E-15 |
| CLIC2 | 1.17 (1.10-1.24) | 2.43E-07 | 1.25 (1.17-1.34) | 9.57E-11 | - | - |

| | | | | | | |
|---|---|---|---|---|---|---|
| NPPB_2 | 1.15 (1.09-1.21) | 5.29E-07 | 2.12 (1.98-2.28) | 1.78E-103 | - | - |
| SLITRK1 | 0.96 (0.89-1.04) | 3.36E-01 | 0.90 (0.82-1.00) | 4.96E-02 | 0.66 (0.57-0.76) | 2.59E-08 |
| PTPRD | 0.90 (0.84-0.97) | 6.35E-03 | 0.73 (0.67-0.80) | 3.65E-11 | - | - |

[1]HR: hazard ratio.

**Supplementary Table 14. Summary of SHAP-based feature importance in the overlapping proteins (N=1,737) in MESA test set (N=1,332)**

| Direction | Tiers | N protein | % Top1 is self | Top-5 predictor share (median)[1] | Top-10 predictor share (median) | Top-20 predictor share (median) |
|---|---|---|---|---|---|---|
| SomaScan to Olink | 1 | 334 | 82.63 | 0.303 | 0.327 | 0.360 |
| | 2 | 602 | 65.45 | 0.186 | 0.226 | 0.271 |
| | 3 | 442 | 7.47 | 0.127 | 0.177 | 0.233 |
| | 4 | 359 | 1.11 | 0.077 | 0.110 | 0.157 |
| Olink to SomaScan | 1 | 334 | 81.44 | 0.308 | 0.340 | 0.391 |
| | 2 | 613 | 65.58 | 0.169 | 0.212 | 0.261 |
| | 3 | 411 | 10.71 | 0.075 | 0.108 | 0.156 |
| | 4 | 379 | 0.53 | 0.067 | 0.098 | 0.143 |

[1]Top-X feature: the percentage of feature importance of the top X features / total (SHAP importance of all the predictors).

**Supplementary Table 15. Direction- and pairwise- comparison for all Alphafold 3 metrics.**

| Metric | Direction | Comparison | P-value[1] | Bonferroni adjusted P-value | Significant?[2] |
|---|---|---|---|---|---|
| | | Tier 1 vs Tier 2 | 4.04E-01 | 1.00E+00 | ns |
| | | Tier 2 vs Tier 3 | 1.41E-03 | 8.48E-03 | ** |
| | | Tier 3 vs Tier 4 | 6.81E-02 | 4.09E-01 | ns |
| | | Tier 1 vs Tier 3 | 5.02E-02 | 3.01E-01 | ns |
| | | Tier 2 vs Tier 4 | 2.81E-07 | 1.69E-06 | *** |
| | SomaScan to Olink | Tier 1 vs Tier 4 | 2.87E-04 | 1.72E-03 | ** |
| | | Tier 1 vs Tier 2 | 6.83E-01 | 1.00E+00 | ns |
| | | Tier 2 vs Tier 3 | 3.61E-03 | 2.17E-02 | * |
| | | Tier 3 vs Tier 4 | 4.18E-01 | 1.00E+00 | ns |
| | | Tier 1 vs Tier 3 | 3.38E-02 | 2.03E-01 | ns |
| | | Tier 2 vs Tier 4 | 1.24E-04 | 7.42E-04 | *** |
| Fraction Disordered | Olink to SomaScan | Tier 1 vs Tier 4 | 4.49E-03 | 2.69E-02 | * |
| | | Tier 1 vs Tier 2 | 6.16E-01 | 1.00E+00 | ns |
| | | Tier 2 vs Tier 3 | 2.62E-02 | 1.57E-01 | ns |
| | | Tier 3 vs Tier 4 | 5.16E-03 | 3.10E-02 | * |
| | | Tier 1 vs Tier 3 | 2.19E-02 | 1.32E-01 | ns |
| | | Tier 2 vs Tier 4 | 2.05E-07 | 1.23E-06 | *** |
| | SomaScan to Olink | Tier 1 vs Tier 4 | 1.54E-06 | 9.21E-06 | *** |
| | | Tier 1 vs Tier 2 | 3.56E-01 | 1.00E+00 | ns |
| | | Tier 2 vs Tier 3 | 5.07E-02 | 3.04E-01 | ns |
| | | Tier 3 vs Tier 4 | 7.75E-02 | 4.65E-01 | ns |
| Average pLDDT | Olink to SomaScan | Tier 1 vs Tier 3 | 1.85E-02 | 1.11E-01 | ns |

| | | | | | |
|---|---|---|---|---|---|
| | | Tier 2 vs Tier 4 | 8.18E-05 | 4.91E-04 | *** |
| | | Tier 1 vs Tier 4 | 4.44E-05 | 2.66E-04 | *** |
| | | Tier 1 vs Tier 2 | 7.71E-01 | 1.00E+00 | ns |
| | | Tier 2 vs Tier 3 | 6.93E-02 | 4.16E-01 | ns |
| | | Tier 3 vs Tier 4 | 2.13E-01 | 1.00E+00 | ns |
| | | Tier 1 vs Tier 3 | 2.01E-01 | 1.00E+00 | ns |
| | | Tier 2 vs Tier 4 | 1.88E-03 | 1.13E-02 | * |
| | SomaScan to Olink | Tier 1 vs Tier 4 | 1.79E-02 | 1.08E-01 | ns |
| | | Tier 1 vs Tier 2 | 9.56E-01 | 1.00E+00 | ns |
| | | Tier 2 vs Tier 3 | 6.19E-02 | 3.71E-01 | ns |
| | | Tier 3 vs Tier 4 | 8.87E-01 | 1.00E+00 | ns |
| | | Tier 1 vs Tier 3 | 1.02E-01 | 6.12E-01 | ns |
| | | Tier 2 vs Tier 4 | 9.26E-02 | 5.55E-01 | ns |
| pTM | Olink to SomaScan | Tier 1 vs Tier 4 | 1.37E-01 | 8.21E-01 | ns |
| | | Tier 1 vs Tier 2 | 4.85E-01 | 1.00E+00 | ns |
| | | Tier 2 vs Tier 3 | 7.93E-01 | 1.00E+00 | ns |
| | | Tier 3 vs Tier 4 | 5.99E-01 | 1.00E+00 | ns |
| | | Tier 1 vs Tier 3 | 3.62E-01 | 1.00E+00 | ns |
| | | Tier 2 vs Tier 4 | 3.58E-01 | 1.00E+00 | ns |
| | SomaScan to Olink | Tier 1 vs Tier 4 | 1.26E-01 | 7.54E-01 | ns |
| | | Tier 1 vs Tier 2 | 3.52E-01 | 1.00E+00 | ns |
| | | Tier 2 vs Tier 3 | 6.50E-01 | 1.00E+00 | ns |
| Ranking Score | Olink to SomaScan | Tier 3 vs Tier 4 | 4.60E-01 | 1.00E+00 | ns |

| | | | | | |
|---|---|---|---|---|---|
| | | Tier 1 vs Tier 3 | 2.23E-01 | 1.00E+00 | ns |
| | | Tier 2 vs Tier 4 | 8.68E-01 | 1.00E+00 | ns |
| | | Tier 1 vs Tier 4 | 3.93E-01 | 1.00E+00 | ns |

[1]The pairwise comparison p-value is obtained using Wilcoxon rank-sum test, and further adjusted for multiple testing using Bonferroni.

[2] *: adjusted P<0.05; **: adjusted P<0.01; ***: adjusted P<0.001; ns: not significant.
